# MRS-Sim: Open-Source Framework for Simulating In Vivo-like Magnetic Resonance Spectra

**DOI:** 10.1101/2024.12.20.629645

**Authors:** John LaMaster, Georg Oeltzschner, Yan Li

**Affiliations:** Munich Institute of Biomedical Engineering, Technical University of Munich, Bavaria, Germany; School of Computation, Information, and Technology, Technical University of Munich, Bavaria, Germany; Russel H. Morgan Department of Radiology and Radiological Sciences, The Johns Hopkins’ University School of Medicine, Maryland, USA; Department of Radiology and Biomedical Imaging, University of California, San Francisco, California, USA

**Keywords:** MRS, spectroscopy, in vivo, data simulation, synthetic data, open-source

## Abstract

Realistic, in vivo-like synthetic data is increasingly needed to develop and validate methods in magnetic resonance spectroscopy. MRS-Sim is a powerful, open-source framework for simulating such data while providing known ground truth values. Its modularity enables modeling the complexities of MRS data for various in vivo scenarios. The underlying physical equations include both commonly used spectral components of linear-combination fitting routines and two novel components. The first is a 3D *B*_0_ field map simulator that models *B*_0_ field inhomogeneities, ranging from slight variations to severe distortions. The second is a novel semi-parametric generator that mimics signals from poorly characterized residual water regions and spectral baseline contributions. This framework can simulate scenarios ranging from raw multi-coil transients to preprocessed, coil-combined multi-average data.

Simulating realistic in vivo-like datasets requires appropriate model parameter ranges and distributions, best determined by analyzing the fitting parameters from existing in vivo data. Therefore, MRS-Sim includes tools for analyzing the ranges and statistical distributions of those parameters from in vivo datasets fitted with Osprey, allowing simulations to be tailored to specific datasets. Additionally, the accompanying repository of supplemental information assists non-expert users with general simulations of MRS data.

The modularity of this framework facilitates easy customization various in vivo scenarios and promotes continued community development. Using a single framework for diverse applications addresses the inconsistencies in current protocols. By simulating in vivo-like data, MRS-Sim supports many MRS tasks, including verifying spectral fitting protocols and conducting reproducibility analyses. Readily available synthetic data also benefits deep learning research, particularly when sufficient in vivo data is unavailable for training. Overall, MRS-Sim will promote reproducibility and make MRS research more accessible to a wider audience.

## 1 INTRODUCTION

Magnetic resonance spectroscopy (MRS) is a non-invasive MR modality that provides in vivo metabolic profiling of tissues. It facilitates the evaluation of various pathologies by fitting the spectra to estimate metabolite concentrations. In the brain, MRS has been extensively used to investigate a wide range of pathologies, including neurodevelopmental ^1,2,3^ and neurodegenerative diseases ^4,5,6,7^, inborn errors of metabolism ^8,9,10^, and brain tumors ^11,12,13,14^, as well as age-related changes ^15,16,17^.

After data acquisition, pre-processing and spectral fitting are required before metabolic information can be analyzed. Every step in this pipeline must be calibrated to ensure accurate and reliable spectral fitting. However, achieving this is challenging due to variations in pulse sequences, acquisition parameters, data quality regimes, and biological differences (adult versus pediatric subjects ^18^ and disease related metabolic patterns ^19^). Moreover, such large and comprehensive collections of in vivo data are generally unavailable for method development and evaluation often due to high costs and data privacy restrictions. Furthermore, ground truth values are not available for in vivo data, complicating the evaluation of the accuracy of MRS protocols. While phantom data can help in developing and validating acquisition and analysis methods, it often fails to adequately reflect the complexities of in vivo spectra. Synthetic data is the only option that can address these limitations because it can be simulated for diverse scenarios and always has known ground truth values. However, generating it is still challenging because it requires adequately incorporating all physical phenomena underlying in vivo data. Critically, one must choose adequate model parameters that reflect different physiological and pathological conditions. But arguably more important is the need to replicate nuisance signals and acquisition-induced artifacts commonly found in in vivo data that are difficult to reproduce in a phantom, such as effects of susceptibility and field inhomogeneity along with contributions from macromolecules ^20^, lipids ^21^, and residual water ^21^. Many research groups now routinely use synthetic data, but the data generation models and parameter distributions are rarely made publicly available. This lack of transparency hampers systematic comparison and the formation of consensus best practices for synthetic data generation.

### 1.1 Background

The use of synthetic MRS data has increased significantly in recent years, largely driven by the rise of machine learning ^22^ (ML) and deep learning ^23,24,25^ (DL) methods, which require large datasets for training and validation. Since few centers worldwide possess sufficient in vivo data, generating realistic synthetic datasets has become essential. Synthetic data provides known ground truth values that are essential to evaluate the accuracy and precision of the analysis methods being developed. However, growing reliance on synthetic data in ML and DL applications may exacerbate reproducibility and generalizability challenges already present in traditional MRS research ^26,27,28,29,30^. Addresses these issues requires establishing standards and best practices for synthetic MRS data simulation, which will require researchers to collaborate and build consensus, as well as develop and adopt open-source frameworks for data generation.

There is no consensus in literature on best practices for simulating synthetic data. Existing works use physical models with varying complexity and spectral components ^24,31,32,33^, typically starting with metabolite basis functions simulated with pulse sequence parameters for their specific scenario. These metabolite basis functions are then modulated by scaling factors representing their underlying concentrations. A simple Lorentzian lineshape is generally applied ^24,31,32^ with optional phase offsets ^32,33^ and frequency shifts ^24^. Finally, some form of broad baseline is typically added. These signal models are simple and do not capture the full complexity of in vivo data.

There are several software packages that simulate in vivo spectra, including FID-A ^34^, GAVA ^35^, NMRScope-B ^36^, and VESPA ^37^. The fundamental difference between these methods and the current work lies in the definition of “in vivo spectra”. Until now, this term has referred to simulating basis functions and combining them into idealized spectra to evaluate parameter settings for developing new in vivo pulse sequences. In this work, “in vivo spectra” refers to the data directly acquired from an in vivo scanner. This new definition aims to reframe the task of simulating synthetic data, to challenge what are assumed to be acceptable simplifications, and to advance the use of synthetic data for MRS applications. The most relevant previous work is the recent extension of MARSS ^38^ which has added a new functionality called synMARSS ^39^ that simulates synthetic spectra and incorporates many spectral artifacts. They provide an extensive user manual with equations describing their implementation, however, their work is closed-source.

### 1.2 Contribution

This work presents MRS-Sim: a modular framework for synthetic data generation to address some of these challenges. The simulation model uses well-defined physical parameters commonly used in linear-combination modeling software, corresponding to amplitudes, lineshapes, phases, and frequency shifts. Additionally, it features a realistic *B*_0_ map generator to simulate in vivo-like field inhomogeneities and uses semi-parameterized models for residual water and smooth, non-macromolecule background (non-MMBG) signal contributions.

MRS-Sim can simulate spectra at any stage of acquisition from individual coil elements to unaligned transients to fully processed spectra. This flexibility allows researchers to compare new data processing methods beyond spectral modeling. The code base is highly flexible and can simulate custom-tailored datasets for a large variety of in vivo scenarios. This framework includes a detailed compilation of simulation parameters as well as tools to analyze existing in vivo datasets and extract parameter distributions, facilitating the augmentation of in vivo data with synthetic data. By being open-source, MRS-Sim enables seamless incorporation of future additions from the community, expanding its capabilities. This work is intended to be a community resource to provide researchers, trainees and experts alike, with access to high-quality and comparable synthetic data as a source of continuity in the field.

## 2. METHODS

### 2.1 Terminology

To clarify potentially ambiguous terms, a list of terms and corresponding definitions is presented in the Supporting Information Section 1 Terminology. Readers are strongly encouraged to briefly review those definitions before proceeding.

### 2.2 Physics Model Algorithm

The proposed physics model, shown in Eqns. 1a, 1b, and 1c, was designed to mirror the actual data acquisition sequence by reverse engineering the spectral fitting process. It was developed after a thorough review of current state-of-the-art fitting algorithms (LCModel ^40^, Osprey ^41^, FSL-MRS ^42^, etc.). Commonly modeled spectral features and components were identified for inclusion, allowing for a large variety of scenarios and artifacts. The model was parameterized as follows:

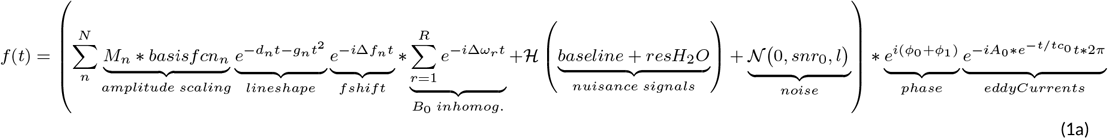

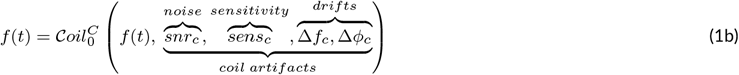

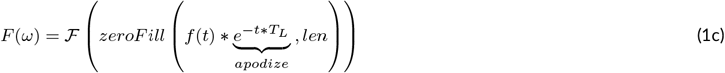

where,

1. *N* is the whole set of metabolites being modeled
2. *M*_*n*_ is the scaling factor for metabolite *n*
3. *d*_*n*_ is the Lorentzian lineshape variable
4. *g*_*n*_ is the Gaussian lineshape variable
5. *t* is the time vector
6. Δ*f*_*n*_ is the metabolite-specific frequency shift
7. *ϕ*_0_ is the global zero-order phase offset
8. *ϕ*_1_ is the global first-order phase offsets
9. *A*_0_ is the amplitude of the eddy current
10. *tc*_0_ is the time constant of the eddy current
11. Δ*ω*_*r*_ are the *B*_0_ inhomogeneities
12. *snr*_0_ is the desired SNR of the spectrum
13. *baseline* is the semi-parameterized, broad baseline offset
14. *resH*_2_*O* is the semi-parameterized residual water contribution
15. ℋ is the transform to create the imaginary components
16. *Coil* generates *C* spectral transients, e.g. multi-coil
17. *snr*_*c*_ scales the transient SNR values
18. *sens*_*c*_ simulates coil sensitivity values
19. Δ*f*_*c*_ are the frequency drifts of the transients
20. Δ*ϕ*_*c*_ are the phase drifts of the transients
21. *T*_*L*_ is the apodization frequency
22. *zeroFill* is the zero-padding operator
23. *len* is the zero-filled target length
24. ℱ is the Fourier transform
25. *f* (*t*) is the FID
26. *F* (*ω*) is the spectrum

Before starting the simulations, an appropriate basis set must be simulated. For each component in the basis set, i.e. metabolite, macromolecule, or lipid, the basis function is scaled and then the lineshape distortion and *B*_0_ inhomogeneities are applied. Moiety-level frequency shifts are applied and then the basis functions are summed into a single FID. Next, the noise, baseline offset, and residual water signal are added. Then the phase offsets and eddy currents are applied. If spectral transients are being simulated, then the noise is not applied until the next stage. In Eqn. 1c, the *Coil* operator generates *C* transients and applies a distribution of scaling factors for SNR values and coil weights. The transients are then scaled and corresponding noise vectors are sampled. Then, phase and frequency drifts are applied to each transient. At this point, the in vivo FIDs are complete. However, if pre-processing steps are also desirable, then apodization and zero-padding can be performed. Then the Fourier transform ℱ can convert the FIDs to the frequency domain. The following sections are presented in order according to their implementation in the physics model and discuss the terms above in greater detail. A step-by-step visualization of this model is illustrated in Fig. 1.

**FIGURE 1.**
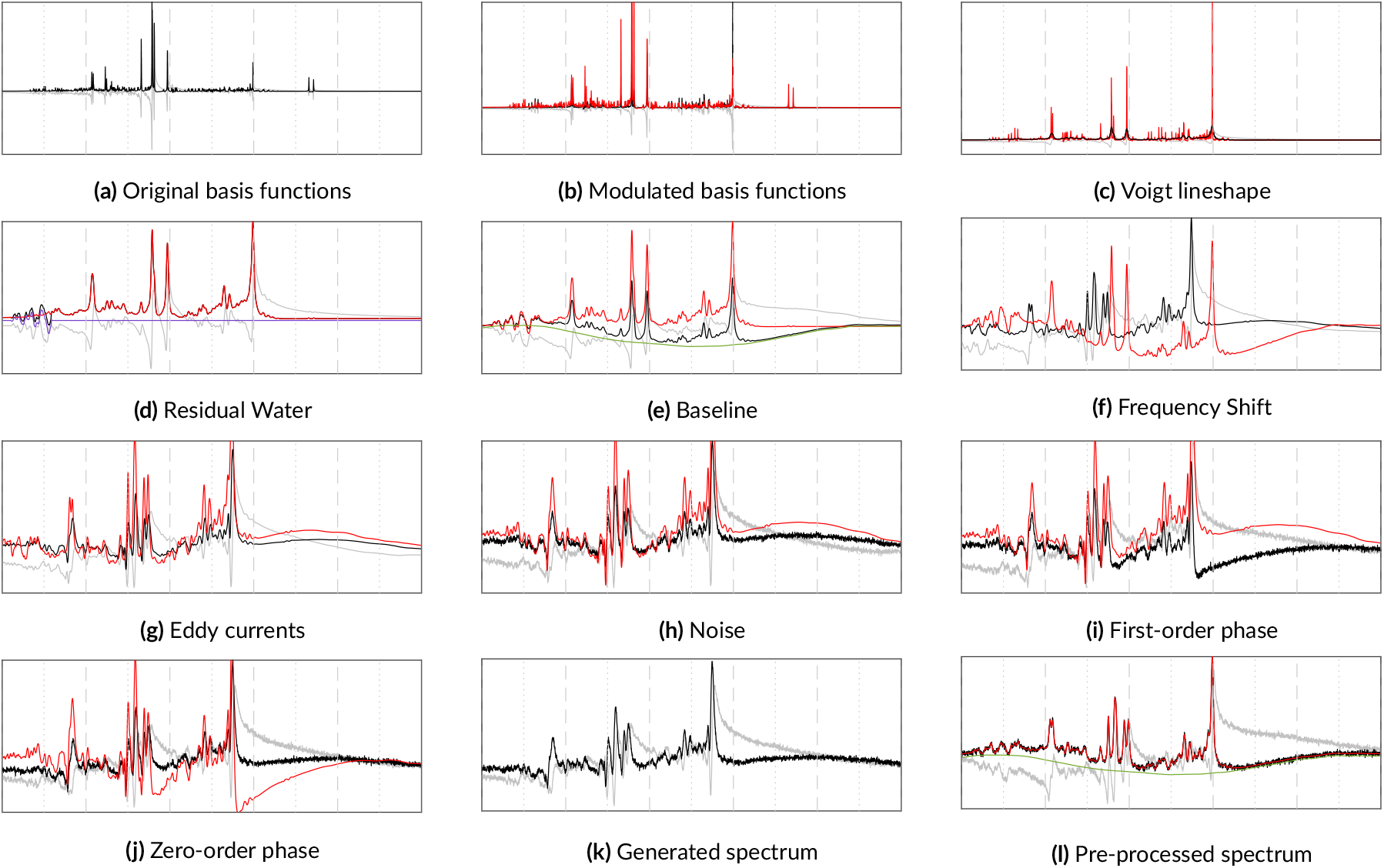
A step-by-step progression through the physics model for simulating a 3T PRESS spectrum. The real and imaginary components are depicted in black and gray, respectively. The red line includes only the metabolites and the offsets from the preceding steps. The purple line is the residual water and the green line is the spectral baseline. 1k shows the final spectrum with all artifacts applied. 1l is the pre-processed spectrum with the phase and frequency shifts removed.

The selection of simulations presented in the rest of this manuscript focus on short echo (TE=30ms) 3T PRESS spectra with a spectral width of 2000Hz and randomly sampled parameters. The choice of short echo spectra is motivated by their ability to capture a broader range of metabolite peaks compared to long echo spectra. The simulations presented include metabolites selected from a comprehensive set of common brain metabolites including ascorbate (Asc), aspartate (Asp), choline (Ch), creatine (Cr), *γ*-aminobutyric acid (GABA), glutamine (Gln), glucose (Glu), glycerylphosphocholine (GPC), glutathione (GSH), lactate (Lac), myo-inositol (mI), N-ascetylaspartate (NAA), N-acetylaspartylglutamate (NAAG), phosphorylcholine (PCh), phosphocreatine (PCr), phosphoethanolamine (PE), scyllo-inositol (sI), taurine (Tau), and a variety of macromolecular and lipid basis functions. For consistency, the SNR was fixed to 15 and the chemical shift reference point was set to 4.65ppm. The metabolite concentrations and 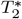 values, adapted from Wyss *et al*. ^43^, were fixed to further emphasize the effects of the simulated artifacts. Any deviations from these settings are noted in the respective figure captions. Not every figure depicts spectra with acceptable quality, the definition of which changes for different scenarios, e.g. comparing SVS with MRSI or diffusion MRS. Each use case will have different requirements for spectral components and artifacts. This framework facilitates simulating them, but the appropriate numerical ranges are the responsibility of the user.

#### 2.2.1 Metabolite Basis Functions

Landheer *et al*.’s Magnetic Resonance Spectrum Simulator (MARSS) ^38^ software package was used for simulating the default metabolite basis functions provided with this simulator, which represent GE 3T PRESS TE=30ms. MRI, and its derivatives, are spatially resolved modalities, meaning that MR pixels are actually voxels that represent 3D volumes. These volumes have a spatial distribution, as shown in Fig. 2a, that needs to be accounted for in simulations. MARSS produces high-fidelity outputs by simulating 128 points in each direction within a voxel, accurately capturing the spatial nature of the RF pulses and slice-selective gradients. Using vendor-specific pulse sequences, MARSS can simulate individual or summed spins for a large number of common brain metabolites, including parameterized macromolecule and lipid resonances.

**FIGURE 2.**
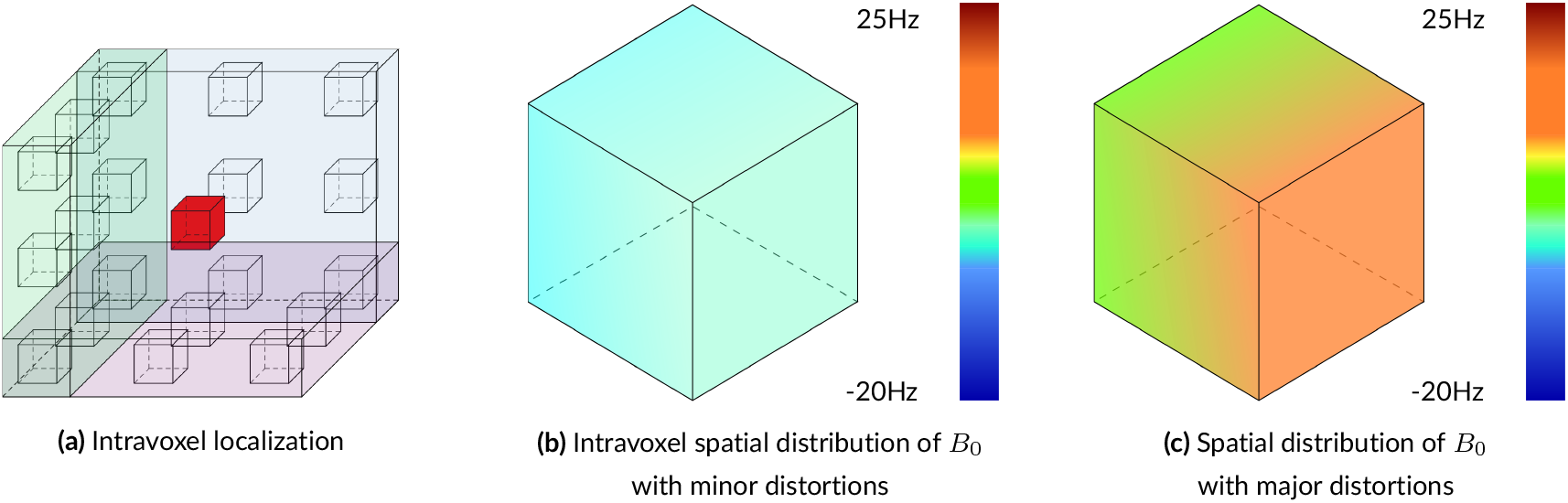
Illustration of intra-voxel spatial localizations. Voxels are often treated as singularities, meaning the entire volume is considered to be a single point like the red box in the center of 2a, which highlights the spatial, 3D nature of voxels. 2b and 2c show normal and extreme levels of intravoxel *B*_0_ inhomogeneity, respectively. Accounting for these intravoxel distributions leads to more accurate and more realistic simulations.

**FIGURE 3.**
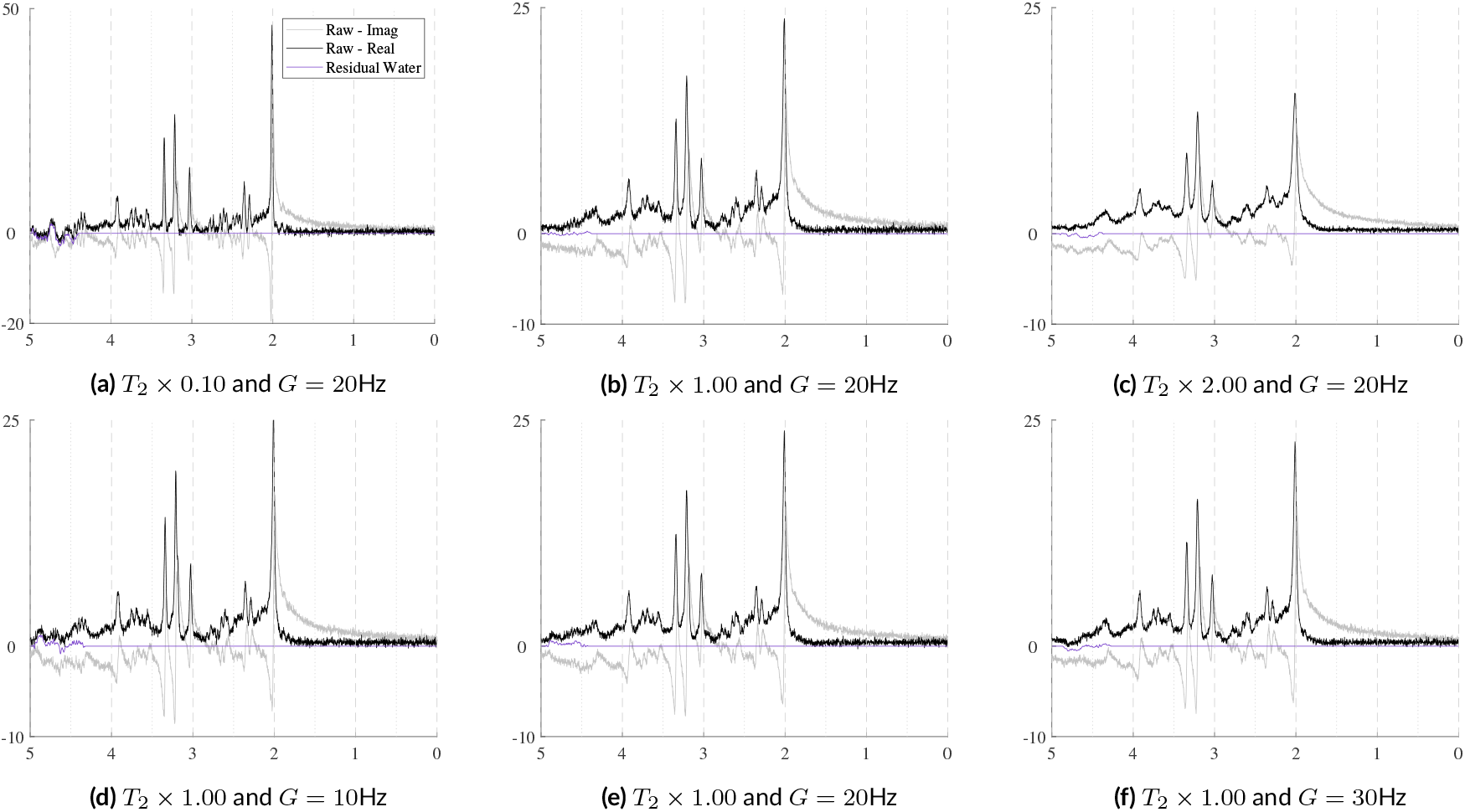
Illustrations of samples exploring the effects of Lorentzian (top) and Gaussian (bottom) broadening on the Voigt lineshape. The spectral SNR = 15. No phase offsets, eddy currents, macromolecules, or spectral baselines were included. The top row scales the Lorentzian broadening while maintaining a fixed Gaussian value whereas the bottom row fixes the Lorentzian broadening values and varies the Gaussian values. The effects on signal intensity should be noted by comparing the apparent peak heights with the scale of the y-axis.

#### 2.2.2 Amplitude Modulation

The amplitude scaling factors *M*_*n*_ reflect the metabolites’ concentrations. For this framework, comprehensive metabolite ranges including physiological and pathological values were adapted from the meta-analysis by Gudmundson *et al*. ^44^ and are presented in the Supporting Information Section 2. To convert metabolite concentrations to amplitudes, this work recommends assuming that the signal in each basis function represents the signal from 1.0mM of the given metabolite. This assumption allows for using the metabolite concentrations directly. Should relative concentrations be desired, this can be easily calculated afterwards because the default ranges are provided in units of millimolar (mM), which allows any metabolite to be used as a reference. This flexibility is important because as Near ^45^ and Larsen ^46^ discussed, the choice of reference metabolite plays a non-trivial role with respect to quantification and interpretability.

#### 2.2.3 Lineshape Profiles

Spectral lineshapes are primarily influenced by metabolite- and moiety-specific relaxation values, magnetic field homogeneity, and the tissue microenvironment. These effects are most commonly modeled in spectral fitting routines with the Voigt lineshape profile, as it most closely matches in vivo data. ^47^ This lineshape is a combination of a Lorentzian and a Gaussian and is used in fitting packages such as LCModel ^40^, TARQUIN ^48^, jMRUI’s ^49^ NMRScope-B ^36^, and Osprey ^41^. For completeness, it is also possible to specify either a purely Lorentzian or a purely Gaussian lineshape. At the most basic level, each metabolite is assigned a Lorentzian value while a single Gaussian value is applied to the set of metabolites and a second value can be applied to the macromolecules and lipids. As is standard practice from the aforementioned software packages, MRS-Sim assigns a single Lorentzian value to each metabolite. A detailed table is provided in the Supporting Information Section 2 with the current information for 3T data; however, an up-to-date digital version will be maintained in the repository. The metabolite-level relaxation values were primarily derived from the meta-analysis from Gudmundson *et al*. ^44^. Future work will allow for specifying moiety-level *T*_2_ relaxation values. Information for selected moieties at 3T was compiled from Wyss *et al*. ^43^. As more metabolites are characterized, new information can be added to the model.

#### 2.2.4 B0 Inhomogeneities

In vivo acquisitions never have perfectly homogeneous magnetic fields. Small *B*_0_ inhomogeneities are, in general, sufficiently modeled by the Gaussian term of the Voigt lineshape. However, to simulate more severe distortions from large inhomogeneities or to model the gaussian broadening, signal damping, and frequency shift with a unified model, a *B*_0_ field volume needs to be modeled and applied to the basis functions. Fig. 2 illustrates the difference between small and severe *B*_0_ inhomogeneities. Fig. 2b shows the normal, subtle *B*_0_ changes across the volume of a spectroscopy voxel while Fig. 2c illustrates high field inhomogeneity. In such cases, spectra from these regions exhibit significant lineshape distortions that cannot be adequately characterized using idealized lineshape profiles. The top row in Fig. 4 illustrates the spectral effects of increasing intra-voxel field inhomogeneity, i.e. increasing Δ, given a fixed field offset, highlighting the impact of the field heterogeneity on lineshape and signal intensity. The bottom row uses a fixed inhomogeneity and modulates the mean *B*_0_ offset, which maintains the same lineshape but induces a global frequency shift.

**FIGURE 4.**
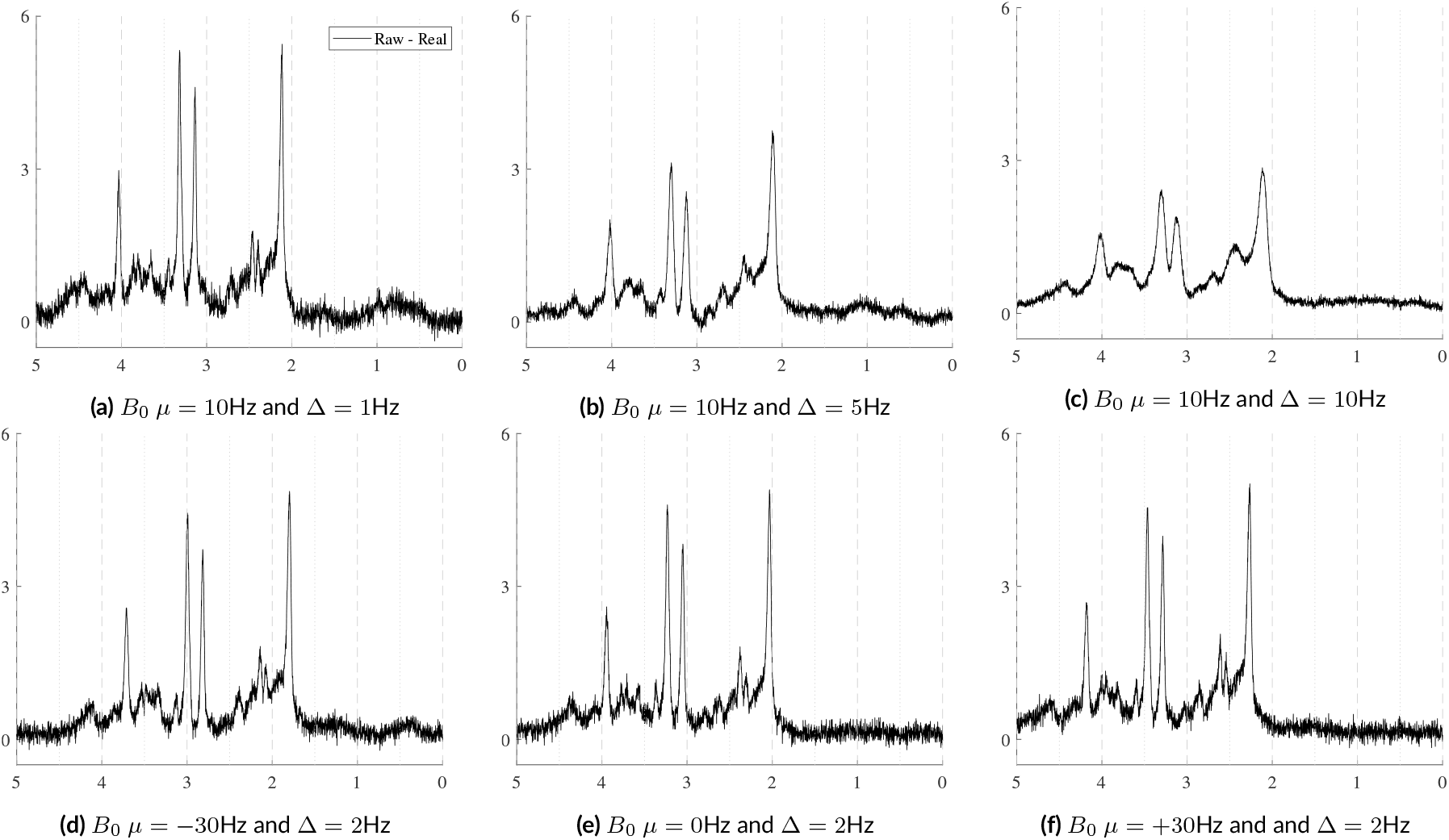
Illustration of the effects of *B*_0_ inhomogeneities. The top row shows the effects of increasing heterogeneity (Δ) within the voxel given a fixed mean *B*_0_ offset of 10Hz. The bottom row shows the effects of changing the mean *B*_0_ offset with a fixed distribution Δ across the voxel. Here, Δ is modeled as linear and isotropic and is defined as Δ = 2*dx* = 2*dy* = 2*dz*.

In general, the current approach mirrors Li *et al*. ^50^ in which the basis functions are modulated by the *B*_0_ field map, but in the implementation the *B*_0_ field map is simulated rather than acquired. As with MARSS, Li *et al*. ^50^ suggest using multiple points in each direction instead of a single value per voxel. The exact number of points used in each direction is described by the size of the spectroscopy voxel divided by the size of an anatomical imaging voxel. Any cuboidal shape, rectangular or otherwise, can be modeled. The *B*_0_ field is defined in Hertz by four variables: *±dx, ±dy, ±dz*, and *µ. dx, dy*, and *dz* describe half of the change in *B*_0_ in their respective axis from the voxel’s center and *µ* is the mean of the entire voxel. A linear *B*_0_ gradient is assumed, but different gradient profiles can be implemented by adding new functions that manipulate how the *x*−, *y*−, and *z*−components are calculated when generating the *B*_0_ field map. Section 3 in the Supporting Information provides a more detailed explanation of how the *B*_0_ is simulated.

#### 2.2.5 Baseline and Residual Water Signal

Proton MRS is particularly susceptible to spectral baseline offsets and residual water contamination. Currently, there is no physics-based model for simulating these components. In fact, it is unknown what a true baseline actually looks like. Theory and protocols regarding baseline modeling have been developed extensively, but the physical phenomena underlying the spectral baseline signal are still poorly understood. Therefore, different modeling protocols using different assumptions have been developed, producing different baselines. For example, Bazgir *et al*. ^51^ interpolate between selected minima in the spectra. Wilson *et al*. ^52^ use penalized B-splines to overparameterize the problem space which is solved using an optimized smoothing constraint. Simicic *et al*. ^53^ assumed the baseline came from macromolecular signal and fitted it by parameterizing the macromolecular spectrum into 10 components, each fitted with multiple Lorentzian lines, and using prior knowledge derived from in vivo data. And Osprey ^41^ uses unregularized B-splines spread every 0.4 ppm to enforce smoothness. Even though all of these techniques have proven useful, in vivo data does not have known ground truths meaning there is no way to identify which method is the most accurate. Similarly, the residual suppressed water region is also poorly characterized. Without a physics-based model, other methods of baseline and residual water simulation are necessary.

This work proposes a smoothed, pseudo-random, bounded walk generator for both the broad spectral baseline and the more irregular residual water region. Because these phenomena are poorly understood, a naive random model can be used in conjunction with observed constraints to approximate what is expected in vivo. The approach is elaborated on in Algorithm 1. Customizable profiles were developed for each artifact to more closely approximate what is expected in vivo. Immense variety of outputs can be achieved by randomly sampling the parameters from distributions instead of using fixed values. Once simulated, they are resampled to match the ppm range of acquired data and the order of magnitude is matched to the spectra. The corresponding complex components were generated by adding a 90 deg phase shift in the time-domain. These were then combined with the original real parts, finishing the complex-valued components. As shown in Fig. 5, this generator produces very different outputs depending on the specified configurations. Fig. 5a shows very broad, smooth lines while Fig. 5b shows highly irregular lines that closely resemble residual water regions. All outputs are then scaled to modulate the impact on the final spectra. For the baseline offsets, these scaling factors can diminish the baseline contribution and effectively smooth it, as shown in the bottom left plot of Fig. 5a. A more detailed exploration of this algorithm and the effects of each parameter are presented in the Supporting Information Section 4 Baseline and Residual Water Simulations.

**FIGURE 5.**
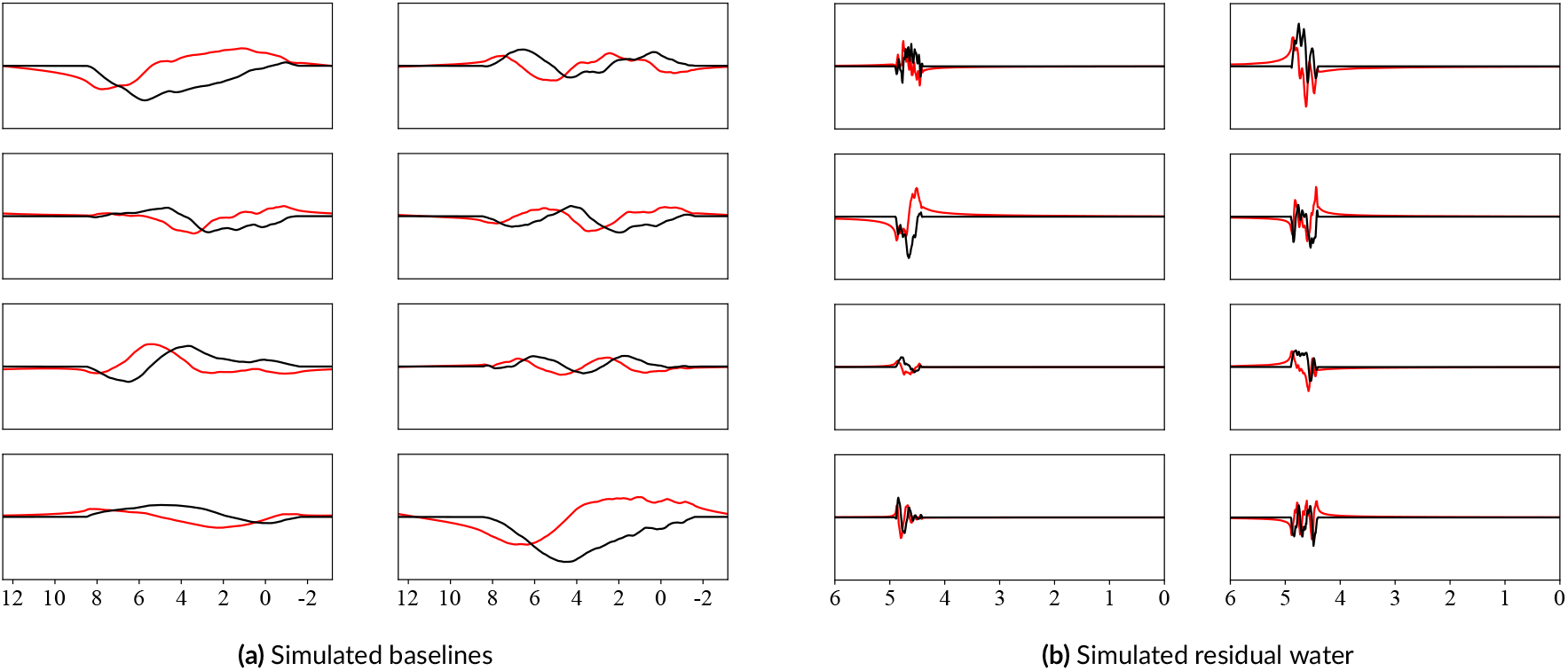
Simulated samples of spectral baselines and residual water regions using the pseudo-random bounded walk generator. The black and red lines are the real and imaginary components, respectively. The generated walks are smoothed and then added to the simulated spectra.

#### 2.2.6 Noise

Noise is always present in MRS, typically arising from sources such as coil temperature and the quantization in the analog-to-digital (ADC) converter. The signal-to-noise ratio (SNR) depends on the magnetic field strength and the nucleus being measured. This work uses the base definition of SNR from the terminology consensus paper: ^54^ one unit of signal divided by the standard deviation of the noise. Several options were provided, e.g. time-domain or frequency-domain and using peak height or signal power, but only required explicitly stating the adopted definition. Here, the spectral SNR is defined as the maximum peak height of the real component divided by one standard deviation of the noise. The input SNR is randomly sampled first and then the standard deviation for the noise is calculated using the maximum height of a metabolite of choice and the desired SNR. The noise in this model assumes a Gaussian distribution. The real and imaginary components of the noise are assumed to be uncorrelated and are therefore sampled using separate noise vectors, but defined with the same parameters for each component. Optionally, they can be correlated by sampling a single vector and then combined with a corresponding imaginary component using the procedure from 2.2.5.

##### Algorithm 1

Smoothed Bounded Pseudo-Random Walk

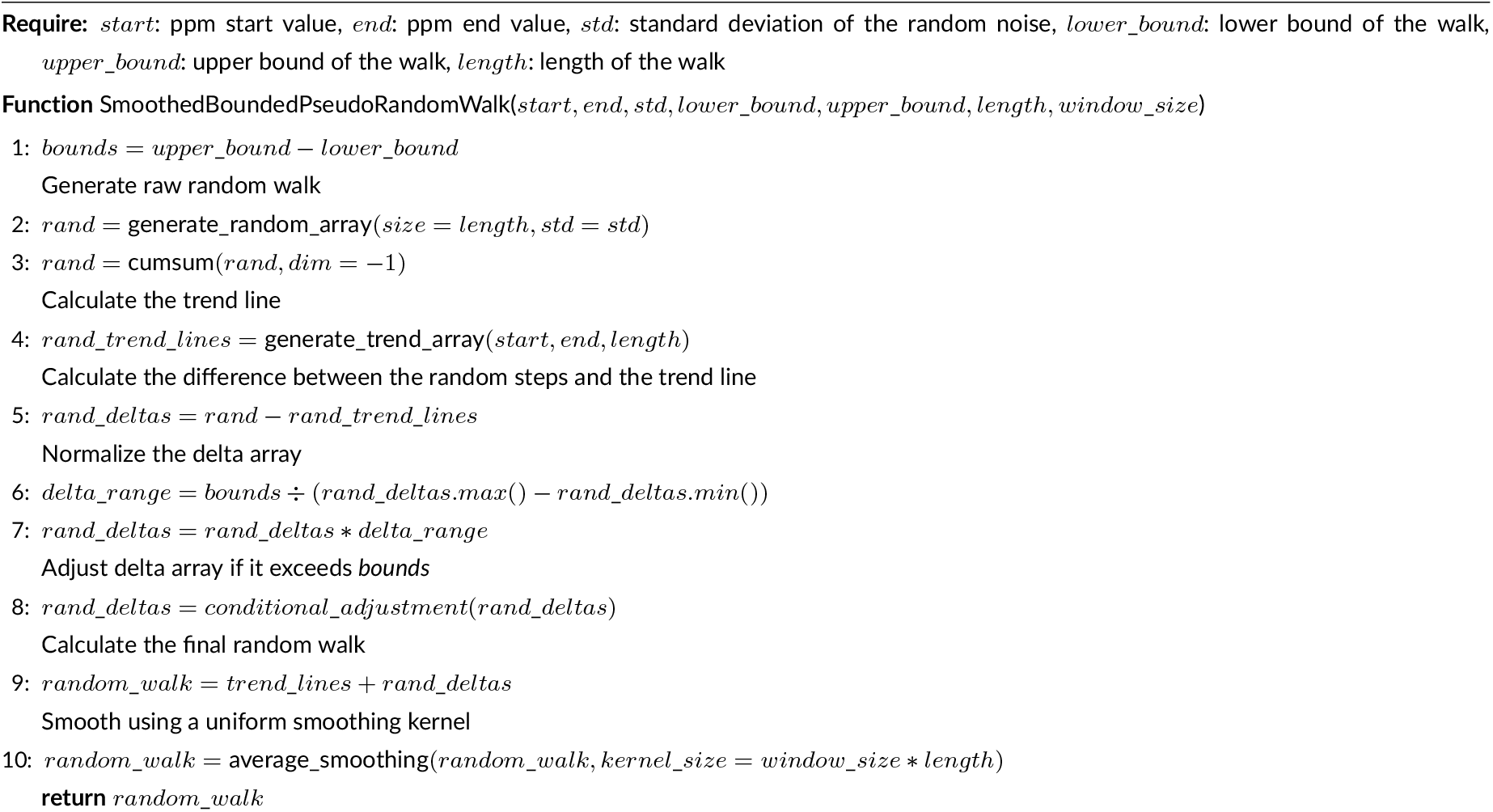

#### 2.2.7 Phase Offsets

##### Zero-Order Phase

FIDs and spectra are complex data types consisting of real and imaginary components. A 0° zero-order phase offset results in absorption and dispersion spectra in these components, respectively. An absorption spectrum exhibits peaks with idealized lineshapes that are purely positive or purely negative, while dispersion spectra exhibit anti-symmetrical peaks which are both positive and negative. As shown in Fig. 6, non-zero degree offsets result in a mixture of absorption and dispersion spectra. In its current form, a phase offset is selected randomly and independently during parameter sampling and applied in the time domain using the following complex exponential:

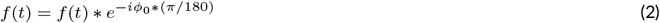

**FIGURE 6.**
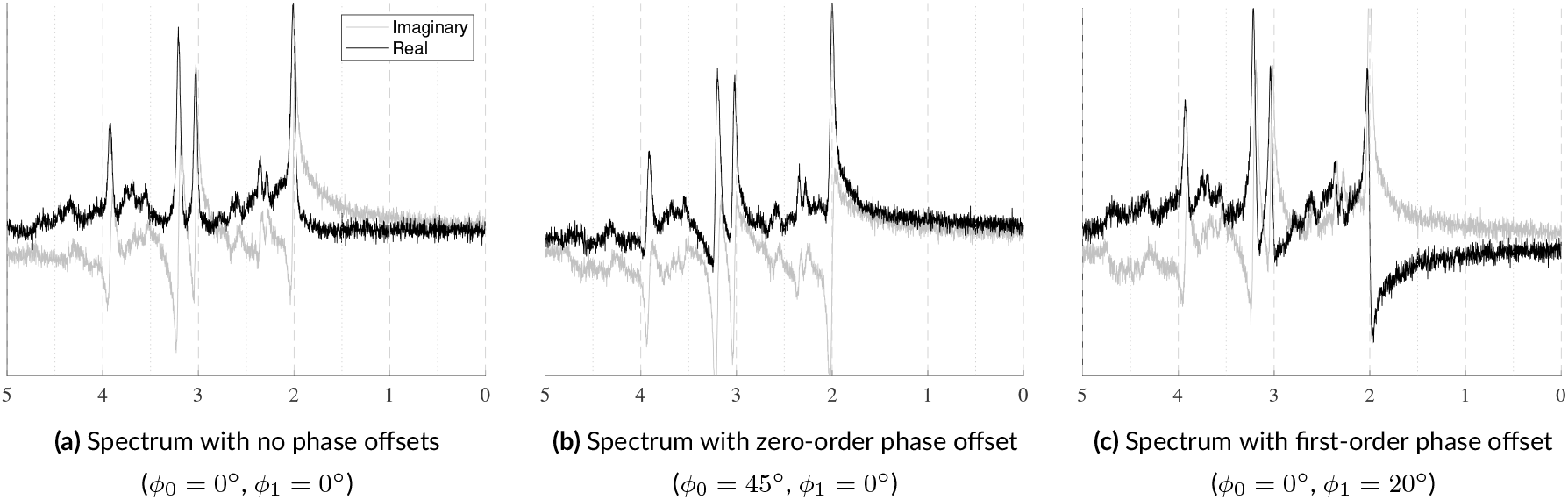
Samples illustrating how identical spectra are affected by zero- and first-order phase offsets. In 6a, the real component (black) is in absorption mode exhibiting narrow line widths and is fully positive. In 6b, a zero-order phase offset is applied. As the spectrum shifts from absorption to dispersion mode, the peaks uniformly lose their symmetry and negative values from the imaginary component are transferred to the real component. In 6c, a first-order phase shift is applied. This is evident by the increasing antisymmetry across the spectrum emanating from the water peak.

##### First-Order Phase

First-order phase, also referred to as linear phase, is a frequency-dependent linear offset that emanates from a reference point, i.e. the center frequency referred to as *ppm*_*ref*, which is typically the water peak at 4.65ppm. This offset is caused by a time delay between the end of the pulse sequence and the start of measuring the FID. A linear phase offset creates asymmetrical line shapes that grow larger as one moves away from the reference point. This is illustrated in Fig. 6 by comparing 6a and 6c. Currently, the *ppm*_*ref* term is fixed to 4.65ppm while the *ϕ*_1_ term is randomly sampled and then applied in the frequency domain using the following complex exponential:

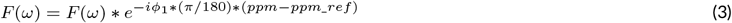

#### 2.2.8 Frequency Shifts

During data acquisition, the FID typically experiences a global frequency shift, primarily caused by the mean *B*_0_ offset. However, some functional groups can experience individual frequency shifts which are attributed to effects such as temperature and pH. Similar to Sec. 2.2.3, the MRS-Sim uses separate global frequency shifts for the metabolites and nuisance signals. However, this model also allows each basis function to have an independent frequency shift in addition to the global shift which is in line with common spectral fitting protocols. Both of these options are illustrated in Fig. 7. For more in vivo-like, realistic spectra, values can be used from the work by Wermter *et al*. ^55^, which characterized the temperature-induced frequency shift of several brain metabolite moieties with temperature sensitivity. As more metabolites are characterized for their temperature- and pH-sensitivities, this information can be added to simulate more realistic spectra. Currently available data will be included in the table in the Supporting Information Section 2.

**FIGURE 7.**
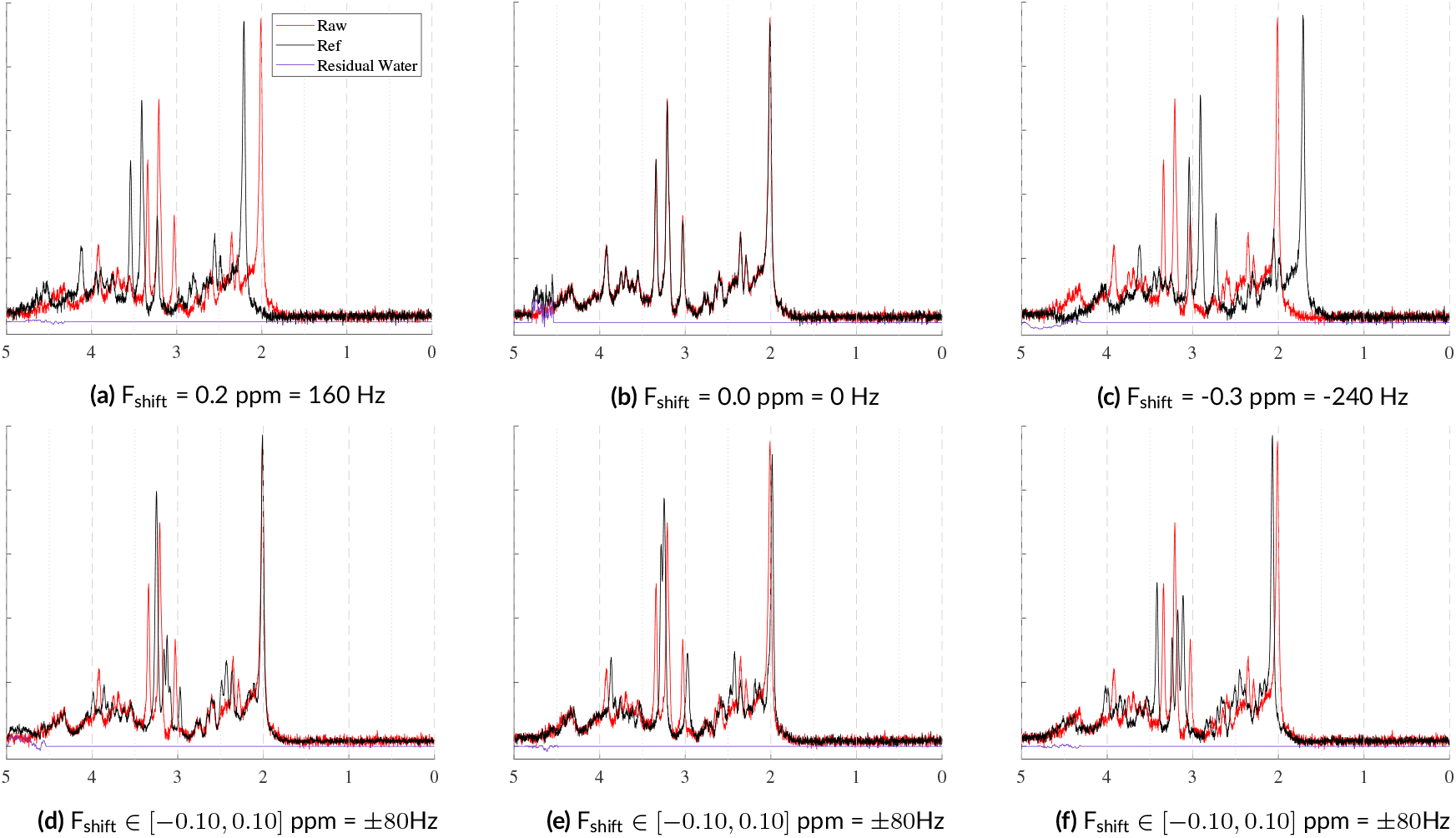
These examples illustrate two types of frequency shifts. The top row shows global frequency shifts in which a single shift is applied to the entire signal. The bottom row shows the effects of using randomly sampled metabolite-level frequency shifts. *±*80Hz is high, but was selected to improve the effect’s visibility. The spectral SNR = 15 with fixed Lorentzian and Gaussian broadening. No spectral baselines, macromolecules, phase offsets, or eddy currents were included.

#### 2.2.9 Eddy Currents

Eddy currents are sometimes induced by the rapidly switching gradients used during MR acquisitions. Eddy current correction techniques, such as the Klose ^56^, tend to be non-parameterized, making it difficult to model the exact effect of each approach. Near *et al*. in FID-A ^34^, however, provide a parameterized equation for simulating first-order eddy currents. These artifacts are applied as a function of amplitude, *A*, time constant, *tc*, and time, *t*. The time constant must be short enough that it occurs entirely within the recorded echo, otherwise it will appear as a simple, global frequency shift. The effects of eddy current variables can be seen in Fig. 8. The top row of Fig. 8 shows the effect of varying amplitude given a fixed time constant and the bottom row shows how an increasing time constant with a fixed amplitude impacts the spectra. Moderate to large amplitudes coupled with moderate to short time constants begin to create new resonances. And increasing amplitudes apply small, but increasing, frequency shifts. These examples highlight the importance of choosing parameter combinations carefully.

**FIGURE 8.**
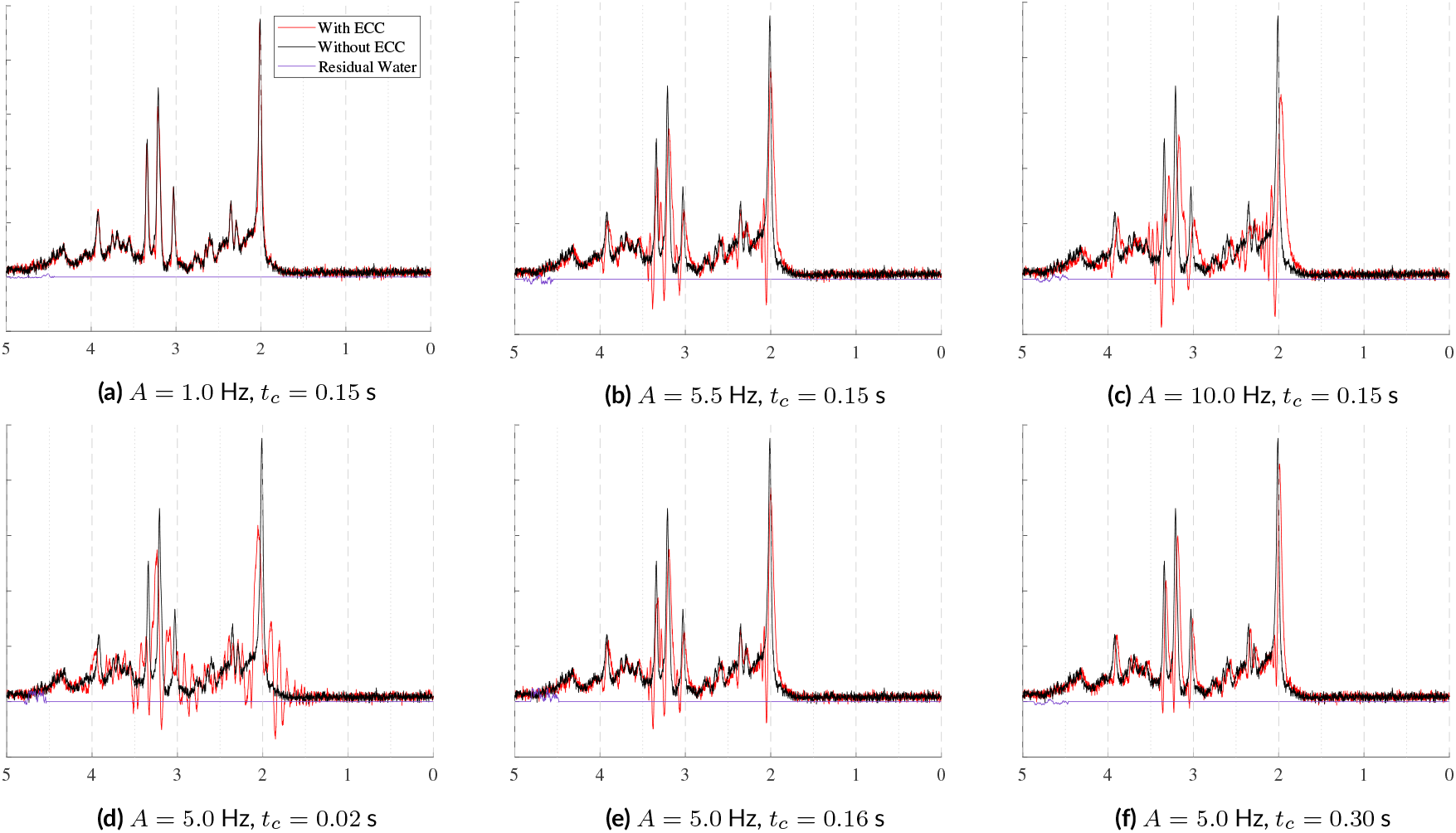
Samples illustrating eddy currents simulated with the two-parameter model from FID-A ^34^: an amplitude, A, and a time constant, *tc*. In the top row, *tc* is fixed to 0.15 seconds and the amplitudes are varied. In the bottom row, the amplitude is fixed to 5Hz and the time constants are varied. The parameter ranges used here come from FID-A ^34^.

#### 2.2.10 Spectral Transients

In this work, the term *spectral transients*, or simply *transients*, refers to one of two scenarios, depending on the context of the simulations. It can either refer to the signal from multiple individual RF coils measured from a phased-array receiver coil for a single average prior to coil combination or it can refer to multiple signal averages. Both multi-channel receiver coils and multiple signal averages are commonly used because averaging them together improves the SNR, which tends to be low for individual coil channels or signal averages. The terminology consensus paper by Kreis *et al*. ^54^ provides many clarifications, but does not specify how to differentiate between data before and after coil-combination nor specifically when a transient becomes a spectrum. For clarity and consistency, *spectrum* and *FID* will refer to data that has been coil-combined or signal averaged. When simulating transients, additional considerations need to be included. During acquisition, averages experience additional artifacts including zero-order phase and frequency drifts while multi-coil transients are affected by scaling, due to coil sensitivity, and decreased SNR values. Each of these are included in the proposed framework. To allow for maximum variation in the simulations, each of the aforementioned parameters can be sampled from distributions and are discussed below.

##### Noise

Averaging multi-coil or multi-average acquisitions leads to an SNR improvement of the final spectrum by a factor of the square root of the number of non-zero weighted transients. To vary the SNR among the transients, this model scales the target linear SNR according to the number of coils and then samples scaling factors from a narrow normal distribution to maintain the mean target SNR. These are sampled randomly, but noise correlations between multiple coils in a phased-array can also be added to the simulations when defining the coil SNR parameters.

##### Coil Sensitivity

A variety of coil combination techniques can be used to combine sets in vivo of multi-coil transients into coil-combined spectra. Hall *et al*. ^57^ present an overview of some of these methods, of which the following use weighting schemes: water signal, water SNR, signal to peak area, metabolite SNR, time domain combination, and PCA noise whitening with weighted summation. These methods can be simulated in the same way by defining a parametric distribution that, when sampled, produces groups of weighting factors similar to what is normally observed with in vivo acquisitions. The current default implementation samples scaling factors from a Gaussian distribution with *µ* = 1.0 and *std* = 0.5 that is subsequently clamped to the range [0,2].

Since these weights are defined in the parameter sampling scheme, which is rewritten for every dataset, the settings used for sampling can be easily redefined. Assigning these weights a context that corresponds to an in vivo protocol, such as water peak height or coil sensitivity maps, can help define the necessary parameter ranges and distributions when planning the simulations. A statistical analysis of the coil sensitivity values from a given in vivo dataset from a specific scanner will provide simulations that more closely approximate experimentally acquired data. This approach is in line with the preprocessing consensus paper that recommends always averaging transients ^45^ instead of summing them. However, if one wishes to sum the transients, then the scaling factors need to be normalized by their sum so as to maintain the overall amplitude. Future work can implement more complex combination techniques.

##### Frequency Drift and Phase Drift

Frequency drifts and phase drifts are phenomena observed in transients from multiple acquisitions in which each transient has an independent offset. These offsets and alignments are shown in Fig. 9.

**FIGURE 9.**
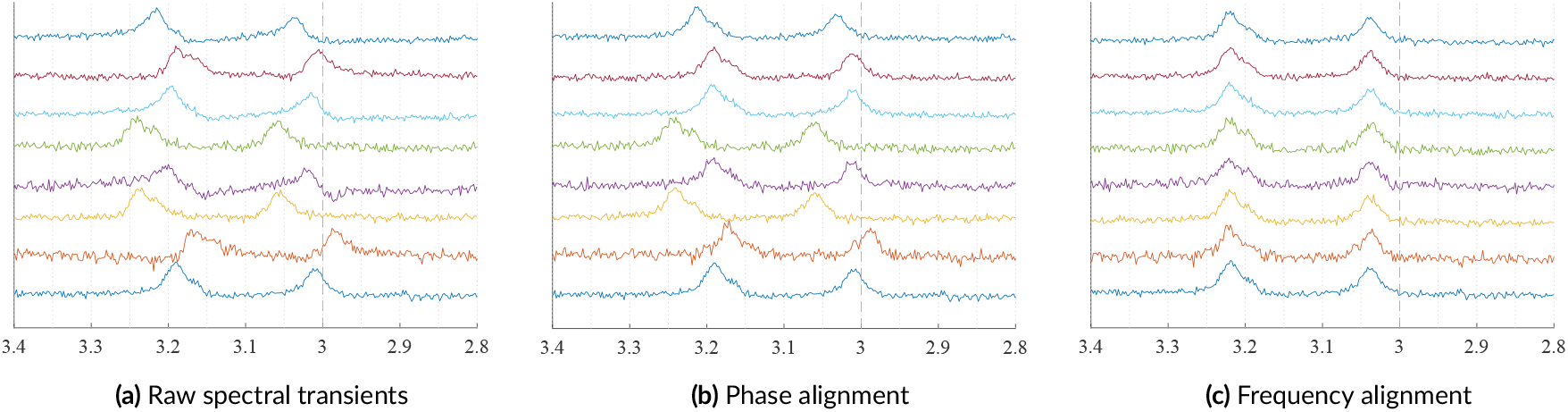
Examples of 8 simulated transients, which can simulate multiple coil elements or multiple signal averages. 9a shows the transients with various SNRs, and coil sensitivities, along with zero-order phase and frequency drifts. 9b shows those transients after phase alignment. Then, 9c shows those transients after frequency alignment.

#### 2.2.11 Final Steps

The desired use case will determine if a FID or a spectrum is necessary. If a FID is required, the simulation is finished and the data will be exported. If a spectrum is required, the Fourier transform will recover the spectrum at which point it can be cropped and resampled to a desired ppm range and spectral length. The default interpolation technique in this framework is a cubic Hermite modified Akima interpolator with momentum.

### 2.3 Exporting Data

The default export file format is .mat. These files include the data, spectral fits, simulation parameters, baseline offsets, and metabolite amplitude coefficients. To facilitate the use of the simulated spectra in various software packages, they are also exported in the NIfTI-MRS format ^58^.

### 2.4 Spectral Fitting Parameter Analysis

The process of simulating a new dataset requires careful consideration of various factors, including the selection of appropriate parameter ranges and distributions. Customization of these parameters depends on the intended use and application of the dataset. For example, when creating deep learning-based quantification models, it is beneficial to use independent, uniform distributions that include all values the model will be expected to encounter. On the other hand, when validating a traditional spectral fitting model that includes soft constraints, it is crucial to incorporate those constraints when defining the parameter distributions. This ensures that the simulated dataset accurately reflects the underlying distribution of the target dataset.

To create a dataset that accurately mimics in vivo conditions, it is crucial to have precise descriptions of in vivo fitting parameters. This work does not provide scenario-specific parameter recommendations, as it is beyond the scope of this work. However, some tools are provided to assist in identifying ranges and distributions to match an existing dataset. In collaboration with the developers of Osprey ^41^, new functionality was added to their software to export the parameters after spectral fitting. The tools in this framework can load those exported files and prepare the data for further analysis. Currently, this framework uses the Python library Fitter ^59^ to identify the best-fitting probability distribution for each parameter. *A priori* knowledge, either from prior knowledge or data exploration, can narrow down the search range and accelerate the analysis. The results for each parameter include evaluation metrics for the best performing distributions as well as a numerical characterization of the best fitting distribution.

### 2.5 Code

This repository was written in PyTorch 1.11.0 and Python 3.9.7. Since this framework generates batches of spectra instead of individual spectra sequentially, a simulation batch size needs to be specified which will be affected by the spectral length and complexity of the simulations. As long as the batch size is set appropriately given the users’ amount of RAM, this framework can be employed on standard computers without any special hardware. After publication, the repository will be available on GitHub, at https://github.com/JohnLaMaster/MRS-Sim, and MRSHub.

Metabolite and model parameter recommendations are provided as default values and are included in the Supporting Information Section 2 and the online repository. Model parameter values generally come from the default values of spectral fitting programs ^34,41,60^ while the parameters describing the metabolites come from literature ^38,43,44,55,61,62^. For the most current information, please refer to the repository.

#### 2.5.1 Data Evaluation

Validating synthetic data is challenging because there is no gold standard and the “realness” of the data depends on both the simulation methods, i.e. equations and components, and the simulation parameter values. This makes directly evaluating the simulated spectra difficult. The scope of this work is focused on the simulation framework itself instead of recommending in vivo parameter values. Even though there is no quantitative metric describing the degree of in vivo realness, it is still important to evaluate the model outputs qualitatively. Therefore, the most appropriate approach for evaluating this protocol is to evaluate the simulation methods described above and then visually assess the outputs. When simulating data to match an in vivo dataset, in vivo-synthetic pairs can be made by using the in vivo fitting parameters to simulate a corresponding spectrum. In such a case, a similarity metric or cross-correlation can be used to evaluate the similarity between the in vivo samples and their corresponding synthetic pairs.

The spectra in Fig. 10 show how the *B*_0_ simulator affects a spectrum with varying mean inhomogeneity offsets. The spectra in Fig. 11 show a single spectrum with various sampled baseline offsets and residual water signals. And finally, the examples in Fig. 12 demonstrate the complete model. Those spectra were generated by sampling and randomly applying all spectral artifacts, except zero-order phase because of how much it obfuscates spectral details.

**FIGURE 10.**
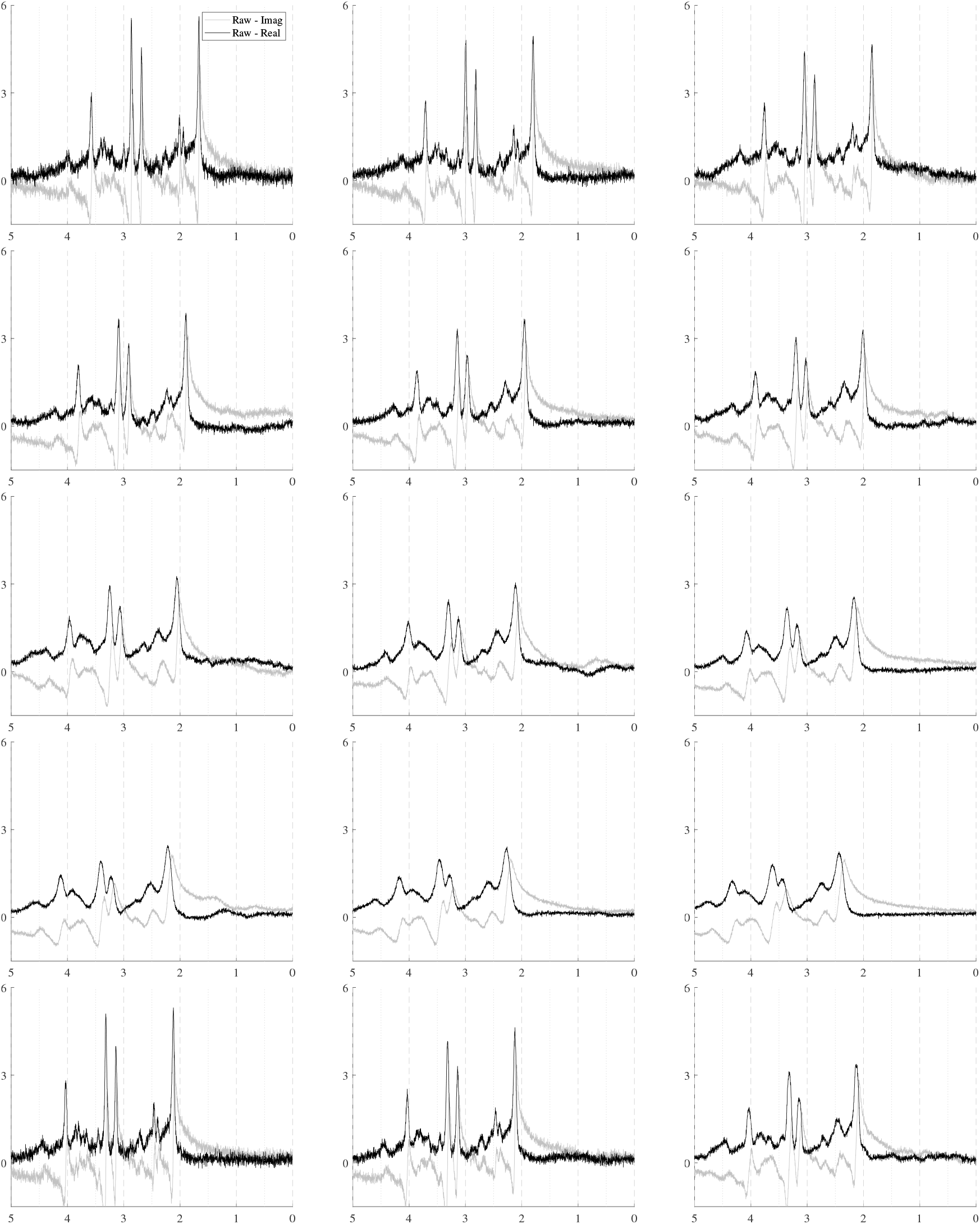
Illustrations of a single sample spectrum simulated for a PRESS sequence with TE=30ms that highlights the effect of *B*_0_ magnitude and distribution on the spectra. The mean offsets range from [-50, +50] Hz and while the inhomogeneities increase from Δ = [0.5, 15.0] Hz. As the inhomogeneities increase, the intensity of the spectra decreases. As the magnitude of the *B*_0_ offset increases, it causes increasing global frequency shifts. The bottom row used 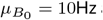 and Δ = [2.0, 5.0, 10.0] Hz.

**FIGURE 11.**
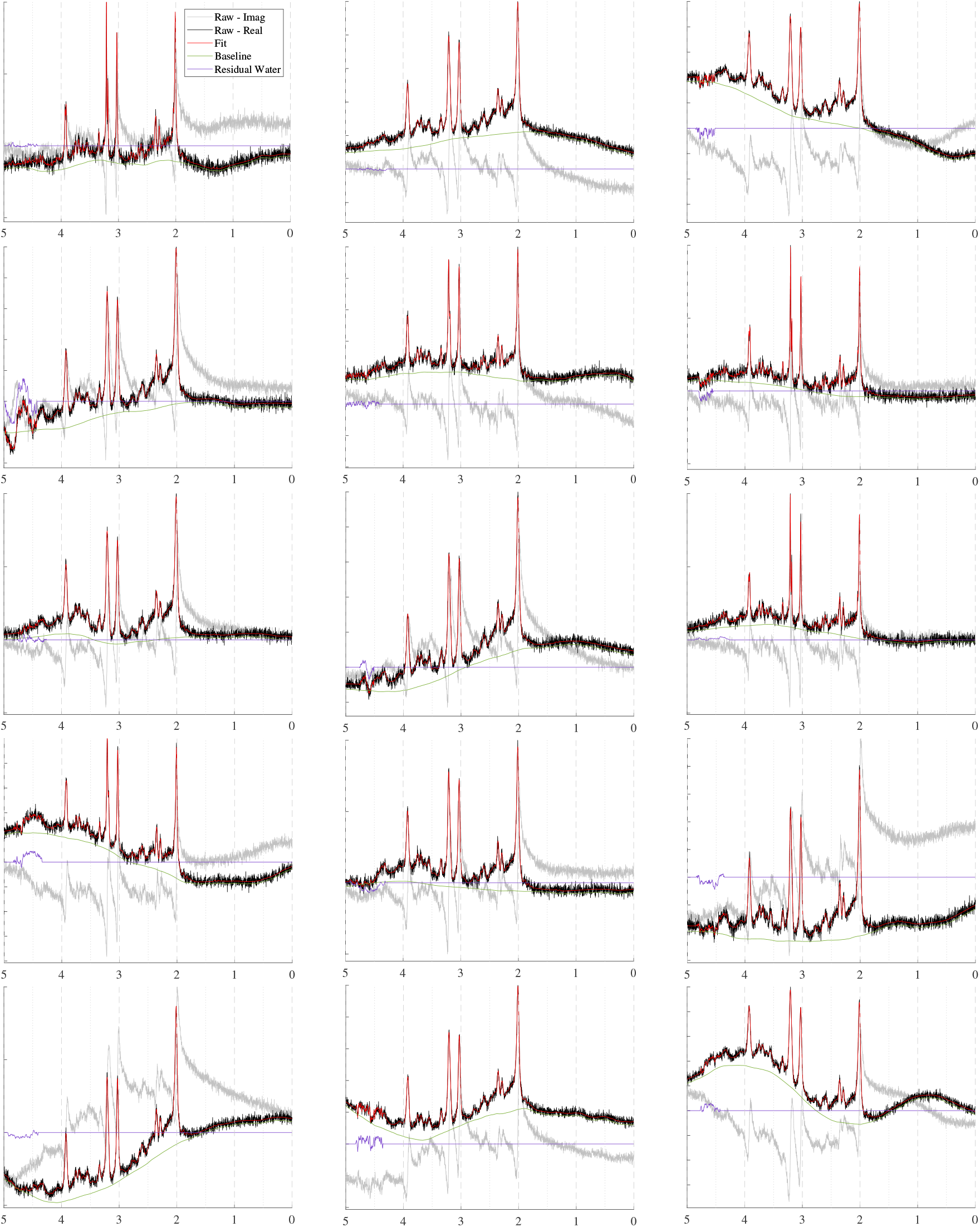
Sample spectra simulated for a PRESS sequence with TE=30ms that highlight the effect of the baseline and residual water contributions.

**FIGURE 12.**
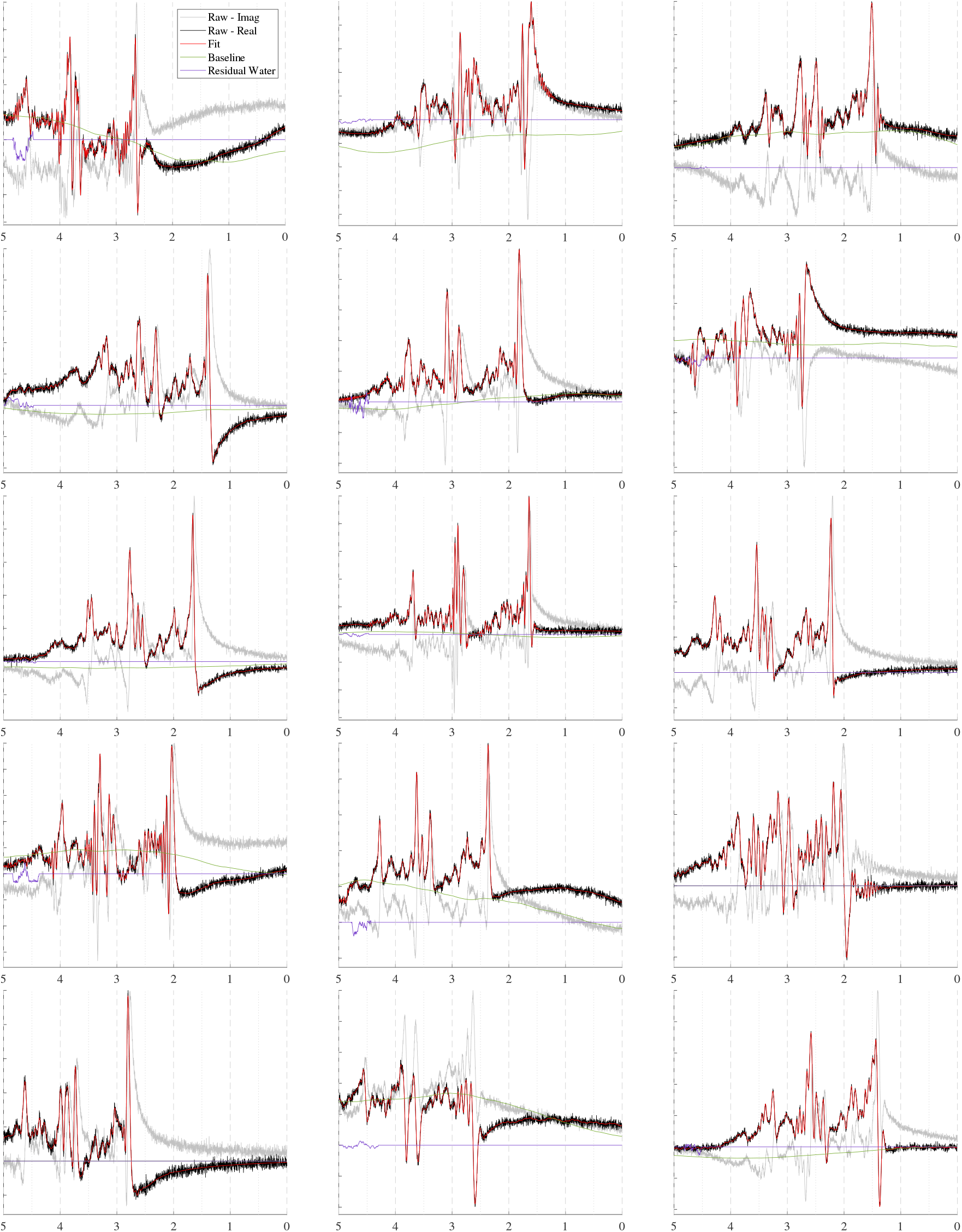
Sample spectra, similar to Fig. 11 [PRESS, TE=30ms], with randomly sampled, uncorrected artifacts to approximate raw data, including both good and bad quality. These plots highlight the comprehensiveness and flexibility of MRS-Sim simulations.

It is important to note that while pulse sequence implications are integral to the overall simulation process, their detailed exploration is beyond the scope of this work. Therefore, the simulations in Sec. 3 are restricted to a single pulse sequence and echo time. The main objective of this work is to manipulate pre-simulated basis sets until they approximate in vivo spectra, meaning the primary machinations of MRS-Sim are the same regardless of the basis functions used. Overall, the presented simulations using the PRESS sequence provide a valuable foundation for understanding the performance and capabilities of the MRS-Sim framework.

## 3 RESULTS

### 3.1 B_0_ Field Inhomogeneity Simulator

*B*_0_ field inhomogeneities cause three main effects: line broadening, signal damping, and a global frequency. Figure 10 shows these effects on a short echo (TE=30ms) spectrum that is corrupted by varying levels of *B*_0_ inhomogeneities. The *B*_0_ parameters increases from left-to-right and top-to-bottom as follows: *µ* = [−50, −30, −23; −17, −10, −3; 3, 10, 17; 23, 30, 50; 10, 10, 10] Hz and the isotropic inhomogeneities are [Δ = 0.50, 1.8, 3.0; 4.4, 5.8, 7.1; 8.4, 9.7, 11.0; 12.4, 13.7, 15.0; 1.0, 1.0, 1.0] Hz. The bottom row of Fig. 10 illustrates examples of anisotropic inho-mogeneities using Δ*z* = [2.0, 5.0, 10.0] Hz from left to right. This figure shows that even several hertz of inhomogeneities noticeably impact the spectral resolution and signal intensity. The mean *B*_0_ offset *µ* in a voxel is responsible for the induced frequency shift. Visually, there is no noticeable difference between isotropic and anisotropic inhomogeneities.

### 3.2 Baseline and Residual Water Generator

A variety of spectral baseline and residual water contributions are illustrated in Fig. 11. The parameter ranges can be selected so that the outputs look reasonable to the user. “Reasonable” is a subjective term because spectral baselines are still poorly characterized. Therefore, instead of claiming to produce in vivo baselines, the proposed generator produces a wide variety of spectral baseline offsets that approximate the baselines extracted by traditional methods. The motivation behind this approach is that if the generator cannot be limited to strictly in vivo-like baselines, then it should incorporate a variety of appropriate baseline profiles such that true baselines are included. Because this generator is not based on a baseline fitting method, future work could use it to evaluate the baseline modeling performance of various fitting protocols given different baseline profiles. The examples shown in Fig. 11 omit additional spectral artifacts to more clearly highlight the variety that can be achieved by randomly sampling parameters for the baseline and residual water generator.

### 3.3 Complete Model

In contrast, Fig. 12 presents a simulated PRESS spectrum (TE=30ms) with randomly sampled artifacts and offsets. This figure highlights the variety a single spectrum can assume just by randomly sampling the artifacts. Sophisticated simulations, such as these, are useful for developing and validating data processing techniques such as artifact removal and new spectral fitting protocols. Spectra can be simulated to be as clean or as artifact-laden as desired, all while maintaining known ground truth values.

Upon closer inspection of Fig. 12, the baselines and residual water regions sometimes appear to be uncorrelated with the depicted spectra, but they are in fact paired. These discrepancies are due to the additional artifacts applied to the simulations after all of the spectral components are combined. The plotted baseline and residual water regions are the ground truth signals and do not have those additional artifacts applied to them. Incorporating a robust collection of artifacts and phenomena commonly encountered with in vivo data into these simulations, makes resulting datasets more closely approximate unprocessed, in vivo data. Even though post-processing techniques are highly accurate, they have limitations and biases, leaving some residue of the corrected artifacts. Including these artifacts in the simulations and removing them with the users’ own post-processing and fitting protocols ensures consistency between simulated and in vivo data. These artifacts can also be scaled down in the simulations to emulate residual artifacts. However, this is not ideal because it cannot be guaranteed that scaling down the artifacts will accurately reflect the biases and residues found in the in vivo data.

Deep learning techniques also benefit from robust and consistent synthetic data. Ensuring congruence of the synthetic training data with the in vivo data promotes better model performance. While including fine-grained detail, such as sampling moiety-level *T*_2_ values, may not be essential for every use case, in a deep learning setting, it acts as a form of data augmentation. Such augmentation techniques maintain the underlying data but manipulate its appearance so that models learn a broader range of features instead of overfitting a single pattern. These techniques are standard practice in deep learning and are an inherent feature of this framework.

## 4 DISCUSSION

MRS-Sim represents a major step forward for the MRS field. This is the first flexible, open-source framework that combines realism, flexibility, and accessibility in a single, open-source platform. By integrating spectral components from state-of-the-art spectral fitting models, MRS-Sim provides a comprehensive and adaptable state-of-the-art signal model. Its modular design and flexibility make it a powerful tool for simulating data from diverse in vivo scenarios.

The increasing reliance on synthetic MRS data stems from the need for large, well-characterized datasets with known ground truth values. Synthetic data enables unlimited spectrum generation, facilitating applications ranging from traditional spectroscopy to deep learning-based approaches, including evaluating the accuracy and precision of data analysis methods. However, current synthetic data generation methods suffer from a lack of standardization, leading to reproducibility challenges and limiting the generalizability of findings. Existing simulation approaches often rely on simplifying assumptions that exclude some spectral components, reducing the fidelity of synthetic data. Similar to phantom data, these methods struggle to fully replicate in vivo spectra, particularly regarding artifacts and nuisance signals. Simulating realistic MRS data presents two major challenges: selecting the appropriate spectral components and artifacts to include and determining biologically relevant parameter values. These choices are difficult even for experts, and the variability in simulation approaches further exacerbates reproducibility issues.

MRS-Sim directly addresses these challenges by providing a comprehensive, open-source framework designed to enhance reproducibility, transparency, and usability of synthetic data. Unlike existing methods, it integrates a wide range of spectral components and acquisition-induced artifacts, making it the most complete publicly available simulation tool to date. There are several key features that set MRS-Sim apart. First is a *B*_0_ field simulator capable of introducing distortions into the simulations stemming from imperfect shimming and high susceptibility effects. This provides a unified model to apply non-relaxation-based broadening, signal damping, and a global frequency shift. The second is a semi-parametric generator that creates both broad, undulating spectral baseline offsets and highly irregular residual water contributions, enhancing the fidelity of simulated spectra. And finally, the modular architecture allows easy customization and expansion to accommodate evolving research needs and advancements.

This framework is not just a tool for generating synthetic spectra—it is a platform for advancing MRS research. Its flexibility enables simulations at any stage of the data acquisition pipeline, from individual coil elements to fully processed spectra, making it uniquely suited for testing and validating new spectral processing methods. Additionally, MRS-Sim fosters community-driven development, allowing researchers to propose, develop, and integrate new features, such as J-difference edited spectra, diffusion spectra, or 2D spectra, through GitHub. This collaborative model ensures continuous improvements and adaptability. To further support the community, MRS-Sim includes a comprehensive reference database in its appendix and digital repository. This resource provides up-to-date information on metabolite spin systems, temperature-induced artifacts, 3T *T*_2_ values, and metabolite concentration ranges. By maintaining a shared repository of validated parameters, MRS-Sim helps standardize synthetic data generation, offering researchers a consistent and well-documented starting point for their simulations.

## 5 CONCLUSIONS

The development of open-source frameworks for data generation is crucial to ensure that synthetic MRS data is widely adopted and to improve the generalizability of synthetic data and the reproducibility of MRS research. By making this framework widely available, the authors hope to further democratize MRS research by providing easy access to high-quality simulations and supporting information. Improving the quality of synthetic data will ultimately lead to better generalizability to in vivo data, directly improving the overall applicability of synthetic data in MRS.

Moving forward, collaboration and consensus-building among researchers will be essential to establish standards and best practices for simulating MRS data. Future developments can focus on expanding the framework to add additional in vivo scenarios, further broadening its utility to more fields within MRS. By addressing key challenges associated with synthetic MRS data generation and promoting standardization, this work provides a foundation for continued advancements in the field.

## Supporting information

MRS-Sim Supplement

## ABBREVIATIONS

Asc: ascorbate.
Asp: aspartate.
Ch: choline.
Cr: creatine.
DL: deep learning.
GABA: gamma-aminobutyric acide.
Gln: glutamine.
Glu: glutamate.
GPC: glycerophosphorylcholine.
GSH: glutathione.
Lac: lactate.
LASER: localized adiabatic selective refocusing.
MARSS: Magnetic Resonance Spectrum Simula tor.
mI: myo-inositol.
ML: machine learning.
NAA: N-acetylaspartate.
NAAG: N-acetylaspartylglutamate.
PCh: phosphocholine.
PCr: phosphocreatine.
PE: phosphatidylethanolamine.
PRESS: point resolved spectroscopy.
sI: scyllo-inositol.
sLASER: semi-localized adiabatic selective refocusing.
STEAM: stimulated echo acquisition method.
Tau: taurine.
tCr: total Creatine.

## ACKNOWLEDGMENTS

The authors would like to thank Dr. Tobias Lasser for his mentorship during this work; Dr. Dhritiman Das for discussions about his experience with simulating MRS SVS data; the Harper House for their support while visiting UCSF to further develop the physics model; and the DAAD and Technical University of Munich’s Graduate Center of Computation, Information and Technology for helping to finance the stay at UCSF in San Francisco, USA.

## Funding Information

This work was supported in part by a doctoral fellowship from the German Academic Exchange Service (DAAD) and the following grants from the National Institutes of Health: R00 AG062230, R01 CA262630, and R21 EB033516.

## Conflict of interest

The authors declare no potential conflict of interest.

## Notes

### Competing Interest Statement

The authors have declared no competing interest.

### Summary of Updates

Terminology and wording have been clarified along with other revisions in the course of peer review. An additional SI section was added for terminology. Many figures were re-plotted to be more uniform. Redundancy between figures in the methods and results sections was eliminated in favor of the more comprehensive figures and explanations. Additional changes were made in the course of peer review.

