## Supplementary material for "MRS-Sim: Open-Source Framework for Simulating In Vivo-like Magnetic Resonance Spectra": MRS-Sim Supplement

### Supporting Information: MRS-Sim

#### 1 | TERMINOLOGY

1. **Basis function**: A basis function is the quantum mechanical simulation of the resonances excited by a given pulse sequence and defined by the spin system of a given metabolite. Semi-parametric basis functions also exist that approximate macromolecules and lipid contributions.

2. **Baseline**: The baseline of a spectrum is the component of the signal that does not contain metabolite information. In a properly pre-processed and fitted spectrum, the baseline should fall along the x-axis at *signal=0*.

3. **Baseline offset**: A spectral artifact that adds non-metabolite signal non-uniformly across frequencies. In this work, a baseline offset does not include macromolecular components, but could include lipids, if explicitly stated.

4. **In vivo scenario**: This refers to specific setups for in vivo acquisitions, e.g. using a 3T GE clinical scanner to acquire TE=35ms SVS PRESS data or a 9.4T Brucker scanner to acquire TE=144ms MEGA-edited MRSI data. These scenarios are defined by the characteristics and parameters necessary to simulate their basis sets.

5. **Offset**: An offset is a spectral component or artifact that can be positive or negative and, when corrected, converges to zero but is never removed. For example, phase and frequency shifts are offsets that are corrected to have $0 \deg$ and $0Hz$ offsets, respectively. The non-macromolecule baseline is an offset that, once removed, should be a flat line along the x-axis.

6. **Parameter**: In this work, parameters are the variables used in signal models to either simulate or fit a spectrum. Fitting parameters include all variables optimized during the spectral fitting process.

7. **Parameter distribution**: This refers to the statistical characterization of the values for a given parameter within a dataset. For example, the estimated NAA concentrations might follow a normal (Gaussian) distribution described by a mean and standard deviation. The SNR values of a dataset might follow a beta distribution defined by $\alpha$ and $\beta$. Once identified, these distributions can be used to simulate more in vivo-like data.

8. **Processing, pre- & post-**: Data processing refers to the protocols of correcting or removing artifacts in the data to prepare it for spectral fitting. Pre-processing and post-processing refer to the same procedure but are used when the focus is on spectral fitting or data acquisition, respectively.

9. **Quantification model**: This is a deep learning-based method to estimate metabolite concentrations. This is different from spectral fitting because it only estimates the metabolite concentrations and none of the remaining fitting parameters. Such an approach is not possible in traditional spectroscopy.

#### 2 | SIMULATION PARAMETERS

The following table contains spin definitions and recommended simulation parameter ranges for 35 brain metabolites and water. This table is based on literature values and does have some gaps. The recommended ranges represent the minimum and maximum values across physiological and pathological differences. The concentration ranges were scaled to represent $[0.9 * min, 1.1 * max]$. To compare a variety of sources and to find the most up-to-date values, please check the GitHub repository.

The data provided here is an excerpt of the database. So far, data has been compiled from textbooks[4, 5] and commonly referenced literature sources[2, 3] for defining the spin systems. The chemical shifts and coupling constants have also been reported from MARRS[8] since that is the recommended software for simulating basis sets. MARRS also compiled several sources, including Near *et al.*'s[9] work describing the GABA spin systems. $T_2$ values were collected from Wyss *et al.*[11] who reported moiety-specific values for selected metabolites and from Gudmundson *et al.*'s[6] recent meta-analysis. Concentration ranges were compiled from De Graaf's textbooks[4, 5] and Gudmundson's meta-analysis[6]. Macromolecule and lipid nuisance signal characterizations come from Gudmundson et. al[6] and Hamilton *et al.*[7]. Finally, Wermter *et al.*'s[10] work describing moiety-level temperature-induced frequency shifts has also been included. The following citations were used to compile the current version of the database. As new information is added, the citation list in the repository will be updated:

| Name | Moieties | Group | Chemical Shift (ppm) [4, 5] | Interaction | Scalar coupling (Hz) [4, 5] | $T_2$ [ms] | | Conc. [mM] | |
|---|---|---|---|---|---|---|---|---|---|
| | | | | | | Low | High | Low | High |
| Acetate | – | $^{2}CH_3$ | 1.904 | – | – | – | – | – | – |
| Alanine | – | $^{2}CH$ | 3.775 | 2-3 | 7.230 | 100.00 | 250.00 | 0.090 | 1.650 |
| | | $^{3}CH_3$ | 1.467 | 3-3' | – | 100.00 | 250.00 | | |
| | | – | – | 3-3" | – | 100.00 | 250.00 | | |
| | | – | – | 3'-3" | – | 100.00 | 250.00 | | |
| Ascorbic Acid | – | $^{4}CH$ | 4.492 | 4-5 | 2.07 | 73.00 | 250.00 | 0.367 | 2.072 |
| | | $^{5}CH$ | 4.002 | 5-6 | 6.00 | 73.00 | 250.00 | | |
| | | $^{6}CH_2$ | 3.743 | 5-6' | 7.60 | 73.00 | 250.00 | | |
| | | – | 3.716 | 6-6' | -11.50 | 73.00 | 250.00 | | |
| Aspartate | – | $^{2}CH$ | 3.891 | 2-3 | 3.647 | 36.00 | 204.550 | 0.900 | 7.668 |
| | | $^{3}CH_2$ | 2.804 | 2-3' | 9.107 | 36.00 | 204.550 | | |
| | | – | 2.670 | 3-3' | -17.426 | 36.00 | 204.550 | | |
| 2-Hydroxy-glutarate [1] | – | $^{2}CH$ | 4.022 | 2-3 / 2-3' | 7.600 / 4.100 | – | – | – | – |
| | | $^{3}CH_2$ | 1.825 | 3-3' | -14.00 | – | – | | |
| | | – | 1.977 | 3-4 / 3-4' | 5.300 / 10.400 | – | – | | |
| | | $^{4}CH_2$ | 2.221 | 3'-4 / 3'-4' | 10.600 / 6.00 | – | – | | |
| | | – | 2.272 | 4-4' | -15.00 | – | – | | |
| Choline | – | $N(CH_3)_3$ | 3.185 | – | 0.570 | 119.00 | 394.00 | 0.00 | 2.750 |
| | | $^{1}CH_2$ | 4.054 | 1-2 / 1'-2' | 3.140 / 3.168 | 93.00 | 345.00 | | |
| | | – | – | 1-1' | 2.572 | 93.00 | 345.00 | | |
| | | $^{2}CH_2$ | 3.501 | 1'-2 / 1-2' | 7.011 / 6.979 | 93.00 | 345.00 | | |
| | | – | – | 2-2' | 2.681 | 93.00 | 345.00 | | |
| Creatine | – | $N(CH_3)$ | 3.027 | – | – | 104.00 | 242.700 | 0.325 | 16.505 |
| | | $^{2}CH_2$ | 3.913 | – | – | 76.00 | 213.800 | | |
| | | NH | 6.650 | – | – | 164.080 | 242.700 | | |
| Ethanol-amine | – | $^{1}CH_2$ | 3.818 | 1-2 / 1'-2' | 3.897 / 3.798 | – | – | 0.00 | 1.650 |
| | | – | – | 1-1' / 1-N | -10.640 / 0.657 | – | – | | |
| | | $^{2}CH_2$ | 3.147 | 1'-2 / 1-2' | 6.694 / 6.794 | – | – | | |
| | | – | – | 2-2' / 1'-N | -11.710 / 0.1420 | – | – | | |
| $\gamma$-Amino-butyric acid | – | $^{2}CH_2$ | 2.287 | 2-2' | -15.938 | – | – | 0.033 | 10.720 |
| | | – | – | 2-3 / 2-3' | 7.678 / 6.980 | 25.00 | 229.00 | | |
| | | – | – | 2'-3 / 2'-3' | 6.980 / 7.678 | 25.00 | 229.00 | | |
| | | $^{3}CH_2$ | 1.892 | 3-3' | -15.00 | 25.00 | 229.00 | | |
| | | – | 1.895 | 3-4 / 3-4' | 8.510 / 6.503 | 25.00 | 229.00 | | |
| | | $^{4}CH_2$ | 3.003 | 4-4' | -14.062 | 25.00 | 229.00 | | |
| | | – | 3.005 | 3'-4 / 3'-4' | 6.503 / 8.510 | 25.00 | 229.00 | | |
| Glucose 36% $\alpha$ / 64% $\beta$ | GlcA | Refer to GlcA below | | | | 38.00 | 250.00 | 0.00 | 2.200 |
| | GlcB | Refer to GlcB below | | | | 40.00 | 250.00 | 0.00 | 2.200 |
| Glucose, $\alpha$-anomer (36% = 0.36 1H) | – | $^{1}CH$ | 5.216 | 1-2 | 3.8 | 38.00 | 250.00 | 0.00 | 2.200 |
| | | $^{2}CH$ | 3.519 | 2-3 | 9.6 | 38.00 | 250.00 | | |
| | | $^{3}CH$ | 3.698 | 3-4 | 9.4 | 38.00 | 250.00 | | |
| | | $^{4}CH$ | 3.395 | 4-5 | 9.9 | 38.00 | 250.00 | | |
| | | $^{5}CH$ | 3.822 | 5-6 | 1.5 | 38.00 | 250.00 | | |
| | | $^{6}CH$ | 3.826 | 5-6' | 6.0 | 38.00 | 250.00 | | |
| | | $^{6'}CH$ | 3.749 | 6-6' | -12.1 | 38.00 | 250.00 | | |
| Glucose, $\beta$-anomer (64% = 0.64 1H) | – | $^{1}CH$ | 4.630 | 1-2 | 8.0 | 40.00 | 250.00 | 0.00 | 2.200 |
| | | $^{2}CH$ | 3.230 | 2-3 | 9.1 | 40.00 | 250.00 | | |
| | | $^{3}CH$ | 3.473 | 3-4 | 9.4 | 40.00 | 250.00 | | |
| | | $^{4}CH$ | 3.387 | 4-5 | 8.9 | 40.00 | 250.00 | | |
| | | $^{5}CH$ | 3.450 | 5-6 | 1.6 | 40.00 | 250.00 | | |
| | | $^{6}CH$ | 3.882 | 5-6' | 5.4 | 40.00 | 250.00 | | |
| | | $^{6'}CH$ | 3.707 | 6-6' | -12.3 | 40.00 | 250.00 | | |

*(continued on the next page)*

| Name | Moieties | Group | Chemical Shift (ppm) [4, 5] | Interaction | Scalar coupling (Hz) [4, 5] | $T_2$ [ms] Low | $T_2$ [ms] High | Conc. [mM] Low | Conc. [mM] High |
|---|---|---|---|---|---|---|---|---|---|
| Glutamine | — | $^{2}CH$ | 3.767 | 2-3 / 2-3’ | 5.846 / 6.500 | 57.00 | 234.00 | 2.700 | 6.600 |
| | | $^{3}CH_2$ | 2.137 | 3-4 / 3-4’ | 9.165 / 6.324 | 57.00 | 234.00 | | |
| | | — | 2.121 | 3-3’ | -14.504 | 57.00 | 234.00 | | |
| | | $^{4}CH_2$ | 2.431 | 3’-4 / 3’-4’ | 6.347 / 9.209 | 57.00 | 234.00 | | |
| | | — | 2.458 | 4-4’ | -15.371 | 57.00 | 234.00 | | |
| | | $^{5}NH_2$ | 6.816 | — | — | 57.00 | 234.00 | | |
| | | $^{2}NH_2$ | 7.529 | — | — | 57.00 | 234.00 | | |
| Glutamate | — | $^{2}CH$ | 3.748 | 2-3 / 2-3’ | 7.331 / 4.651 | 50.00 | 219.580 | 2.692 | 20.774 |
| | | $^{3}CH_2$ | 2.046 | 3-3’ | -14.849 | 50.00 | 219.580 | | |
| | | — | 2.120 | 3-4 / 3-4’ | 6.413 / 8.406 | 50.00 | 219.580 | | |
| | | $^{4}CH_2$ | 2.334 | 3’-4 / 3’-4’ | 8.478 / 6.875 | 50.00 | 219.580 | | |
| | | — | 2.352 | 4-4’ | -15.915 | 50.00 | 219.580 | | |
| Glycine | — | $^{2}CH_2$ | 3.547 | — | — | 28.00 | 206.00 | 0.820 | 2.395 |
| Glycerol | — | $^{1}CH_2$ | 3.552 | 1-2 | 4.427 | — | — | — | — |
| | | — | 3.640 | 1-1’ | -11.715 | — | — | | |
| | | $^{2}CH$ | 3.770 | 1’-2 / 2-3’ | 6.485 / 6.485 | — | — | | |
| | | $^{3}CH_2$ | 3.552 | 2-3 | 4.427 | — | — | | |
| | | — | 3.640 | 3-3’ | -11.715 | — | — | | |
| Glycerophospho-choline | Glycerol moiety | $^{1}CH_2$ | 3.605 | 1-2 / 1’-2 | 5.7700 / 4.5300 | 101.00 | 371.00 | 0.033 | 10.720 |
| | | — | 3.672 | 1-1’ | -14.78 | 101.00 | 371.00 | | |
| | | $^{2}CH$ | 3.903 | 2-3 / 2-3’ | 5.77 / 4.53 | 101.00 | 371.00 | | |
| | | $^{3}CH_2$ | 3.871 | 3-3’ | -14.78 | 101.00 | 371.00 | | |
| | | — | 3.946 | 3,3’-P | 6.03 / 6.03 | 101.00 | 371.00 | | |
| | Phospho-choline moiety | $^{1}CH_2$ | 4.312 | 1-2 / 1-2’ | 3.10 / 5.90 | 124.00 | 396.00 | | |
| | | — | — | 1’-2 / 1’-2’ | 5.90 / 3.10 | 124.00 | 396.00 | | |
| | | $^{2}CH_2$ | 3.659 | 1-1’ / 2-2’ | 2.67 / 2.67 | 124.00 | 396.00 | | |
| | | — | — | 1,1’-P | 6.03 / 6.03 | 124.00 | 396.00 | | |
| | | $N(CH_3)_3$ | 3.212 | — | — | 124.00 | 396.00 | | |
| Glutathione | Glycine moiety | $^{2}CH_2$ | 3.769 | — | — | 38.00 | 188.360 | 0.118 | 3.804 |
| | | NH | 7.154 | — | — | 38.00 | 188.360 | | |
| | Cysteine moiety | $^{2}CH$ | 4.561 | 2-3 | 7.09 | 33.00 | 188.360 | | |
| | | $^{3}CH_2$ | 2.926 | 2-3’ | 4.71 | 33.00 | 188.360 | | |
| | | — | 2.975 | 3-3’ | -14.06 | 33.00 | 188.360 | | |
| | | NH | 8.177 | — | — | 33.00 | 188.360 | | |
| | Glutamate moiety | $^{2}CH$ | 3.769 | 2-3 / 2-3’ | 6.34 / 6.36 | 34.00 | 225.00 | | |
| | | $^{3}CH_2$ | 2.159 | 3-4 / 3-4’ | 6.7 / 7.6 | 34.00 | 225.00 | | |
| | | — | 2.146 | 3-3’ | -15.48 | 34.00 | 225.00 | | |
| | | $^{4}CH_2$ | 2.510 | 3’-4 / 3’-4’ | 7.6 / 6.7 | 34.00 | 225.00 | | |
| | | — | 2.560 | 4-4’ | -15.92 | 34.00 | 225.00 | | |
| Homocarnosine | GABA moiety | $^{2}CH_2$ | 2.969 | 2-3 / 2-3’ | 8.00 / 8.00 | — | — | 0.00 | 0.440 |
| | | — | 2.944 | 2-2’ | -12.500 | — | — | | |
| | | $^{3}CH_2$ | 1.896 | 3-3’ | -13.900 | — | — | | |
| | | — | 1.881 | 2-3’ / 2’-3 | 7.500 / 7.500 | — | — | | |
| | | $^{4}CH_2$ | 2.378 | 3’-4 / 3’-4’ | 7.500 / 7.500 | — | — | | |
| | | — | 2.348 | 3-4 / 3-4’ | 7.500 / 7.500 | — | — | | |
| | | — | — | 4-4’ | -15.200 | — | — | | |
| | Alanine moiety | $^{2}CH$ | 4.467 | 2-3 | 5.020 | — | — | | |
| | | $^{3}CH_2$ | 3.191 | 3-3’ | 8.640 | — | — | | |
| | | — | 3.013 | 3-3’ | -15.300 | — | — | | |
| | Imidazole moiety | $^{5}CH$ | 7.08s | 4-NH | 0.580 | — | — | | |
| | | $^{2}CH$ | 8.08s | 2-NH | 1.130 | — | — | | |

*(continued on the next page)*

| Name | Moieties | Group | Chemical Shift (ppm) [4, 5] | Interaction | Scalar coupling (Hz) [4, 5] | $T_2$ [ms] Low | $T_2$ [ms] High | Conc. [mM] Low | Conc. [mM] High |
|---|---|---|---|---|---|---|---|---|---|
| Lactate | – | $^{2}CH$ | 4.097 | 2-3 | 6.930 | 61.00 | 226.520 | 0.00 | 15.824 |
| | | $^{3}CH_3$ | 1.313 | – | – | 61.00 | 226.520 | | |
| myo-Inositol | – | $^{1}CH$ | 3.522 | 1-2 | 2.889 | 19.00 | 439.00 | 1.380 | 23.393 |
| | | $^{2}CH$ | 4.054 | 2-3 | 3.006 | 19.00 | 439.00 | | |
| | | $^{3}CH$ | 3.522 | 3-4 | 9.997 | 19.00 | 439.00 | | |
| | | $^{4}CH$ | 3.614 | 4-5 | 9.485 | 19.00 | 439.00 | | |
| | | $^{5}CH$ | 3.269 | 5-6 | 9.482 | 19.00 | 439.00 | | |
| | | $^{6}CH$ | 3.614 | 1-6 | 9.998 | 19.00 | 439.00 | | |
| N-acetylaspartate | Acetyl moiety | $^{2}CH_3$ | 2.008 | – | – | 125.00 | 411.00 | 2.465 | 25.344 |
| | Aspartate moiety | $^{2}CH$ | 4.382 | 2-3 | 3.861 | 109.00 | 376.00 | | |
| | | $^{3}CH_2$ | 2.673 | 2-3' | 9.821 | 109.00 | 376.00 | | |
| | | – | 2.486 | 3-3' | -15.592 | 109.00 | 376.00 | | |
| | | NH | 7.820 | 2-NH | 6.400 | 242.700 | 320.170 | | |
| N-acetylaspartyl-glutamate | Glutamate moiety | $^{2}CH$ | 4.128 | 2-3 / 2-3' | 7.330 / 4.650 | 36.00 | 216.110 | 0.119 | 3.182 |
| | | $^{3}CH_2$ | 1.881 | 3-3' | -14.850 | 36.00 | 216.110 | | |
| | | – | 2.049 | 3-4 / 3-4' | 6.410 / 8.410 | 36.00 | 216.110 | | |
| | | $^{4}CH_2$ | 2.190 | 3'-4 / 3'-4' | 8.480 / 6.880 | 36.00 | 216.110 | | |
| | | – | 2.180 | 4-4' | -15.920 | 36.00 | 216.110 | | |
| | | NH | 7.950 | 2-NH | – | 36.00 | 216.110 | | |
| | Acetyl moiety | $^{2}CH_3$ | 2.042 | – | – | 69.00 | 229.00 | | |
| | Aspartate moiety | $^{2}CH$ | 4.607 | 2-3 / 2-3' | 4.412 | 27.00 | 234.00 | | |
| | | $^{3}CH_2$ | 2.721 | 2-3' | 9.515 | 27.00 | 234.00 | | |
| | | – | 2.519 | 3-3' | -15.910 | 27.00 | 234.00 | | |
| | | NH | 8.260 | 2-NH | – | 27.00 | 234.00 | | |
| Phenylalanine | Alanine moiety | $^{2}CH$ | 3.975 | 2-3 | 5.209 | – | – | – | – |
| | | $^{3}CH_2$ | 3.273 | 2-3' | 8.013 | – | – | | |
| | | – | 3.105 | 3-3' | -14.573 | – | – | | |
| | Phenyl moiety | $^{2}CH$ | 7.322 | 2-3 / 2-4 | 7.909 / 1.592 | – | – | | |
| | | $^{3}CH$ | 7.420 | 2-3' / 2-2' | 0.493 / 1.419 | – | – | | |
| | | $^{4}CH$ | 7.369 | 3-4 / 3-3' | 7.204 / 0.994 | – | – | | |
| | | $^{5}CH$ | 7.420 | 2'-3 / 3'-4 | 0.462 / 7.534 | – | – | | |
| | | $^{6}CH$ | 7.322 | 2'-4 / 2'-3' | 0.970 / 7.350 | – | – | | |
| Phosphoryl-choline | – | $^{1}CH_2$ | 4.282 | 1-2 / 1-2' | 2.284 / 7.231 | 93.00 | 345.00 | 0.007 | 4.288 |
| | | – | – | 1'-2 / 1'-2' | 7.326 / 2.235 | 93.00 | 345.00 | | |
| | | $^{2}CH_2$ | 3.643 | 1,1'-N | 2.680 / 2.772 | 93.00 | 345.00 | | |
| | | – | – | 1-1' / 2-2' | -14.89 / -14.19 | 93.00 | 345.00 | | |
| | | – | – | 1,1'-P | 6.298 / 6.249 | – | – | | |
| | | $N(CH_3)_3$ | 3.209 | – | – | 119.00 | 394.00 | | |
| Phospho-creatine | – | $N(CH_3)$ | 3.029 | – | – | 130.00 | 210.00 | 0.779 | 10.123 |
| | | $^{2}CH_2$ | 3.930 | – | – | 100.00 | 180.00 | | |
| | | $^{1}NH$ | 6.58s | – | – | 130.00 | 210.00 | | |
| | | $^{2}NH$ | 7.30s | – | – | 130.00 | 210.00 | | |
| Phosphoethanol-amine | – | $^{1}CH_2$ | 3.977 | 1-2 / 1-2' | 3.182 / 6.716 | 34.00 | 250.00 | 0.00 | 2.200 |
| | | – | – | 1'-2 / 1'-2' | 7.204 / 2.980 | 34.00 | 250.00 | | |
| | | $^{2}CH_2$ | 3.216 | 1-1' / 2-2' | -14.560 / -14.710 | 34.00 | 250.00 | | |
| | | – | – | 1,1'-P | 7.288 / 7.088 | 34.00 | 250.00 | | |
| | | – | – | 1,1'-N | 0.464 / 0.588 | – | – | | |
| Pyruvate | – | $^{3}CH_3$ | 2.358 | – | – | 100 | 250 | 0.090 | 0.550 |
| scyllo-Inositol | – | $^{1\text{-}6}CH$ | 3.340 | – | – | 73.00 | 250.00 | 0.00 | 0.563 |

(continued on the next page)

| Name | Moieties | Group | Chemical Shift (ppm) [4, 5] | Interaction | Scalar coupling (Hz) [4, 5] | $T_2$ [ms] | | Conc. [mM] | |
|---|---|---|---|---|---|---|---|---|---|
| | | | | | | Low | High | Low | High |
| Serine | – | $^{2}CH$ | 3.835 | 2-3 | 5.979 | – | – | 0.00 | 2.200 |
| | | $^{3}CH_2$ | 3.937 | 2-3' | 3.561 | – | – | | |
| | | – | 3.976 | 3-3' | -12.254 | – | – | | |
| Succinate | – | $^{2\text{-}3}CH_2$ | 2.394 | – | – | – | – | 0.090 | 0.550 |
| Taurine | – | $^{1}CH_2$ | 3.420 | 1-2 / 1-2' | 6.740 / 6.460 | 66.00 | 231.140 | 0.00 | 6.600 |
| | | – | – | 1-1' | -12.438 | 66.00 | 231.140 | | |
| | | $^{2}CH_2$ | 3.246 | 1'-2 / 1'-2' | 6.400 / 6.790 | 66.00 | 231.140 | | |
| | | – | – | 2-2' | -12.930 | 66.00 | 231.140 | | |
| Threonine | – | $^{2}CH$ | 3.578 | 2-3 | 4.917 | – | – | 0.00 | 0.550 |
| | | $^{3}CH$ | 4.246 | 3-4 | 6.350 | – | – | | |
| | | $^{4}CH_3$ | 1.316 | – | – | – | – | | |
| Tryptophan | Alanine moiety | $^{2}CH$ | 4.047 | 2-3 | 8.145 | – | – | – | – |
| | | $^{3}CH_2$ | 3.475 | 2-3' | 4.851 | – | – | | |
| | | – | 3.290 | 3-3' | -15.368 | – | – | | |
| | Indole moiety | $^{2}CH$ | 7.312 | – | – | – | – | | |
| | | $^{4}CH$ | 7.726 | 4-5 / 4-6 | 7.600 / 1.00 | – | – | | |
| | | $^{5}CH$ | 7.278 | 4-7 | 0.945 | – | – | | |
| | | $^{6}CH$ | 7.197 | 5-6 / 5-7 | 7.507 / 1.200 | – | – | | |
| | | $^{7}CH$ | 7.536 | 6-7 | 7.677 | – | – | | |
| Tyrosine | Alanine moiety | $^{2}CH$ | 3.928 | 2-3 | 5.147 | – | – | – | – |
| | | $^{3}CH_2$ | 3.192 | 2-3' | 7.877 | – | – | | |
| | | – | 3.037 | 3-3' | – | – | – | | |
| | Phenol moiety | $^{2}CH$ | 7.186 | 2-2' / 2-3 | 2.538 / 7.981 | – | – | | |
| | | $^{3}CH$ | 6.890 | 3-3' | 2.445 | – | – | | |
| | | $^{5}CH$ | 6.890 | 2-3' | 0.311 | – | – | | |
| | | $^{6}CH$ | 7.186 | 2'-3 / 2'-3' | 0.460 / 8.649 | – | – | | |
| Valine | – | $^{2}CH$ | 3.595 | 2-3 | 4.405 | – | – | – | – |
| | | $^{3}CH$ | 2.259 | 3-4 | 6.971 | – | – | | |
| | | $^{4}CH_3$ | 1.028 | 3-4' | 7.071 | – | – | | |
| | | $^{4'}CH_3$ | 0.977 | – | – | – | – | | |
| Water | – | $H_20$ | 4.650 | – | – | – | – | – | – |

**TABLE S2.1** Table of recommended $T_2$ values and concentration ranges covering physiological and pathological ranges for selected brain metabolites. The $T_2$ values can vary by moiety within a metabolite, so they were matched with their respective spin systems. When basis sets consist of individual spins instead of metabolite-level basis functions, it will be necessary to align the spins to correspond properly with the developed parameter sampling scheme.

### 3 | $B_0$ FIELD SIMULATOR

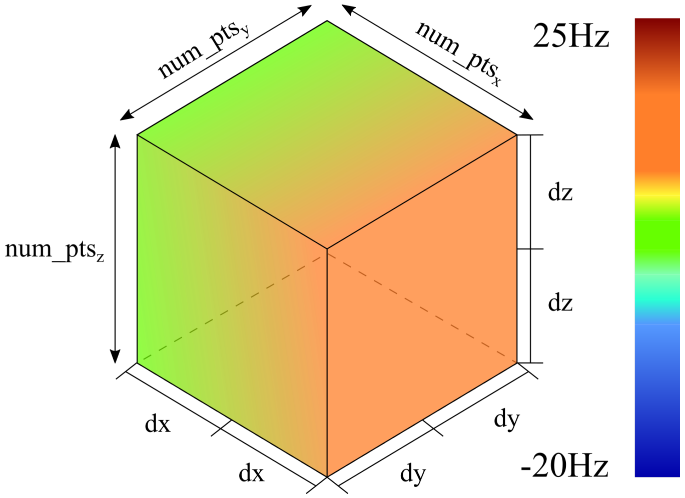


**FIGURE S3.1** $B_0$ field map model diagram. The labels along the x-, y-, and z-dimensions correspond to the terms of the mathematical model for simulating the $B_0$ field volume. Consider the x-dimension, $num\_pts_x$ refers to the corresponding number of points in the simulation and $2dx$ is the average corresponding change in the $B_0$ field across the voxel. The modeling procedure and each of the terms listed above are described in detail below.

As described in Li *et al.*[1], measuring the $B_0$ field map and using it to modulate the basis functions prior to fitting showed significant improvement in metabolite quantification and the fit residual surrounding the metabolite peaks. The fact that this technique has a direct impact on metabolite quantification shows the importance of including it when simulating spectra. The first step is to define the spectral and imaging resolutions of the simulated acquisitions. The defaults, shown below, assume a cubic spectroscopy voxel, but any cuboidal shape is acceptable.

**spectroscopy voxel** [10 mm x 10 mm x 10 mm]

**imaging voxel** [0.5mm x 0.5mm x 0.5mm]

The dimensions of the volume being modeled are the same as the spectroscopy voxel. The number of points included in the simulation is the quotient of the spectroscopy resolution and the anatomical imaging resolution.

$$\mathbf{num\_pts} = \frac{\text{spectroscopy resolution}}{\text{imaging resolution}} = \left[\frac{\text{10mm}}{\text{0.5mm}}_X, \frac{\text{10mm}}{\text{0.5mm}}_Y, \frac{\text{10mm}}{\text{0.5mm}}_Z\right] \tag{S3.1}$$

Once the model volume has been defined, the actual $B_0$ field, $dB0$, can then be modeled. Magnetic fields do not have hard, disjointed heterogeneities, but smooth variations. This is true even in the presence of high susceptibility effects causing strong distortions as long as the voxels are reasonably small. This model therefore assumes linearly varying gradients across the volume of the voxel. This assumption allowed the problem to be reduced to four variables:

$\mathbf{dx}$ half of the mean change in the x-direction

$\mathbf{dy}$ half of the mean change in the y-direction

$\mathbf{dz}$ half of the mean change in the z-direction

$\mu$ the mean offset of the entire voxel

The next step defines the gradients in each direction as shown in Eqns. S3.2, S3.3, and S3.4. As mentioned above, this model assumes linearly varying gradients, but more complex gradients can be implemented manually at this stage.

$$\mathbf{x} = \text{linspace}(\textbf{-dx}, \textbf{dx}, \textbf{num\_pts}_X) \tag{S3.2}$$

$$\mathbf{y} = \text{linspace}(\textbf{-dy}, \textbf{dy}, \textbf{num\_pts}_Y) \tag{S3.3}$$

$$\mathbf{z} = \text{linspace}(\textbf{-dz}, \textbf{dz}, \textbf{num\_pts}_Z) \tag{S3.4}$$

Once the gradients have been defined, the field map can be modeled very easily, as shown in Eqn. S3.5. The primary challenge in this step is reshaping the tensors to get the correct n+3D output.

$$\mathbf{dB0} = \mathbf{x} * \mathbf{y} * \mathbf{z} + \mu, \text{ such that} \tag{S3.5}$$

$$\mathbf{dB0}.size() = torch.Size([batchSize, \ldots, \mathbf{num\_pts}_{\mathrm{X}}, \mathbf{num\_pts}_{\mathrm{Y}}, \mathbf{num\_pts}_{\mathrm{Z}}]) \tag{S3.6}$$

Modulation due to the heterogeneous magnetic field can be applied in both the time-domain and the frequency-domain. In the time-domain, the complex exponential is applied via multiplication with the basis functions, as is shown below in Eqn. S3.7. In the frequency-domain, the complex exponential needs to be convolved with the spectrum. In both mathematical and practical terms, it is simpler to apply the modulation in the time-domain. Therefore, this model has only implemented the time-domain modulation.

$$F(t) = \sum_{n}^{N} M_n * basisfcn_n * \underbrace{\sum_{r=1}^{R} e^{-i\Delta\omega_r t}}_{B_0\ inhomog.}, \text{ where } \Delta\omega = \mathbf{dB0}.flatten(dims = [-3 : -1]) \text{ and } R = cumprod(\mathbf{num\_pts}) \tag{S3.7}$$

In this equation, the model **dB0** is flattened along the last three dimensions representing the **x**-, **y**-, and **z**-dimensions respectively. Therefore, **r** represents the linear indices from [1,R] corresponding the subscripts of the points within the 3D volume. The effective magnetic field at each of those points within the voxel has a direct effect on each moeity that resonates and contributes to the overall signal. Because of this, each basis function is modulated independently before summation.

### 4 | BASELINE AND RESIDUAL WATER SIMULATIONS

A single simulation protocol is used to generate the contributions for both baseline offsets and residual water contributions. The capability of generating signal contributions with such widely varying profiles is entirely dependent on the underlying parameters. In order to better understand the capabilities of the proposed generator and its behavior, the following section will explore the effects the various parameters have on the final signal contributions that are generated. For a detailed explanation of how these sampled parameters are converted into their final outputs, please refer to Algorithm 1 in Sec. 2.1.5 of the main manuscript.

This generator uses a smoothed, pseudo-random bounded walk algorithm for the simulations. For simplicity, this will henceforth be referred to as a *(random) bounded walk*, or simply a *walk*. By using different configuration dictionaries, it is possible to switch from very smooth, undulating baselines to rather erratic and rough residual water regions. The following sections highlight the effects different parameters have on the simulations.

#### 4.1 | Variables

These simulations use at least 9 degrees of freedom with two additional optional inputs:

1. Starting height (start)
2. Ending height (end)
3. Standard deviation of walk (std)
4. Lower bound (lower)
5. Upper bound (upper)
6. Point density (pt_density)
7. PPM range (ppm_range)
8. Length of smoothing window (window)
9. Scale (scale)
10. Dropout probability (drop_prob)
11. *Prime (prime)

Floats or integers can be provided for the point density, prime, and drop_prob. Everything else should be provided as a list with either a single value or a range in the form of [min_range, max_range]. When a single value is provided, that variable will be fixed. When a range is specified, then values will be sampled from that range uniformly. The point density and the PPM range are used to calculate the length of the walks. This length coupled with $std$ control the flexibility of the raw simulations. The length of the smoothing window, $window$, the magnitude of the $std$, and scaling factor, $scale$, of the baseline will affect the smoothness of the resulting walk. It is specified as a fraction of the length of the walk, so that it is independent of walk length, and can also be either fixed or sampled. $scale$ determines how prominent each offset will be when added to the simulated data. $drop\_prob$ is used to randomly omit the offset from that percentage of the simulations. $prime$ [units: ppm] is used for generating the residual water regions. This allows the region to vary in length by sampling two values in the range of $[-prime, prime)$ that can expand, contract, or slightly offset the residual water region.

#### 4.2 | Post-processing

Once the random walks have been generated, they are then smoothed with kernels of either fixed or varying width. At this point, trend lines are calculated between the starting and ending points which are then removed so that they start and end on the x-axis. Afterwards, the walks are normalized to [0,1] and then scaled down according to $scale$. A function called *sim2acquired* then uses zero padding and a nonuniform interpolator to add tails to both sides of the walks so that they match the PPM range of the basis set regarding the spectral width and carrier frequency. Before being added to the FIDs, they are multiplied by the maximum value of the FID in the frequency domain to scale them up to the correct order of magnitude. This makes $scale$ relative to the maximum height in the spectra. The offsets are added to the FIDs in the frequency domain before an inverse FFT returns the data to the time domain.

The parameters presented below control the length of the walks, their trend lines, flexibility, and smoothness. The default values presented in the configuration dictionaries were determined through preliminary experiments to closely approximate what was observed in a clinical dataset.

#### 4.3 | Preparing for the simulations

The following section defines parameters for the spectroscopy scenario and the configuration dictionary for the simulations.

```
# Spectroscopy scenario
spectralwidth = 2000                    # Hz
Ns            = 2048                    # number of spectral points
B0            = 3.0                     # T
gamma_H       = 42.577478518            # MHz/T
ppm_ref       = 4.65                    # ppm

carrier_frequency = B0 * gamma_H        # MHz
ppm = torch.linspace(-0.5*spectralwidth,
                     0.5*spectralwidth,Ns) # Hz
ppm /= carrier_frequency                # ppm
ppm += ppm_ref
t = torch.linspace(0,Ns/spectralwidth,Ns)

num_samples = 10
```

#### 4.4 | Understanding the Simulation Plots

The plots below contain either one or two lines that are blue or red. The default color scheme is as follows:

**blue** raw random walk simulation

**red** smoothed walk that is added to the simulated spectra

In Sections 4.5.4 and 4.6.5, which discuss how to incorporate the walks into the simulated spectra, only a single blue line is plotted. This line is the smoothed, real component of the walk. In Sections 4.5.5 and 4.6.6 which show the effect of applying the Hilbert transform, the blue and red lines represent the real and imaginary components, respectively. Similarly, Sections 4.5.6 and 4.6.7, which include examples of random simulations for each contribution type, also show the real and imaginary components using blue and red, respectively.

#### 4.5 | Baseline Offsets

The following section will explore this generator using the baseline configuration dictionary defined below. The values selected for the plots in the following sections were tailored specifically to the baseline simulations.

```
baseline_cfg = {
        "start":            [    -1,      1],
        "end":              [    -1,      1],
        "upper":            [             1],
        "lower":            [            -1],
        "std":              [  0.05,   0.20],
        "window":           [  0.15,    0.3],
        "pt_density":          128,
        "ppm_range":        [  -1.6,    8.5],
        "scale":            [     0,    1],
        "drop_prob":           0.0
}
```

##### 4.5.1 | Standard Deviation

This section shows the effect of different $std$ values on the random walk. The $std$ variable is used when sampling the noise for the walk. It directly controls the amount of variation between two consecutive points prior to the cumulative summation.

**FIGURE S4.1** Standard Deviation :: STD = 0.05; Window length = random; Kernel_size = random; Point density = 128

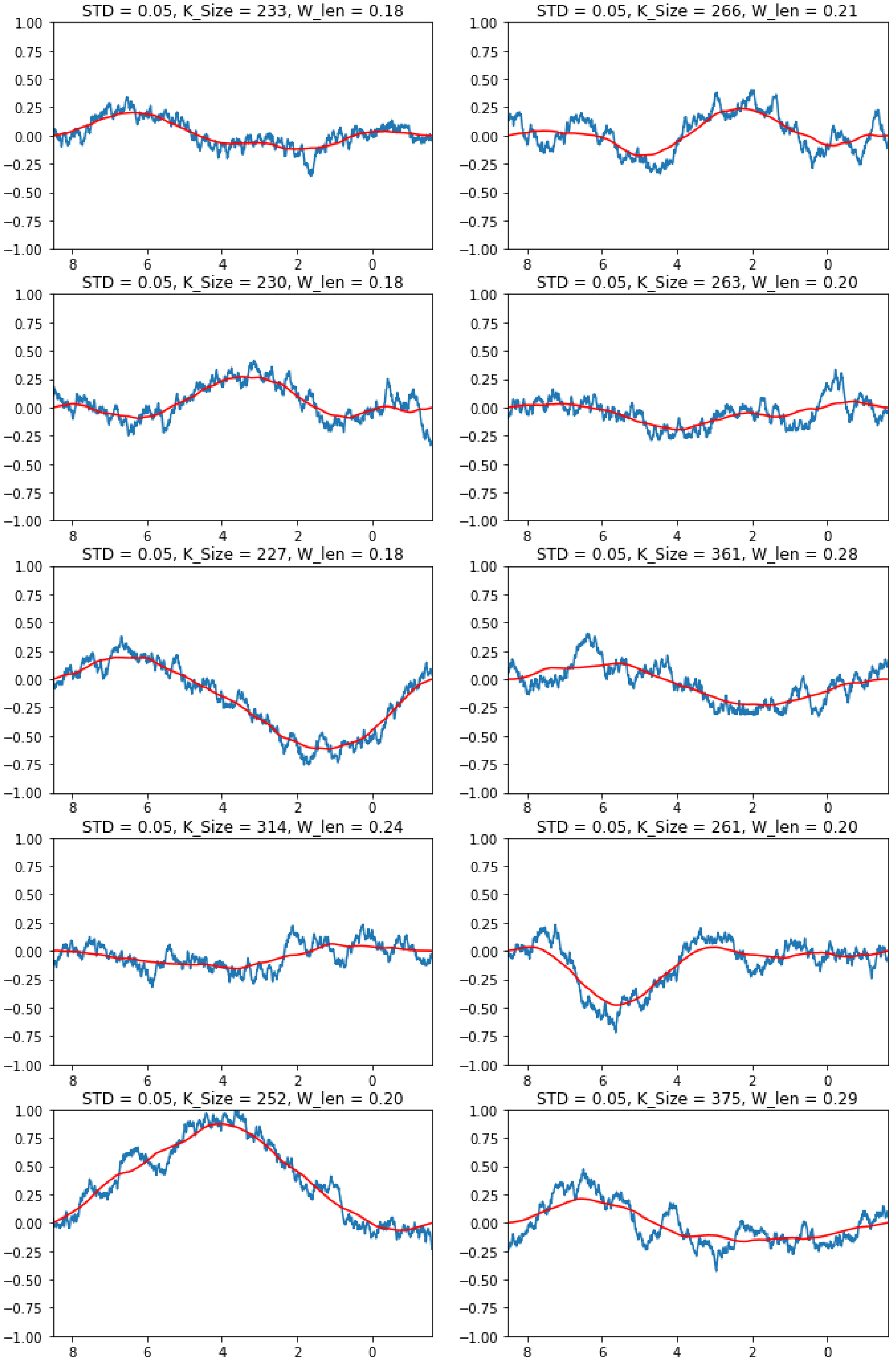

**FIGURE S4.2** Standard Deviation :: STD = 0.10; Window length = random; Kernel_size = random; Point density = 128

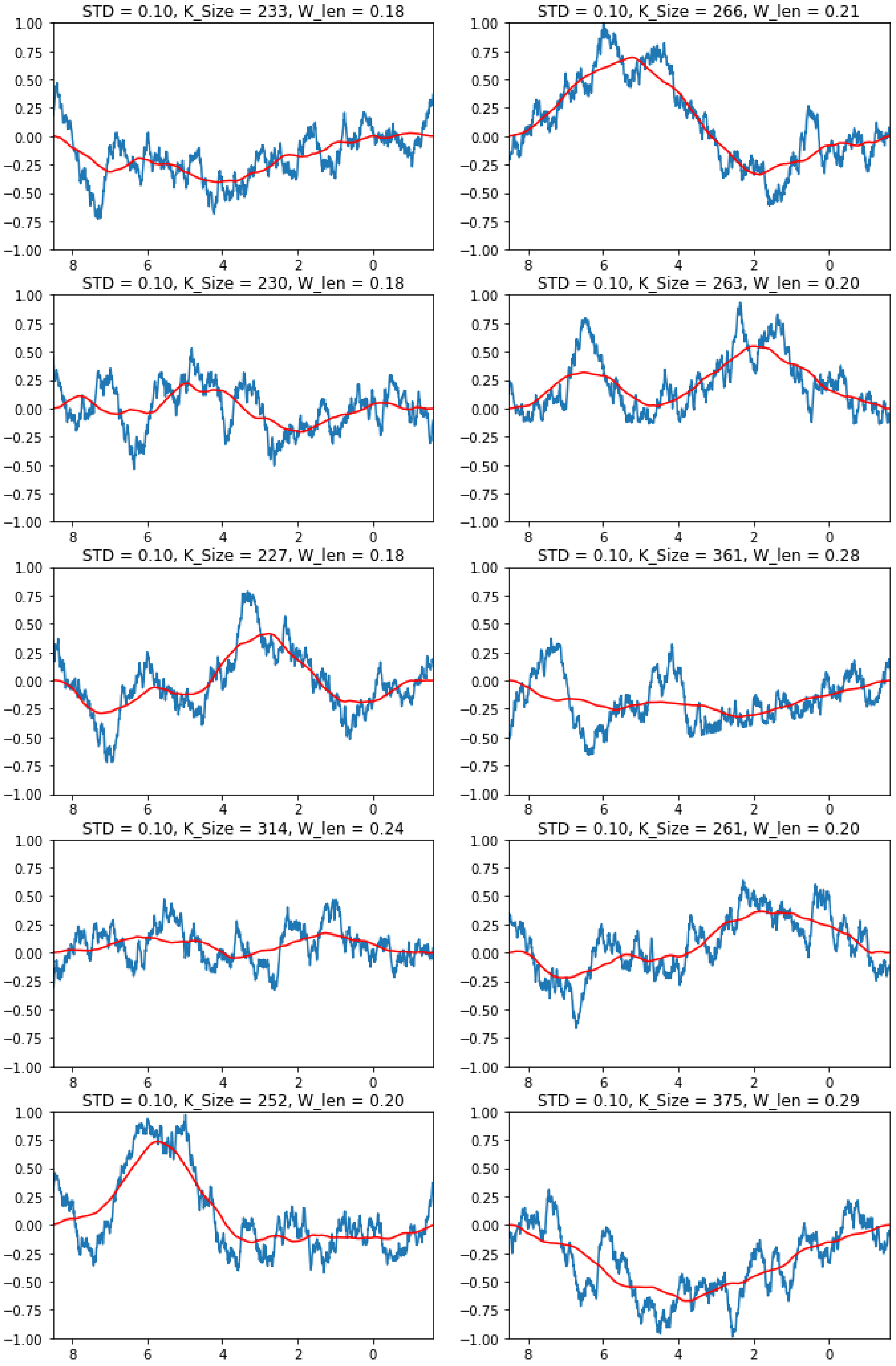

**FIGURE S4.3** Standard Deviation :: STD = 0.15; Window length = random; Kernel_size = random; Point density = 128

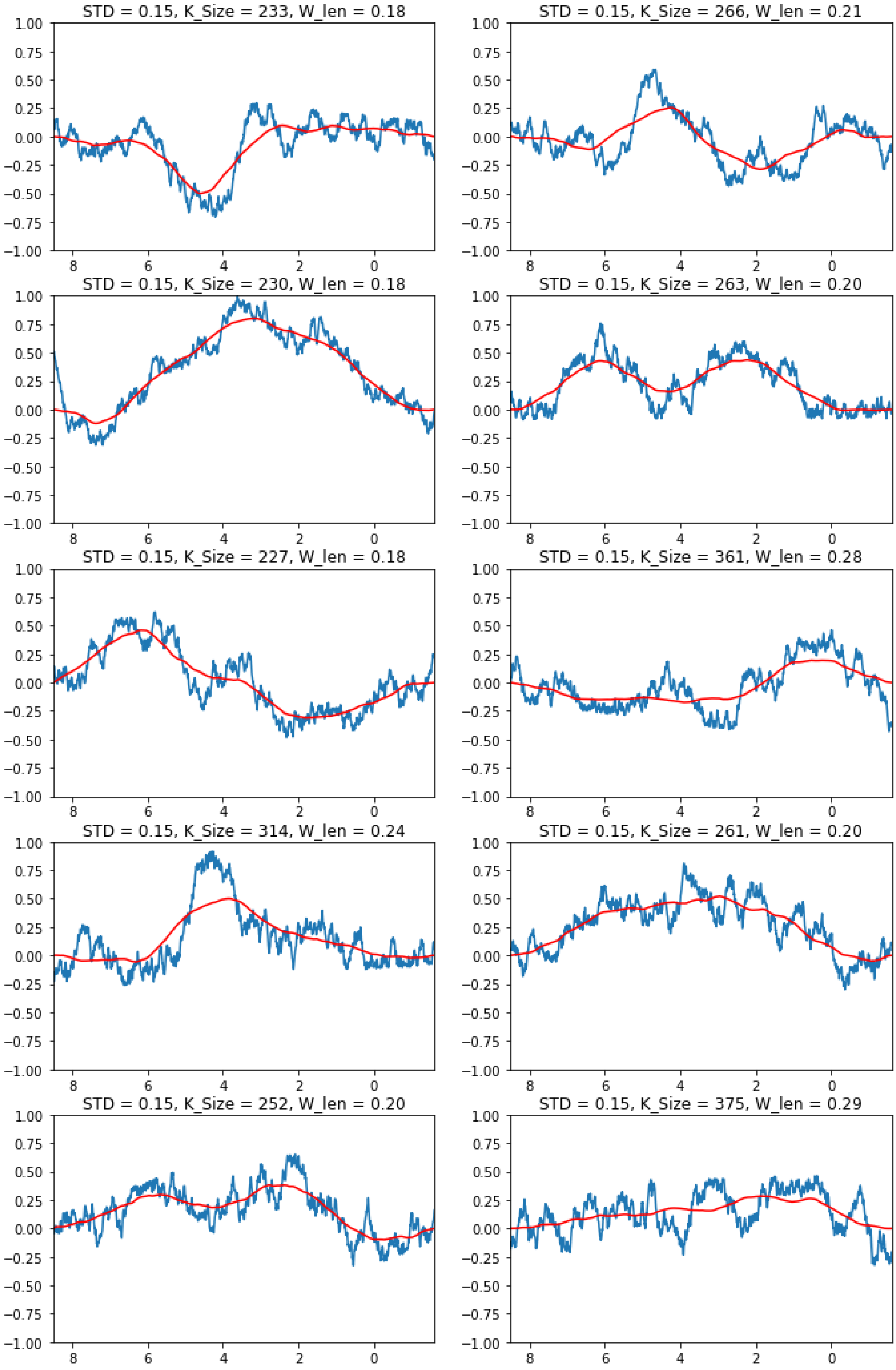

**FIGURE S4.4** Standard Deviation :: STD = 0.20; Window length = random; Kernel_size = random; Point density = 128

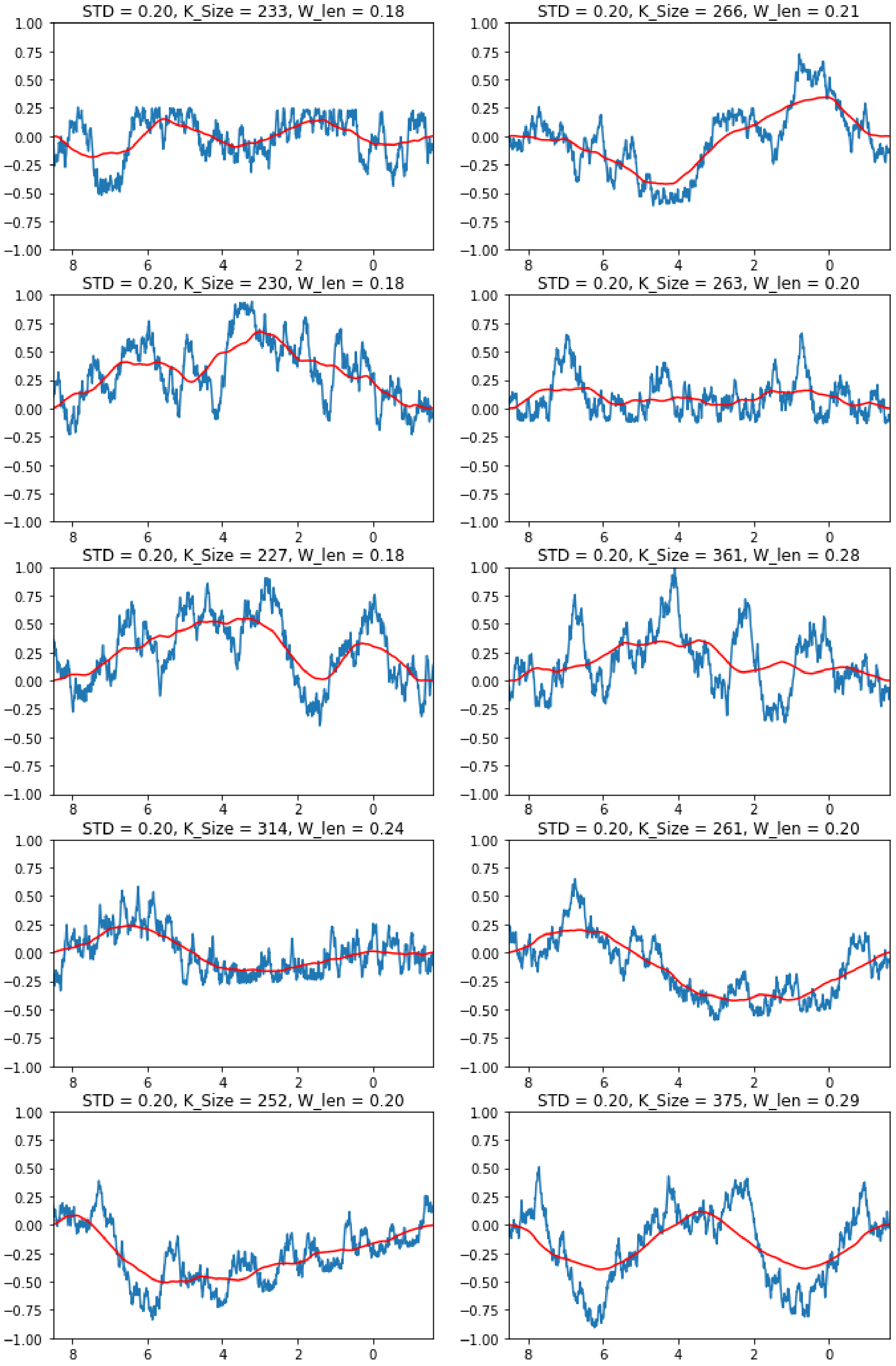

##### 4.5.2 | Smoothing Kernel

In this section, the effects of different smoothing kernel lengths are explored given a fixed *std* value.

**FIGURE S4.5** Smoothing Kernel :: STD = 0.10; Window Length = 0.10, Kernel_size = 60; Point density = 128

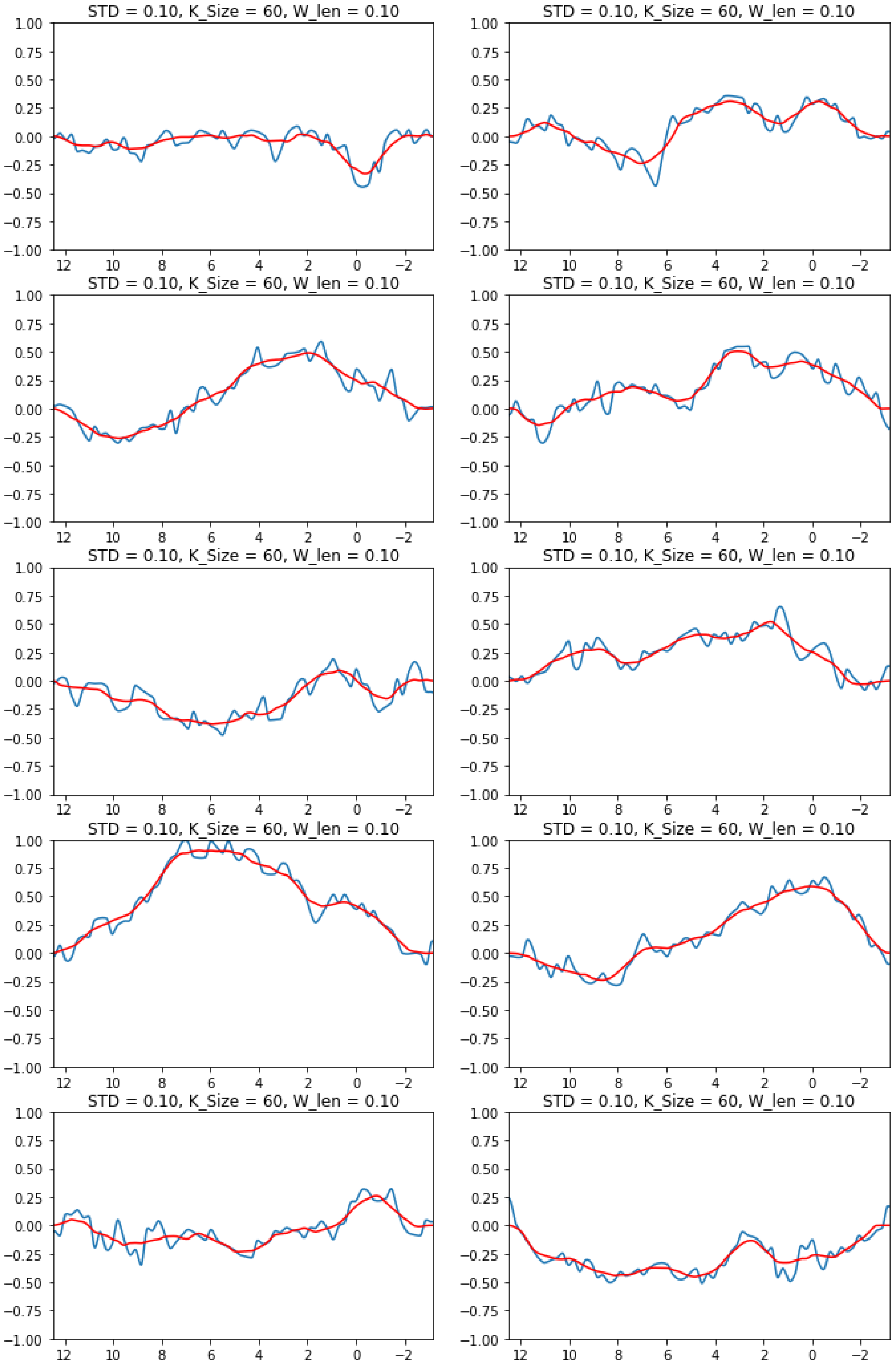

**FIGURE S4.6** Smoothing Kernel :: STD = 0.10; Window Length = 0.20, Kernel_size = 120; Point density = 128

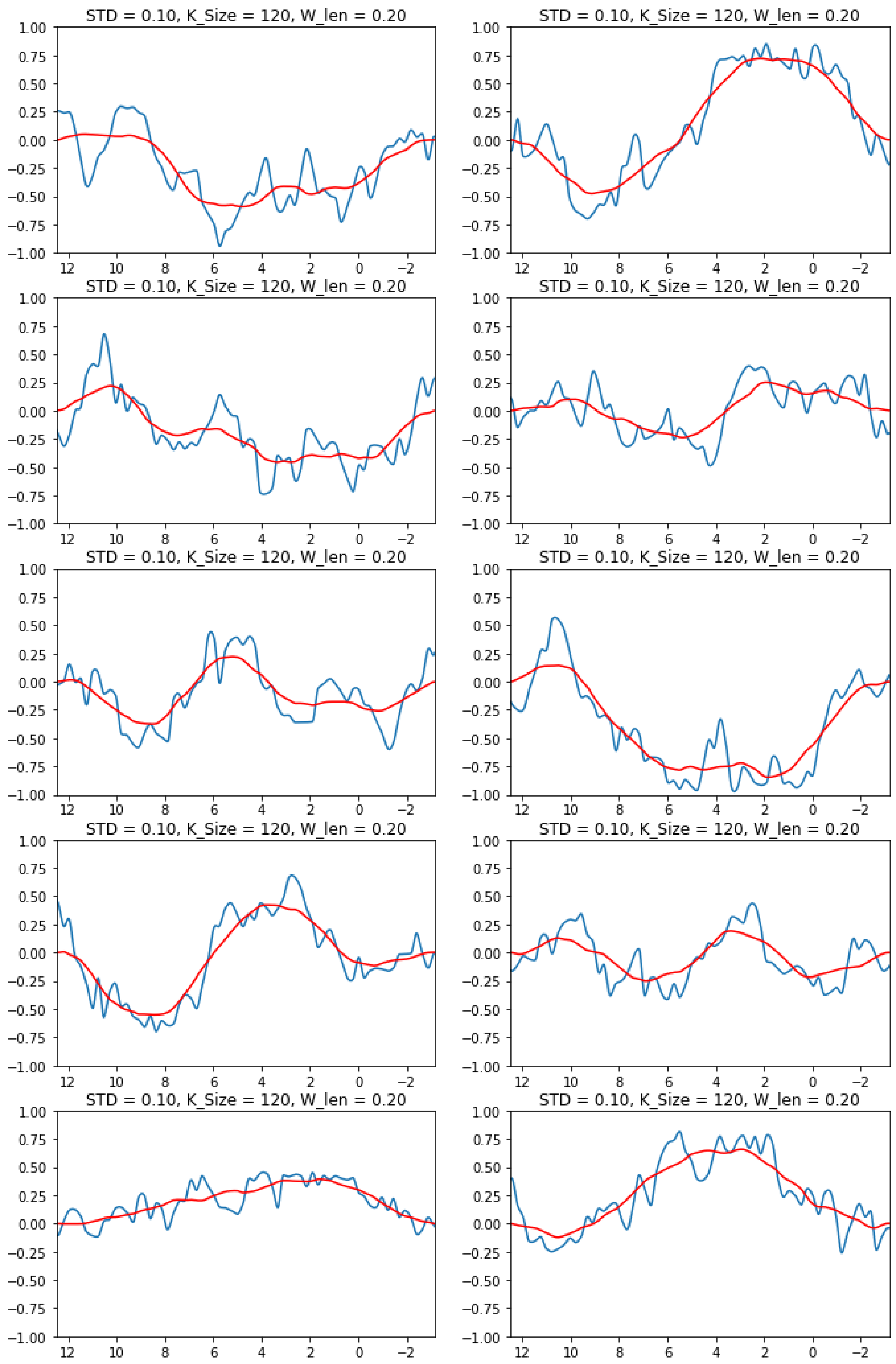

**FIGURE S4.7** Smoothing Kernel :: STD = 0.10; Window Length = 0.30, Kernel_size = 180; Point density = 128

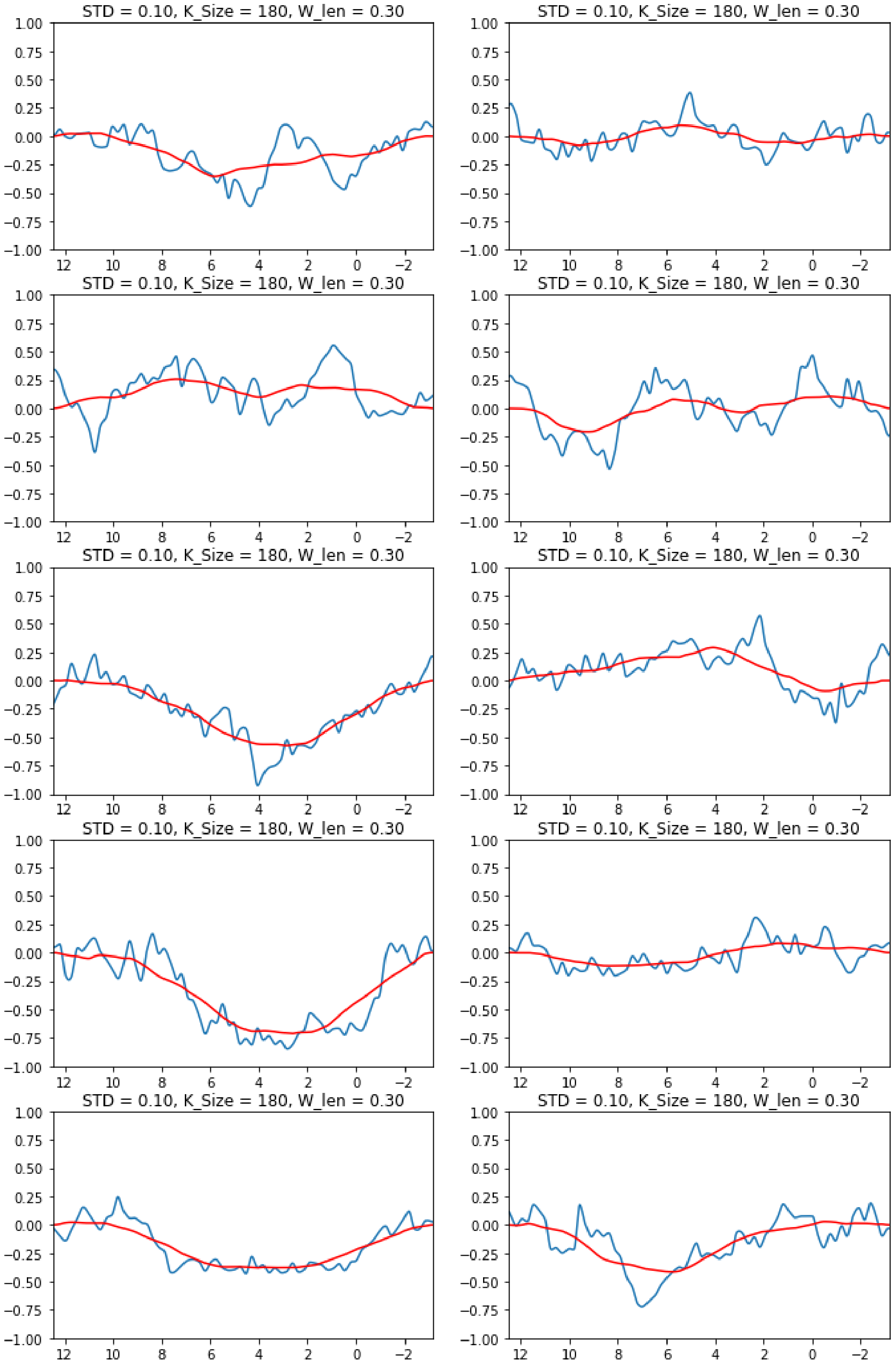

##### 4.5.3 | Point Density

This section explores the effect of varying the point density of the random walk while keeping the *std* fixed. Results are presented using two different kernel sizes for the smoothing.

**FIGURE S4.8** Point Density :: STD = 0.10, Window Length = [0.10, 0.30], Kernel_size = [3,9]; Point density = 64

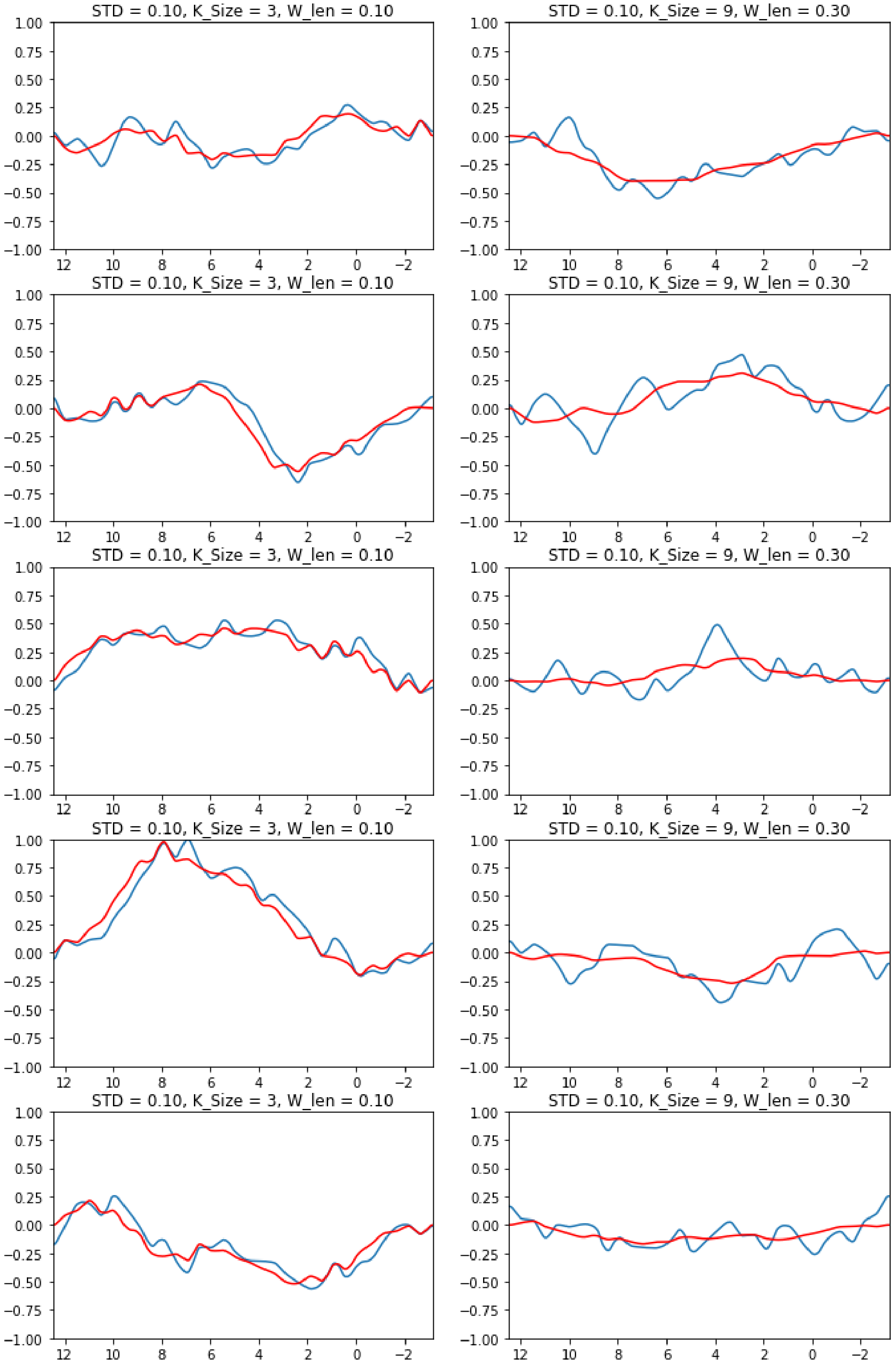

**FIGURE S4.9** Point Density :: STD = 0.10, Window Length = [0.10, 0.30], Kernel_size = [3,9]; Point density = 128

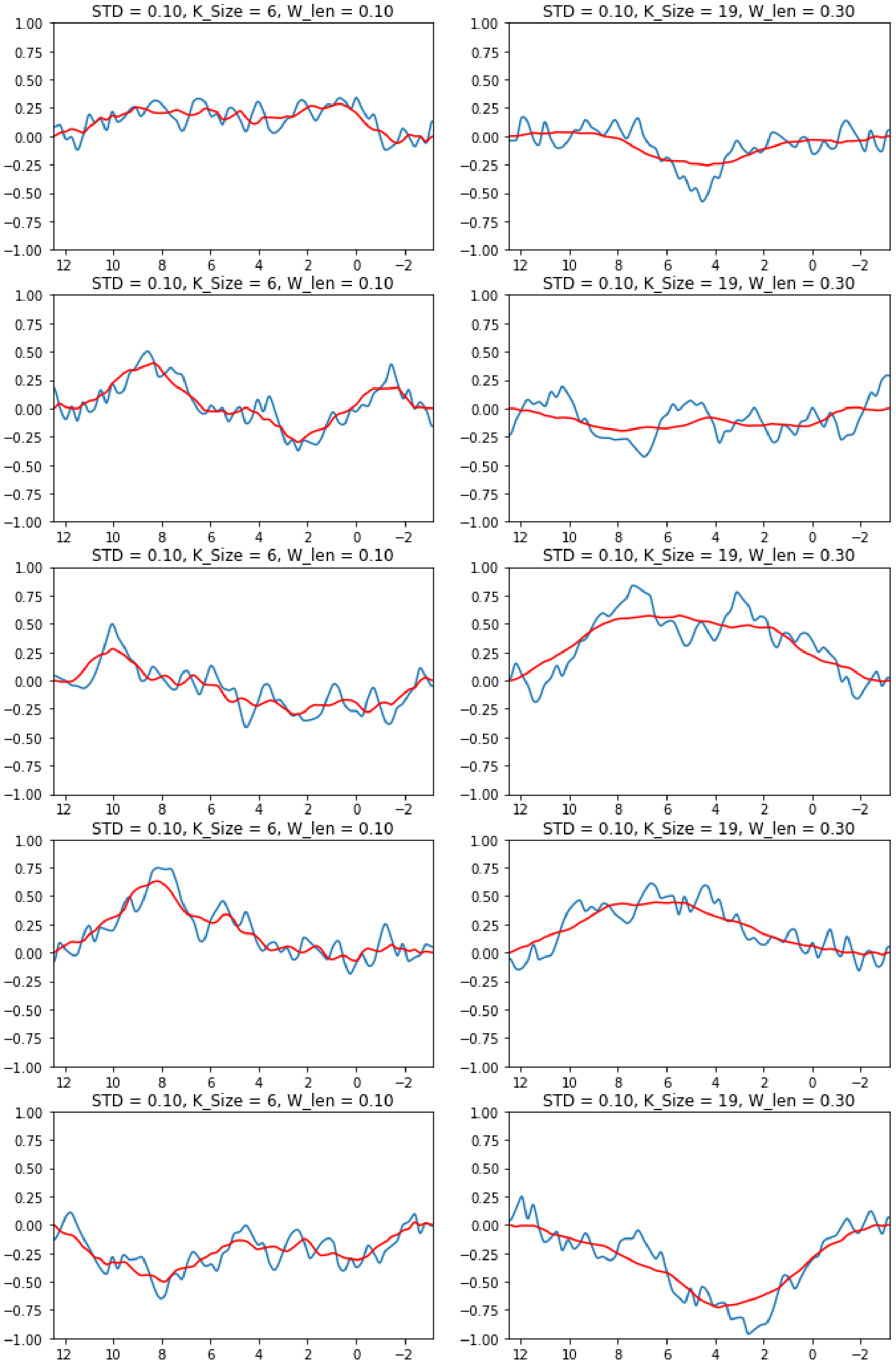

**FIGURE S4.10** Point Density :: STD = 0.10, Window Length = [0.10, 0.30], Kernel_size = [3,9]; Point density = 256

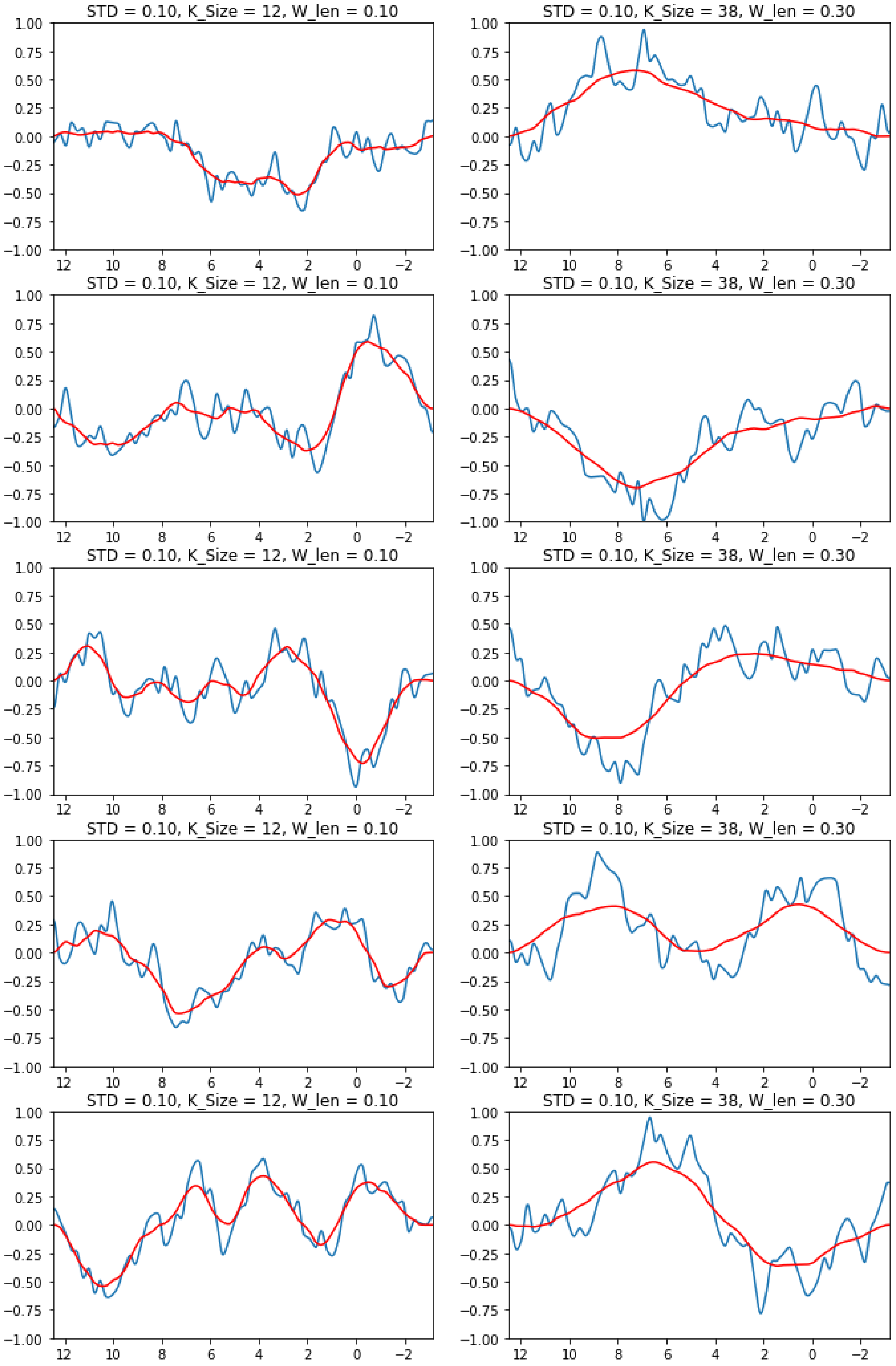

##### 4.5.4 | Incorporating the offsets into the spectra

To increase variability, the starting and ending heights are randomly selected. When considering the entire spectrum, however, there are two options to avoid unrealistic, hard transition points. These variables can be manually adjusted or the trend line can be removed, which is how the current code works.

**FIGURE S4.11** With the original trend lines still included

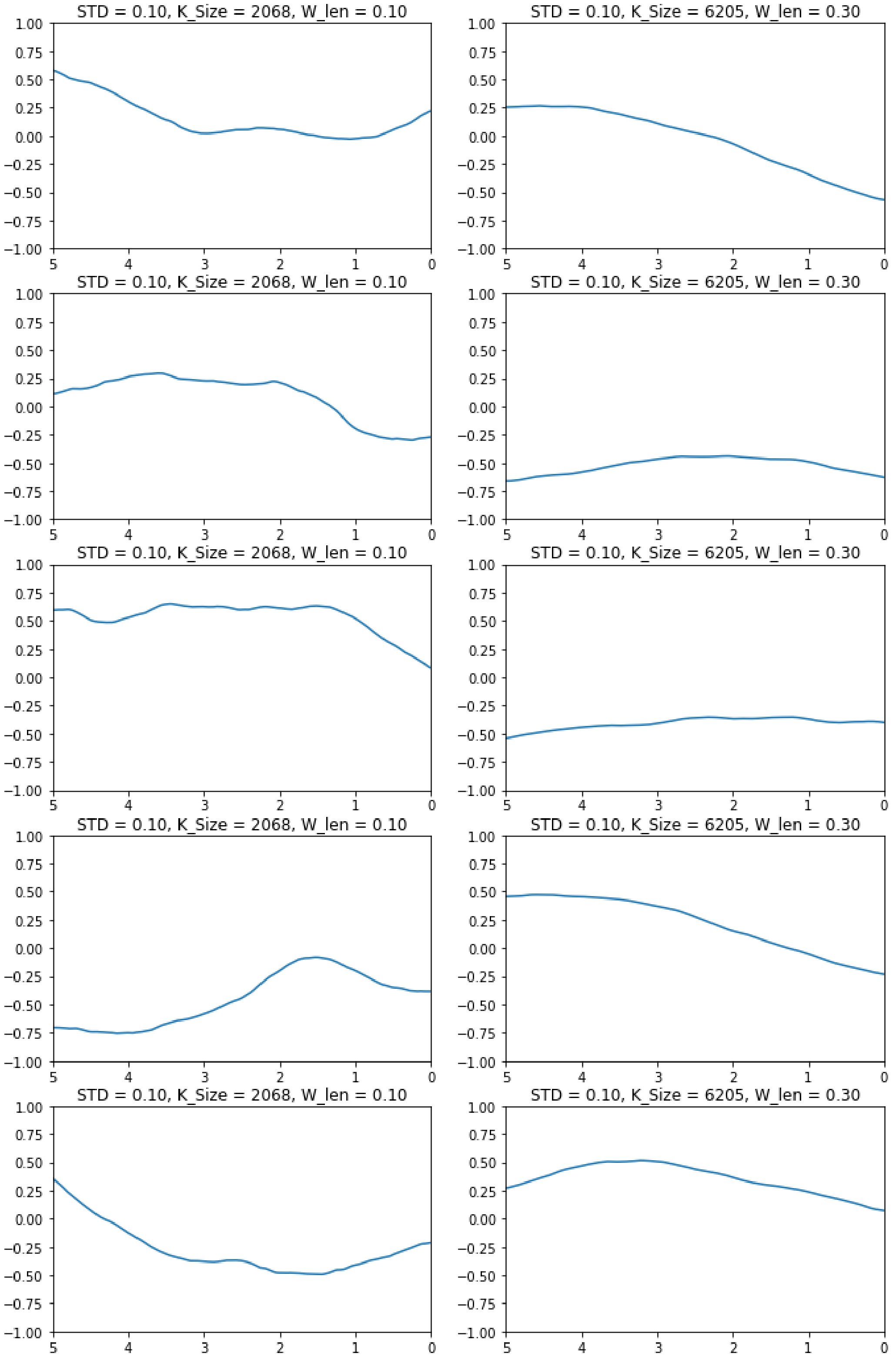

**FIGURE S4.12** With the original trend lines removed

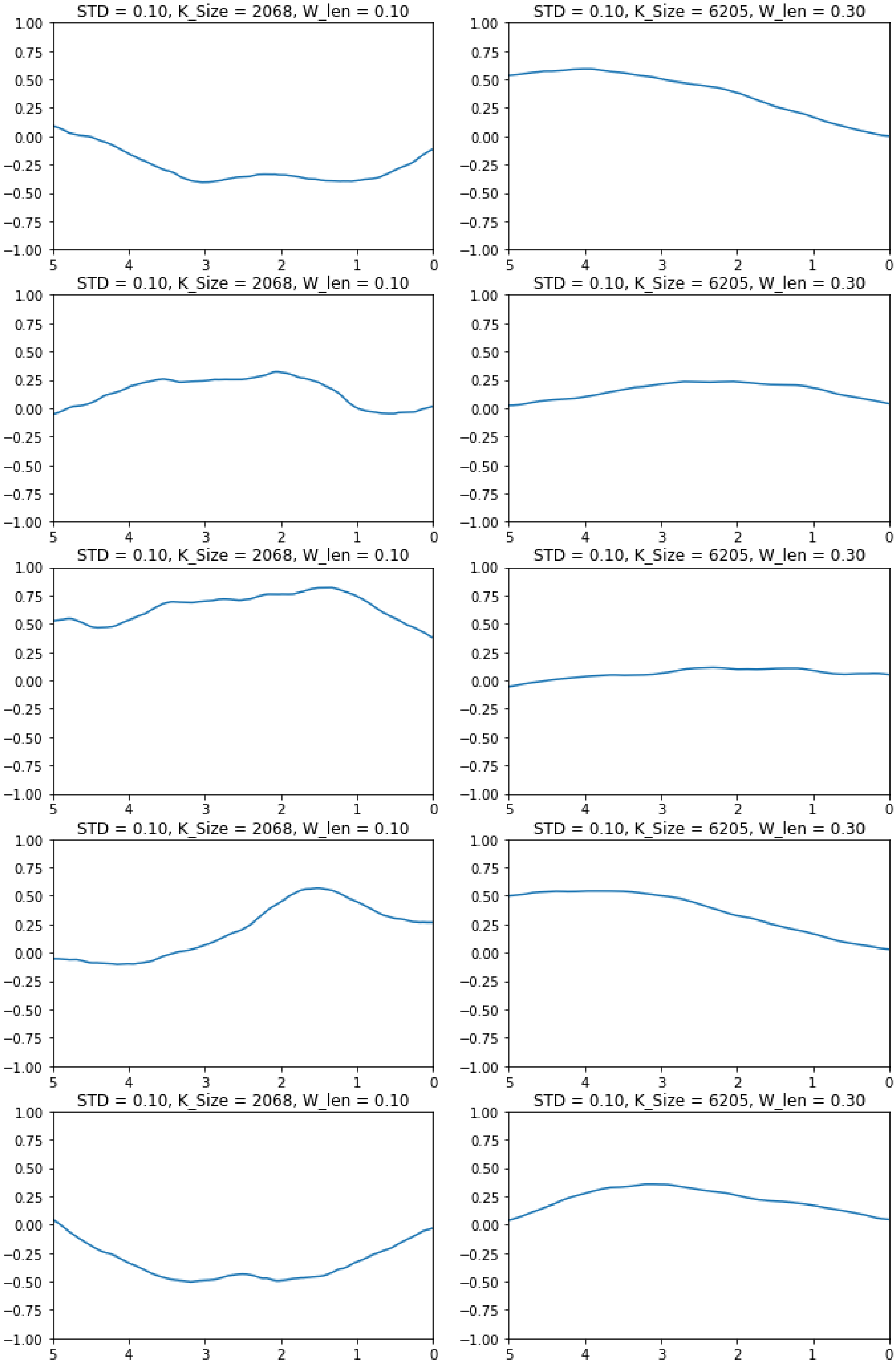

##### 4.5.5 | Effect of the Hilbert Transform

Simulating spectra requires complex spectral components, including the baseline and residual water. The Hilbert transform is used to generate those corresponding imaginary components.

**FIGURE S4.13** Compiled samples with their Hilbert pair

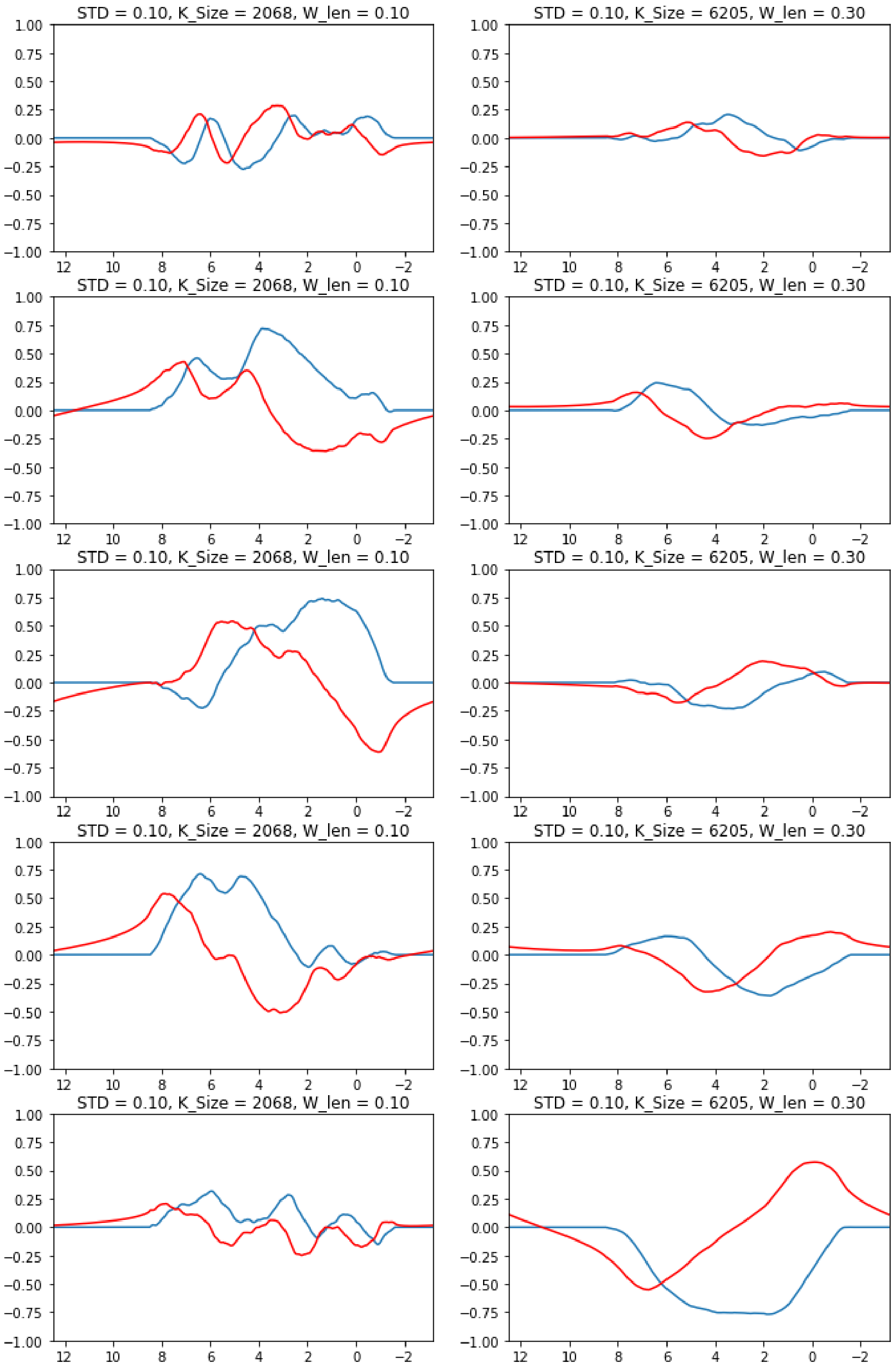

##### 4.5.6 | Compiled Generator

This shows the variety of baselines that can be generated when randomly sampling all variables.

**FIGURE S4.14** Compilation of randomly generated samples with their Hilbert pairs

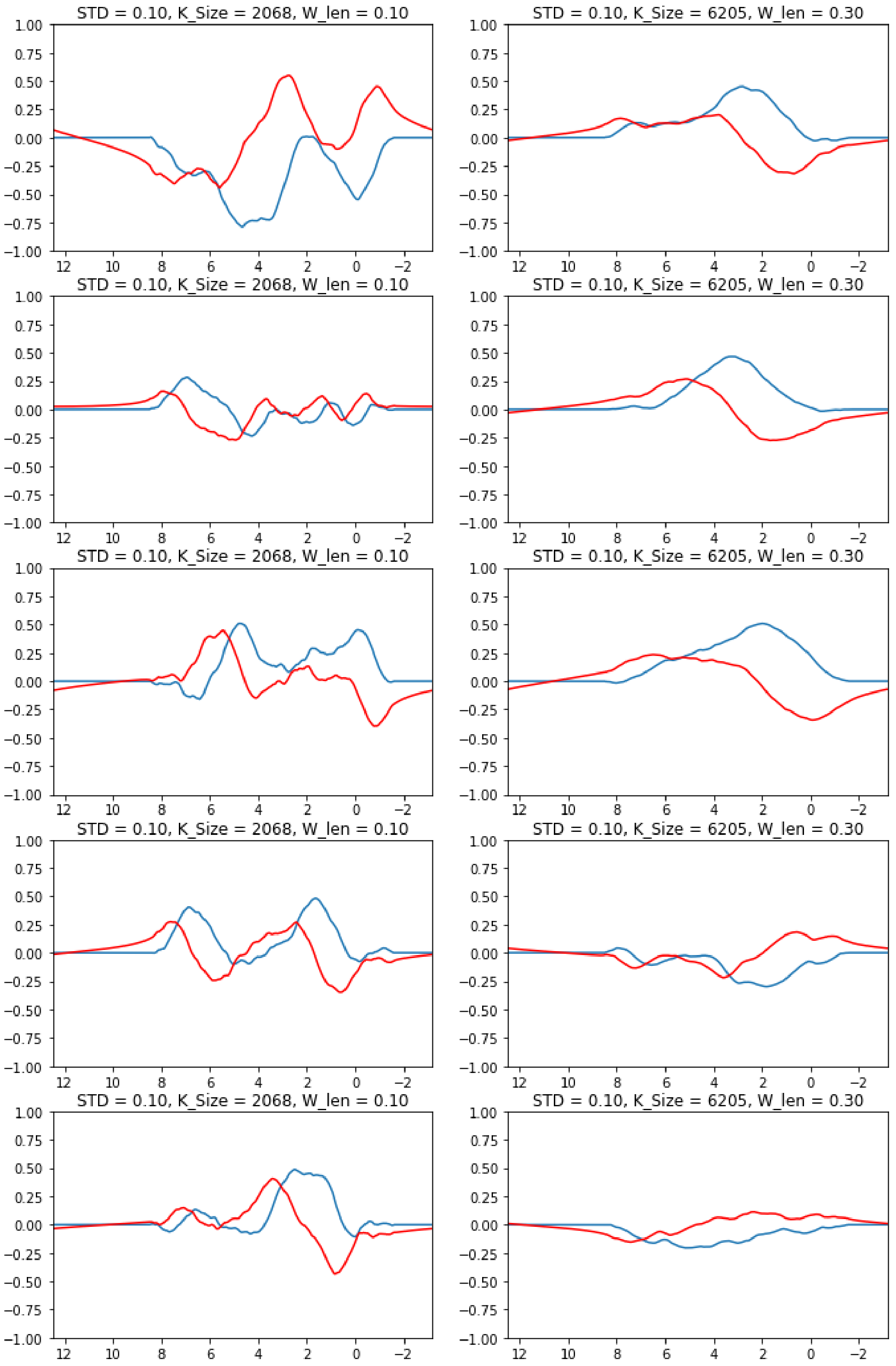

#### 4.6 | Residual Water

The following section will explore this generator using the residual water dictionary defined below. While the sections are the same as in Sec. 4.5, the values displayed are tailored to the ranges for the residual water simulations.

##### 4.6.1 | Configuration dictionaries

The following dictionary entries are the standard defaults in MRS-Sim for generating residual water contributions.

```
resWater_cfg = {
        "start":           [          0],
        "end":             [          0],
        "upper":           [    0,    1],
        "lower":           [    0,    1],
        "std":             [  0.2, 0.40],
        "window":          [ 0.05, 0.15],
        "pt_density":        1204,
        "ppm_range":       [  4.4,  4.9],
        "prime":             0.15,
        "scale":           [  1.0,  1.0], # typically: [0.05, 0.20] - better visualization.
        "drop_prob":          0.0
}
```

##### 4.6.2 | Standard Deviation

This section shows the effect of different *std* values on the random walk. The *std* variable is used when sampling the noise for the walk. It directly controls the amount of variation between two consecutive points prior to the cumulative summation.

**FIGURE S4.15** Standard Deviation :: STD = 0.20; Window length = random; Kernel_size = random; Point density = 1204

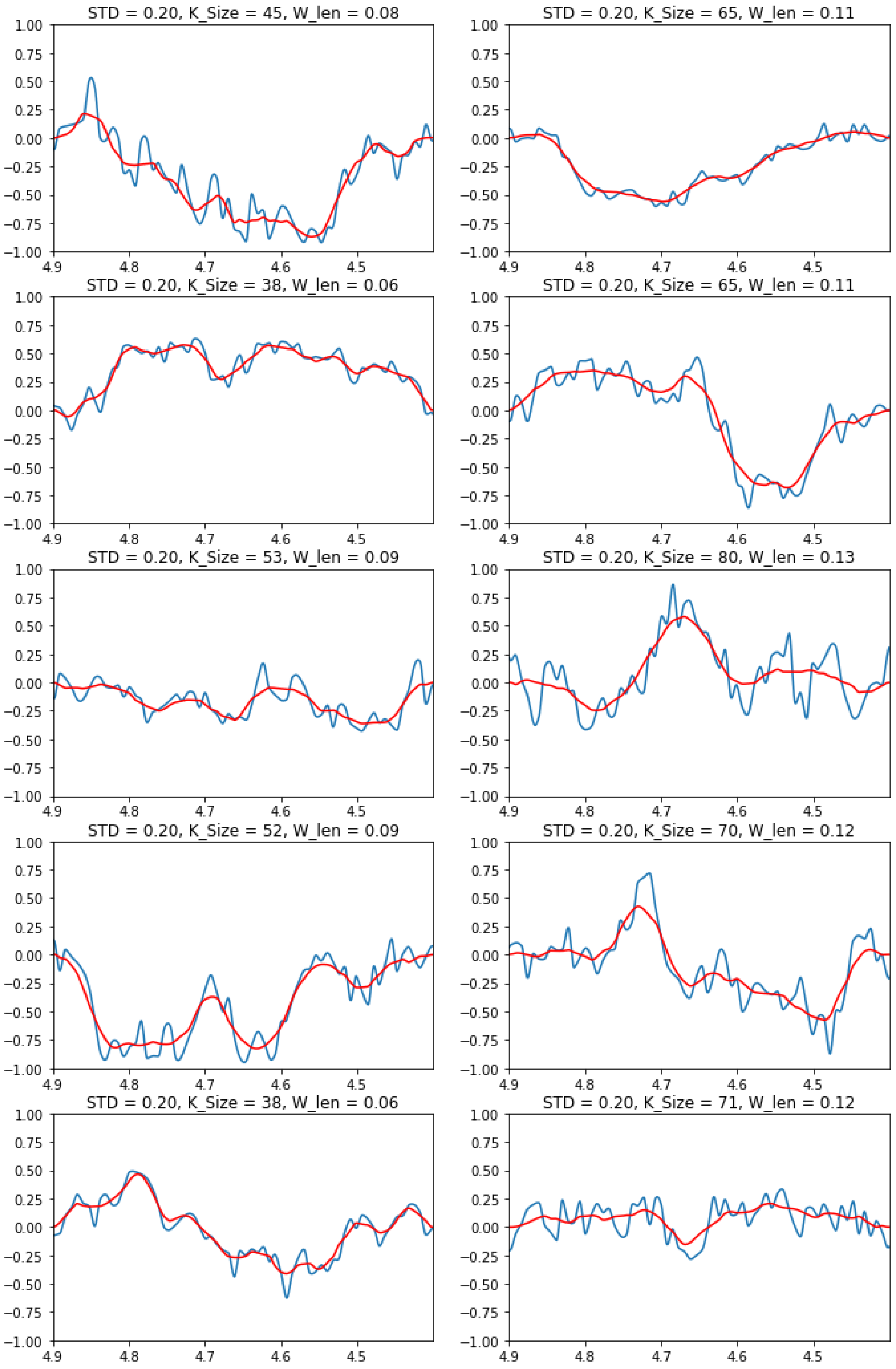

**FIGURE S4.16** Standard Deviation :: STD = 0.30; Window length = random; Kernel_size = random; Point density = 1204

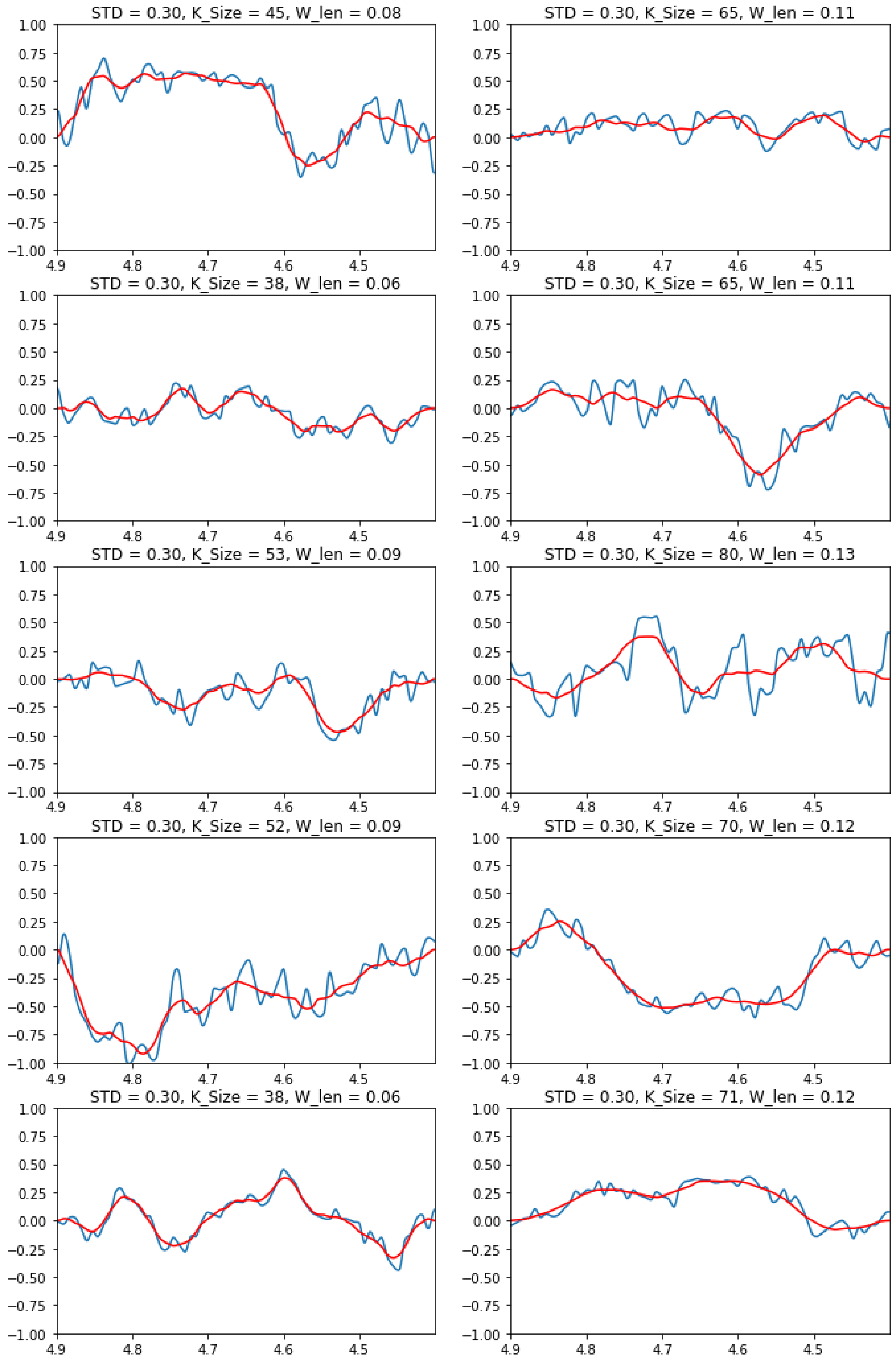

**FIGURE S4.17** Standard Deviation :: STD = 0.40; Window length = random; Kernel_size = random; Point density = 1204

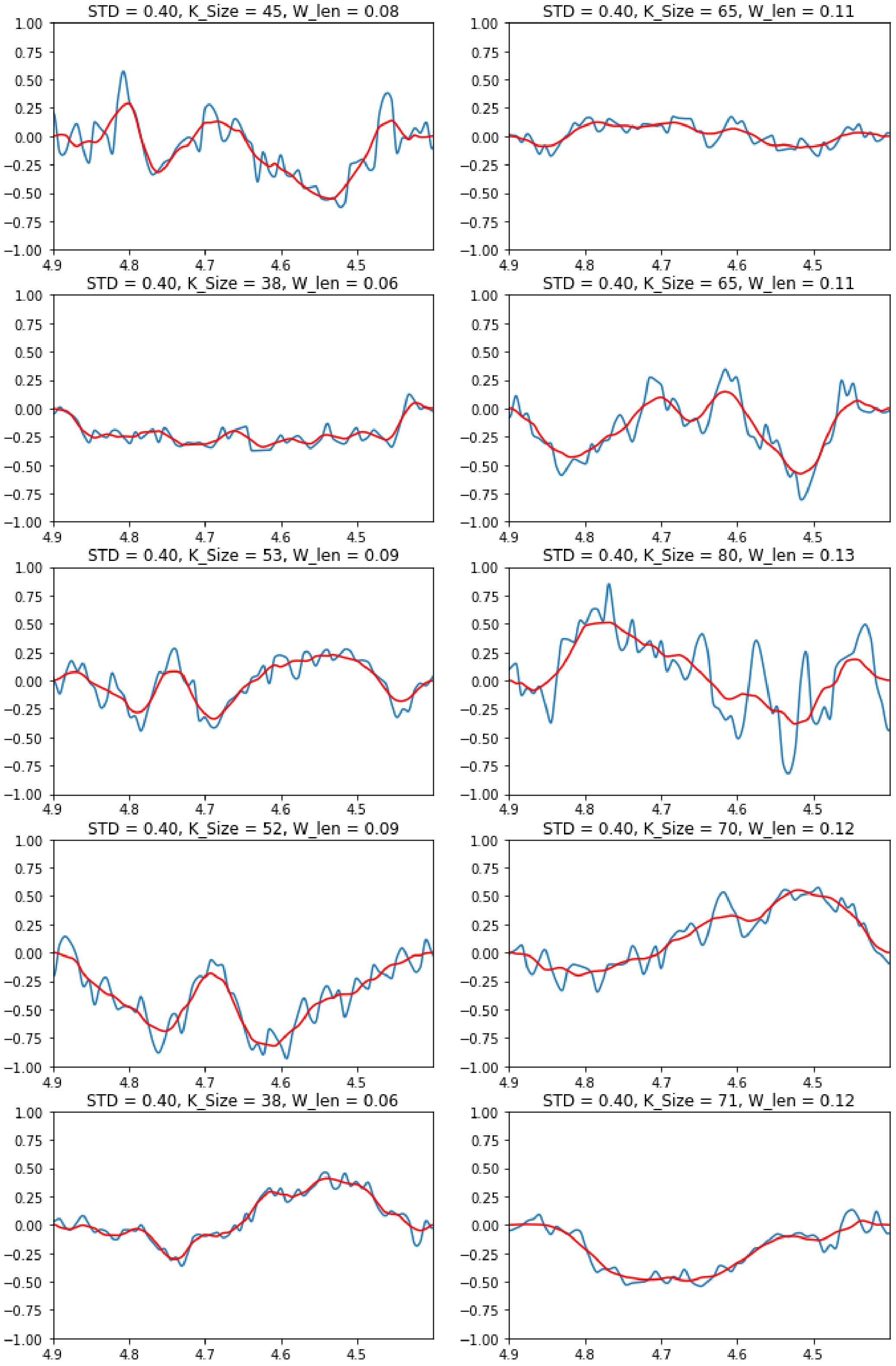

**FIGURE S4.18** Standard Deviation :: STD = 0.50; Window length = random; Kernel_size = random; Point density = 1204

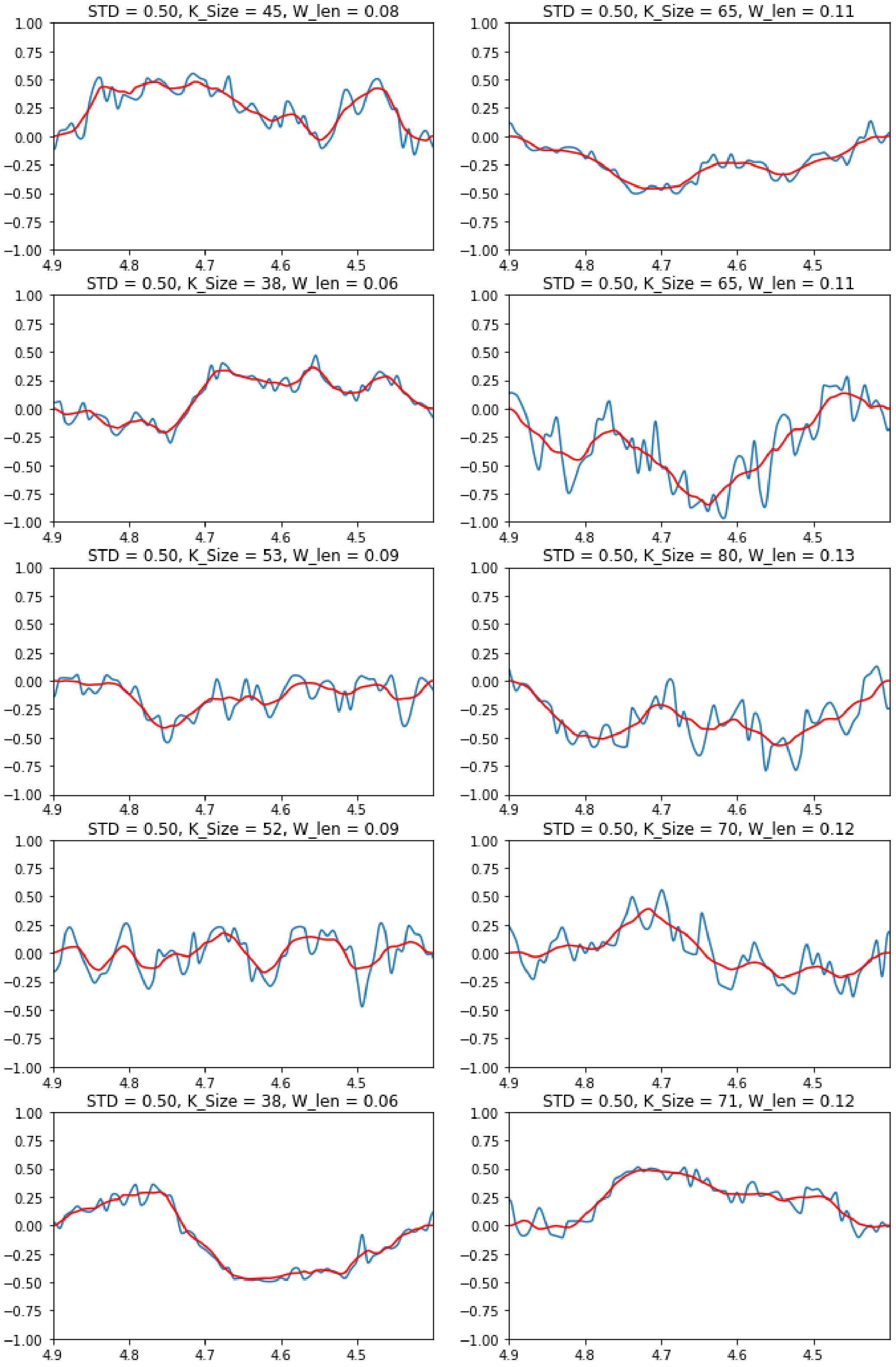

##### 4.6.3 | Smoothing Kernel

In this section, the effects of different smoothing kernel lengths are explored given a fixed *std* value.

**FIGURE S4.19** Smoothing Kernel :: STD = 0.10, Window Length = 0.05, Kernel_size = 30; Point density = 1204

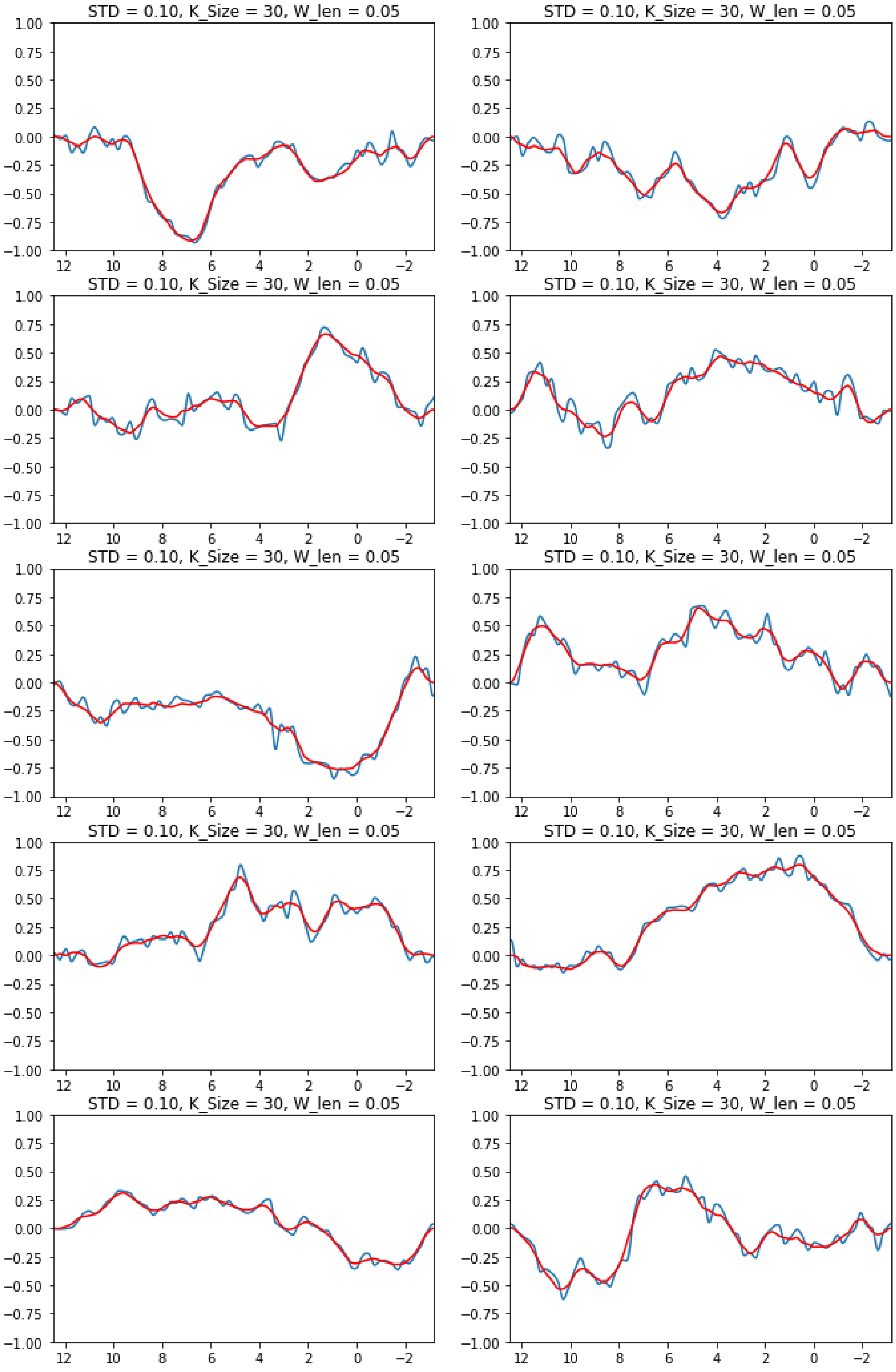

**FIGURE S4.20** Smoothing Kernel :: STD = 0.10, Window Length = 0.10, Kernel_size = 60; Point density = 1204

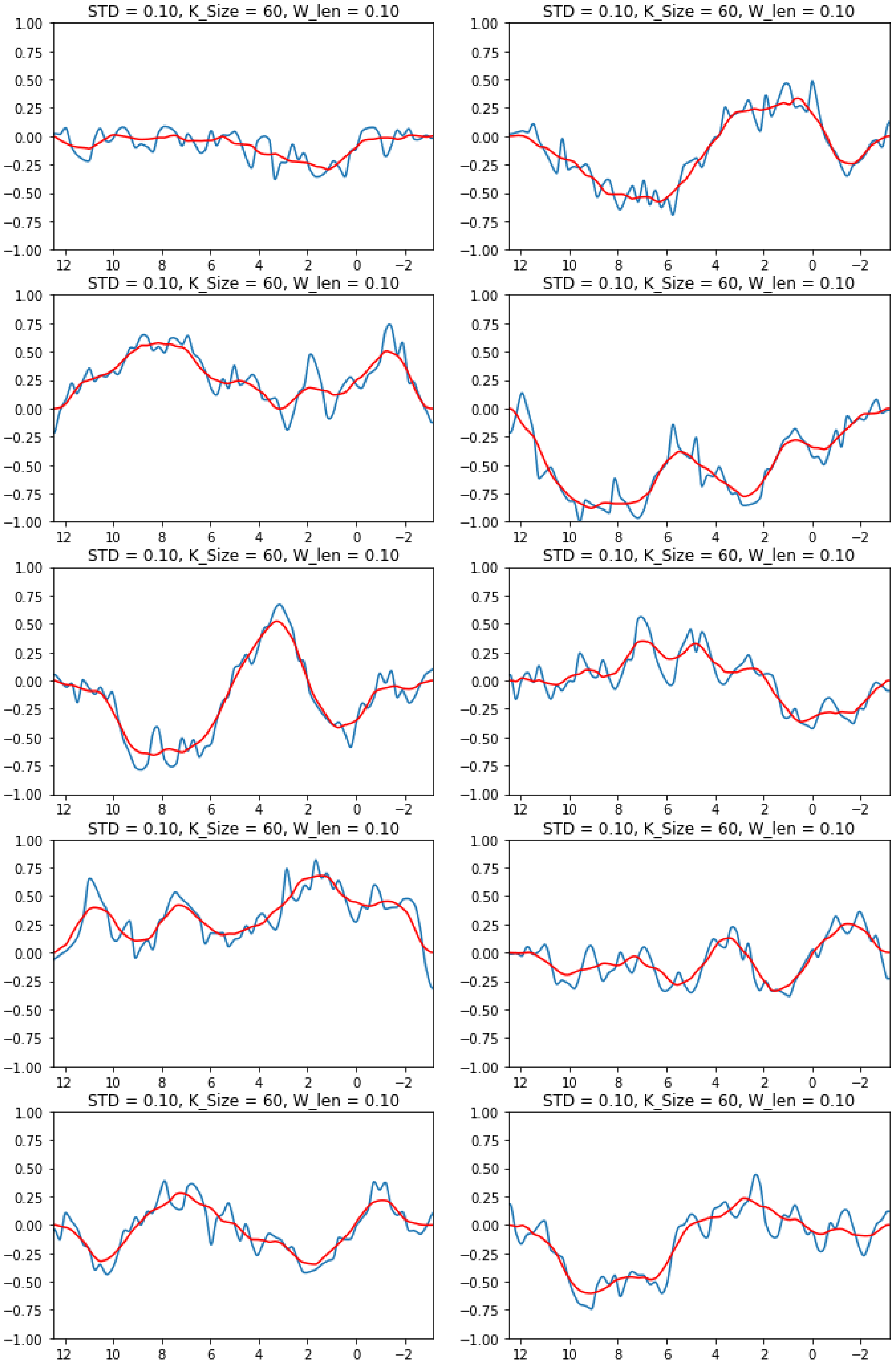

**FIGURE S4.21** Smoothing Kernel :: STD = 0.10, Window Length = 0.15, Kernel_size = 90; Point density = 1204

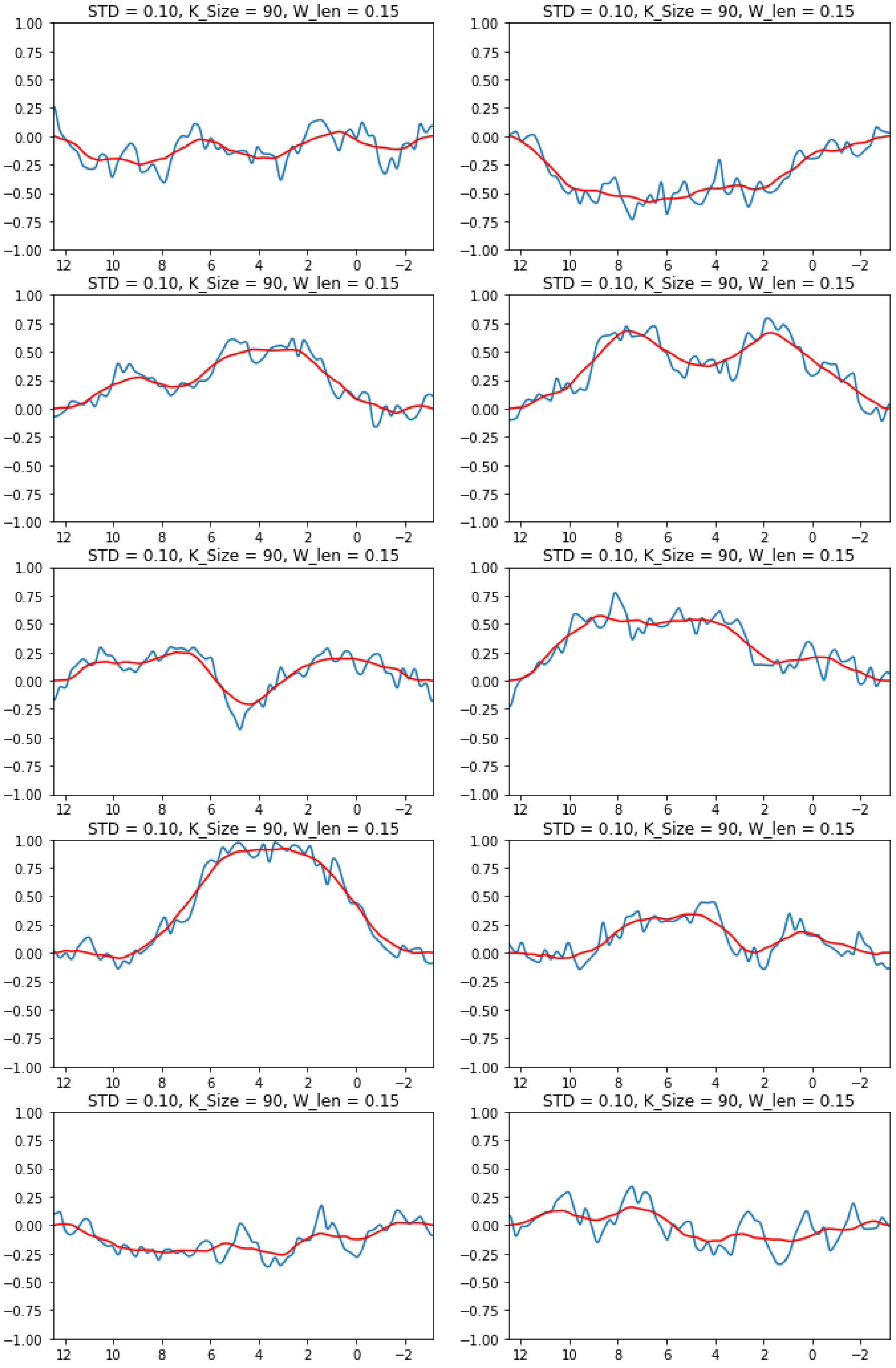

##### 4.6.4 | Point Density

This section explores the effect of varying the point density of the random walk while keeping the *std* fixed. Results are presented using two different kernel sizes for the smoothing.

**FIGURE S4.22** Point Density :: STD = 0.10, Window Length = [0.05, 0.15], Kernel_size = [12,38]; Point density = 512

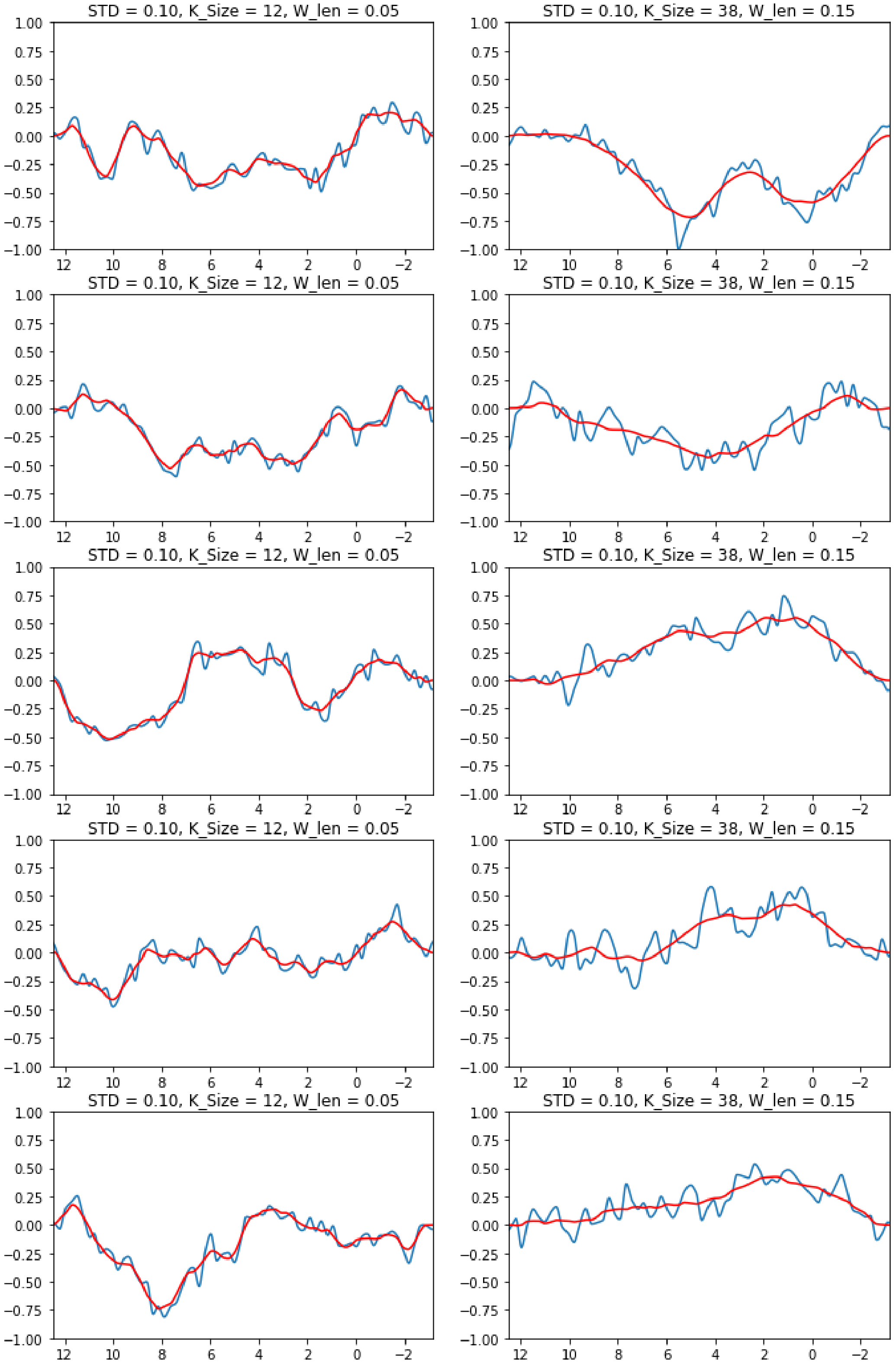

**FIGURE S4.23** Point Density :: STD = 0.10, Window Length = [0.05, 0.15], Kernel_size = [12,38]; Point density = 1024

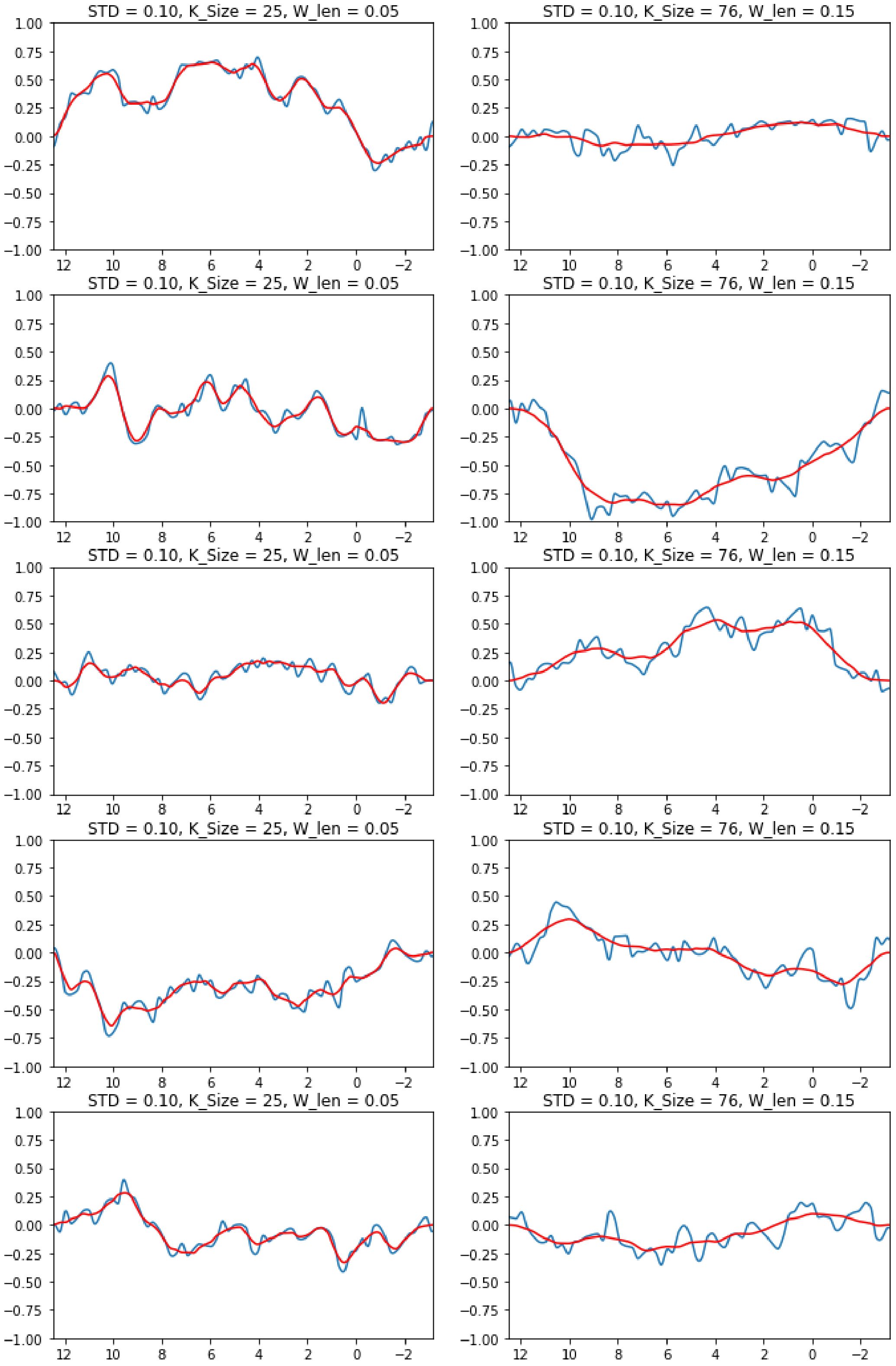

**FIGURE S4.24** Point Density :: STD = 0.10, Window Length = [0.05, 0.15], Kernel_size = [12,38]; Point density = 2048

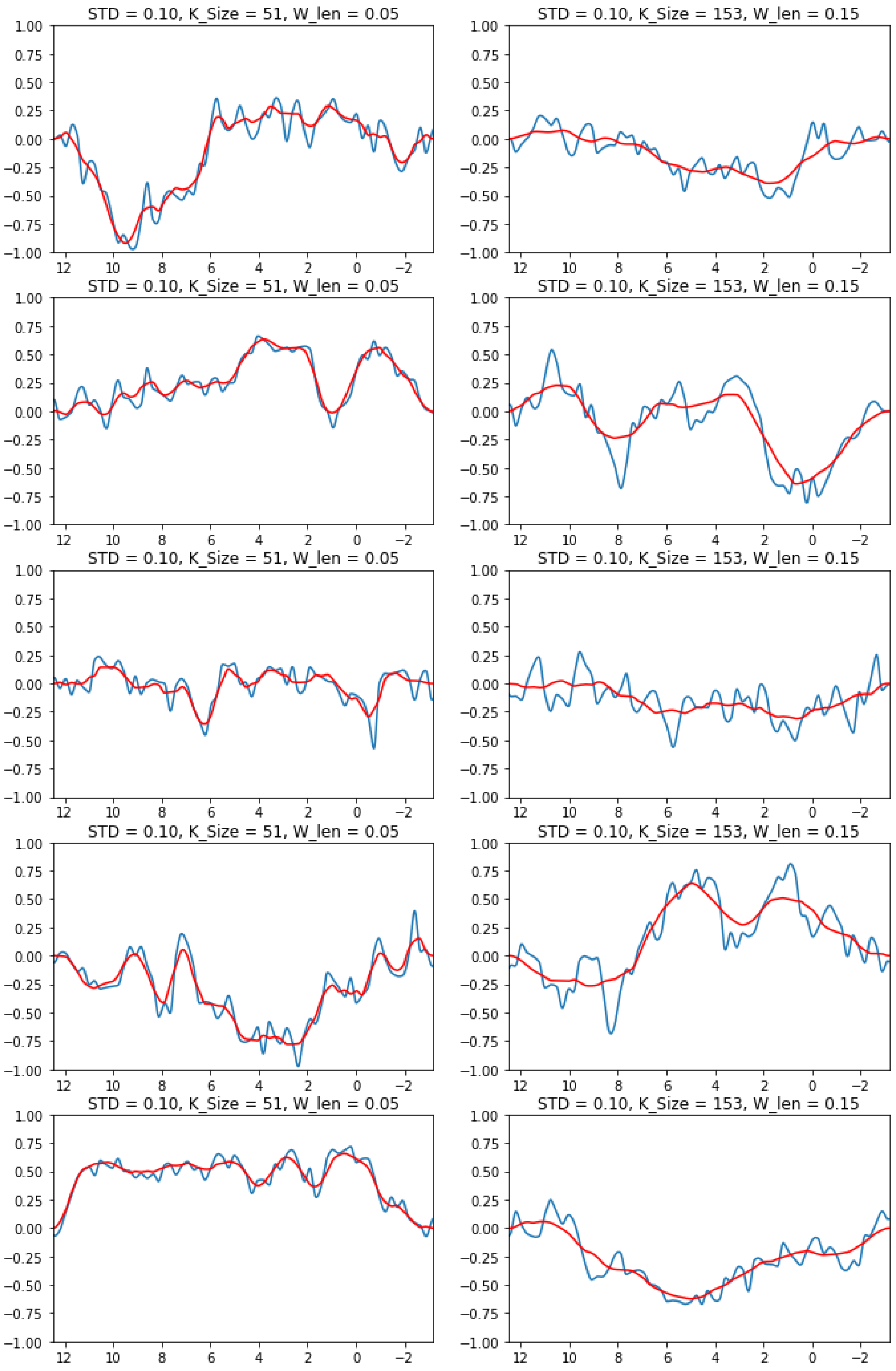

##### 4.6.5 | Incorporating the offsets into the spectra

To increase variability, the starting and ending heights are randomly selected. When considering the entire spectrum, however, these must be adjusted to avoid unrealistic, hard transition points. As mentioned above, this is achieved by removing the trend lines.

**FIGURE S4.25** With the original trend lines still included

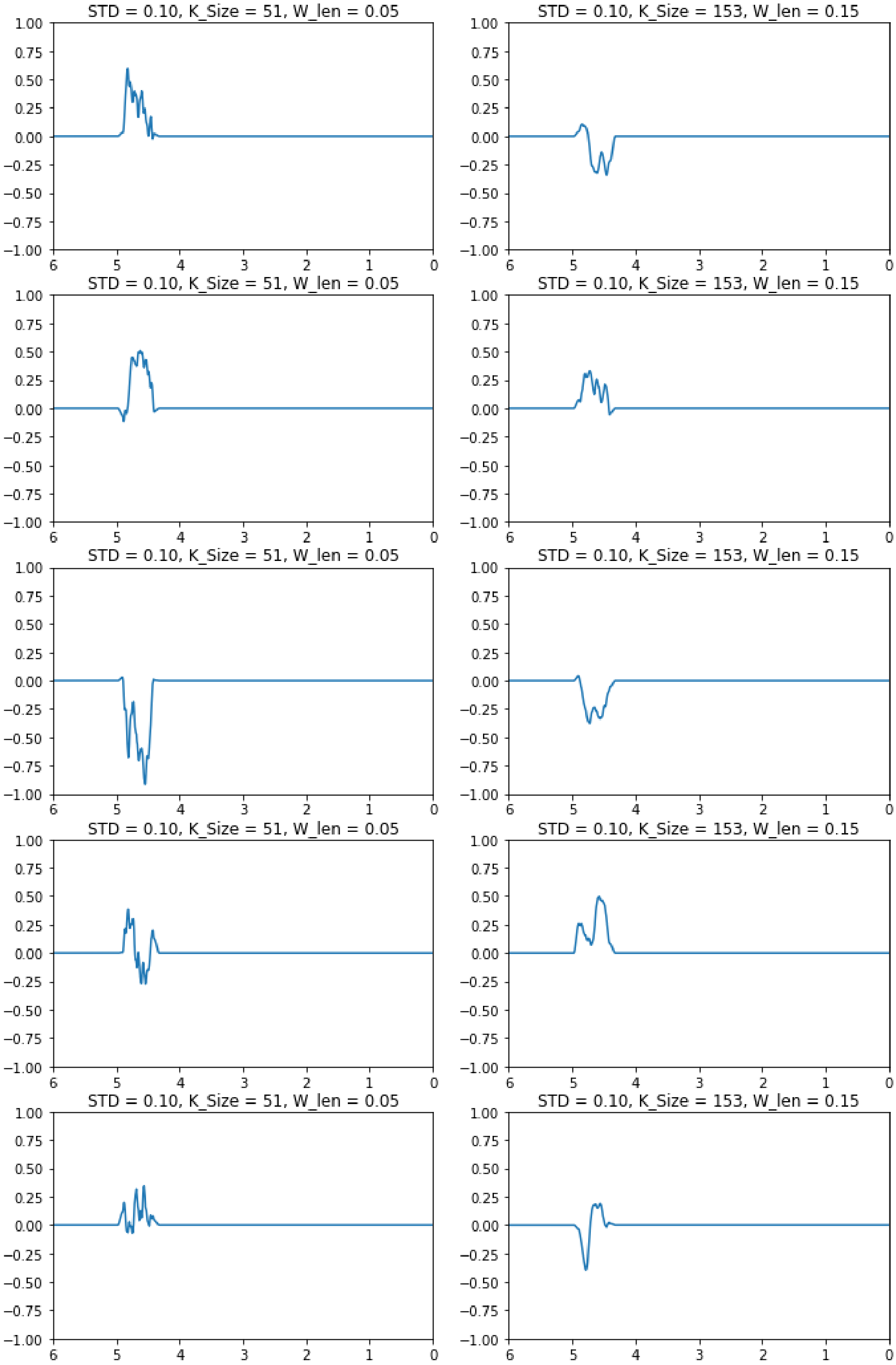

**FIGURE S4.26** With the original trend lines removed

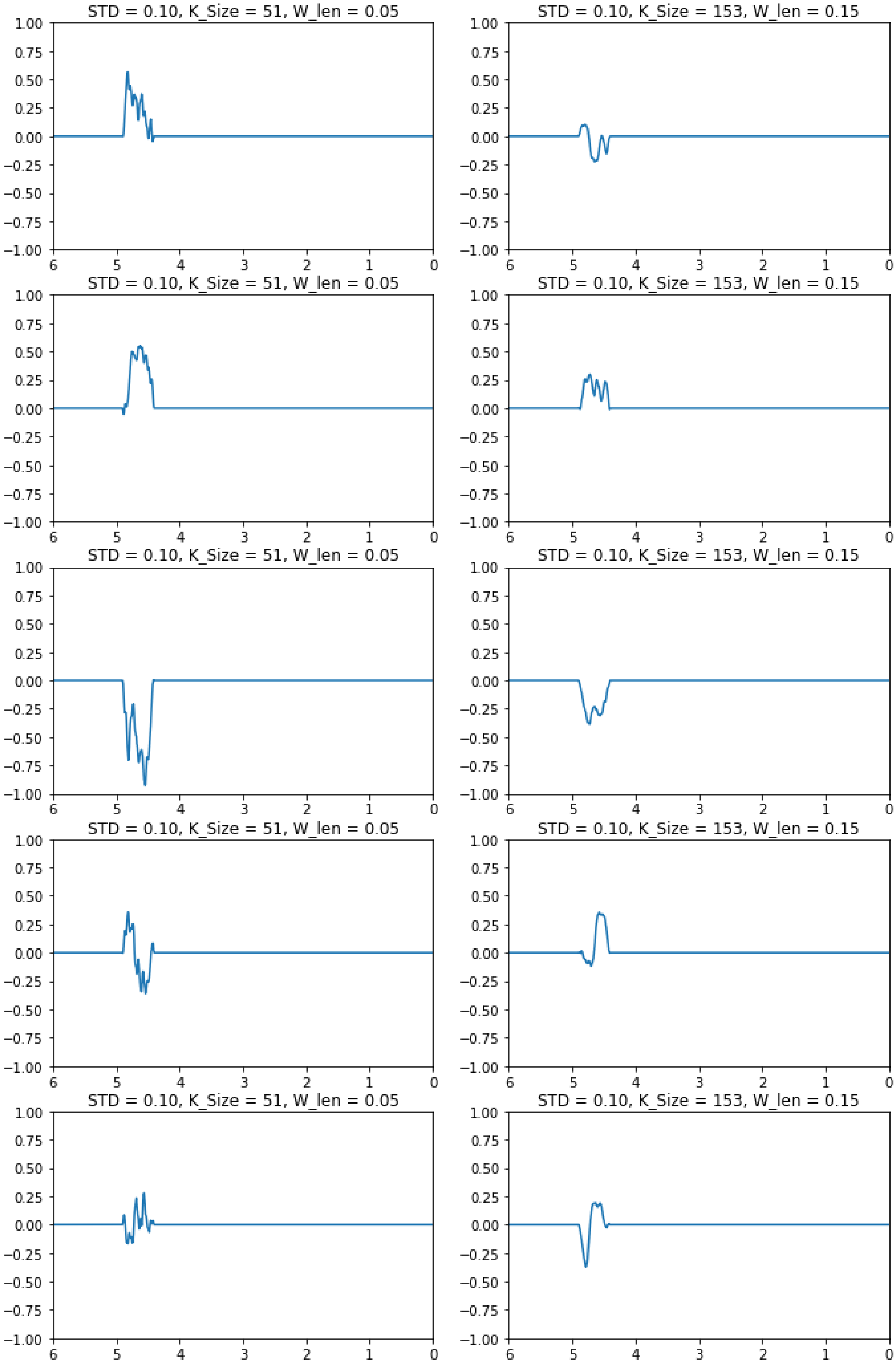

##### 4.6.6 | Effect of the Hilbert Transform

Simulating spectra requires complex spectral components, including the baseline and residual water. The Hilbert transform is used to generate those corresponding imaginary components.

**FIGURE S4.27** Compiled samples with their Hilbert pair

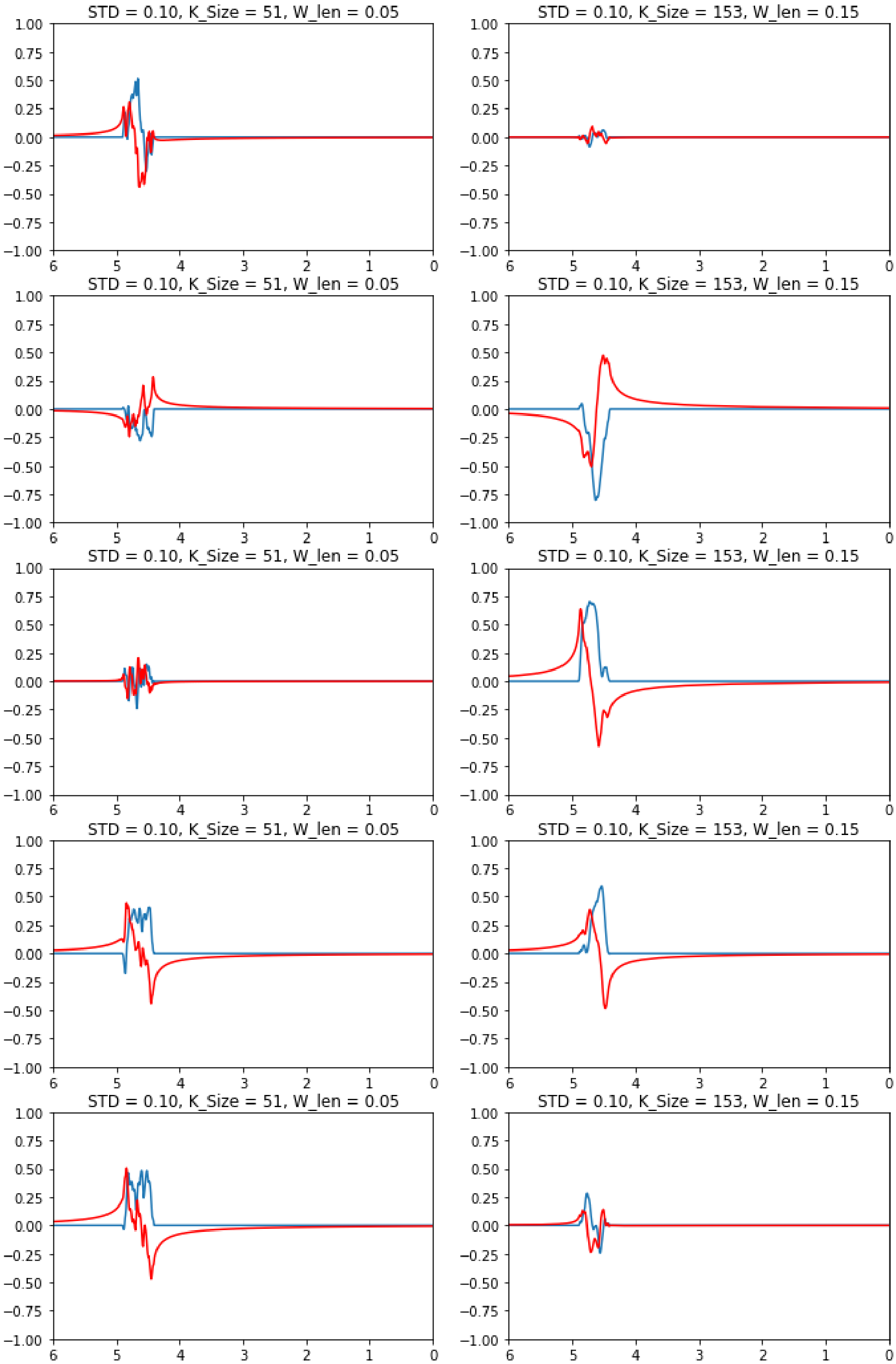

##### 4.6.7 | Compiled Generator

This shows the variety of residual water contributions that can be generated when randomly sampling all variables.

**FIGURE S4.28** Compilation of randomly generated samples with their Hilbert pairs

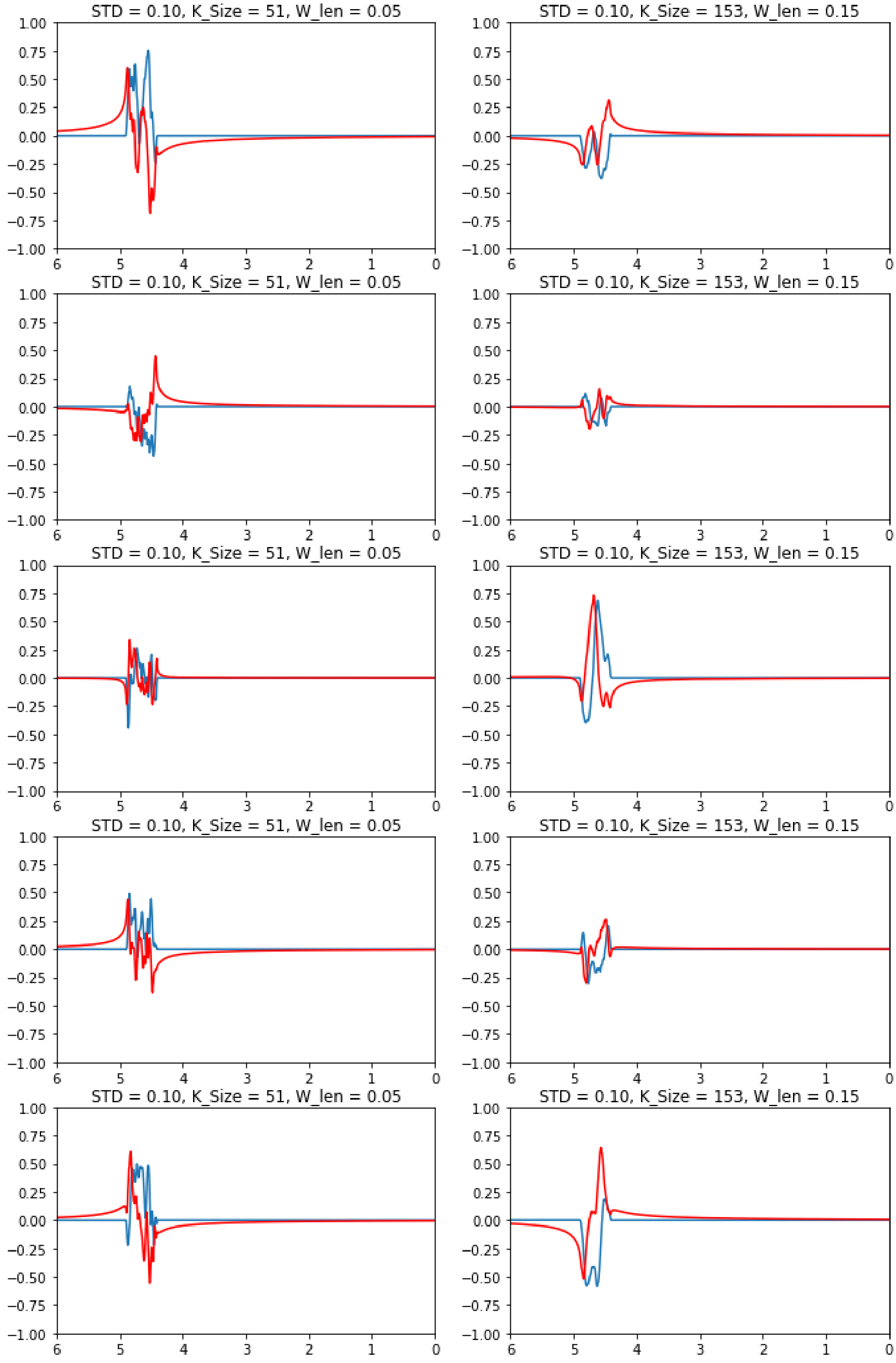
